# OntoContext, a new python package for gene contextualization based on the annotation of biomedical texts

**DOI:** 10.1101/2022.05.27.493696

**Authors:** Walid Bedhiafi, Véronique Thomas-Vaslin, Amel Benammar Elgaaied, Adrien Six

**Affiliations:** Université de Tunis El Manar, Faculté des Sciences de Tunis, LR05ES05 Laboratoire de Génétique, Immunologie et Pathologies Humaines, 2092, Tunis, Tunisie; Sorbonne Universités, UPMC Univ Paris 06, INSERM, UMRS959, Immunology-Immunopathology-Immunotherapy (I3), F-75013 Paris, France

## Abstract

**Motivation:** The automatic mining for bibliography exploitation in given contexts is a challenge according to the increasing number of scientific publications and new concepts. Several indexing systems were developed for biomedical literature. However, such systems have failed to produce contextualised research of genes and proteins and automatically group texts according to shared concepts. In this paper, we present OntoContext, a contextualization system crossing the use of biomedical ontologies to annotate texts containing terms related to cell populations, anatomical locations and diseases and to extract gene, RNA or protein names in these contexts.

**Results:** OntoContext, a new python package contains two modules. The “annot” module for “annotation” function, is based on combination of morphosyntactic labelling and exact matching and on dictionaries derived from the Cell Ontology, the UBERON Ontology (anatomical context), the Human Disease Ontology and geniatagger, (which contains particular tags for gene-related names). The “annot” output is used as input for the second module “crisscross” generating lists of gene-related names obtained by crossing annotations from the three mentioned ontologies. OntoContext showed better performances than NCBO Annotator after evaluation on two text corpuses. OntoContext is freely available in the pypi.

**Availability:** https://pypi.python.org/pypi/OntoContext and https://github.com/walidbedhiafi/OntoContext1.

## 1 Introduction

Information from the scientific literature in the biomedical domain can be used to contextualize gene expression and protein involvement in various biological situations. The number of available biomedical texts is huge and constantly increasing. For example, “PubMed comprises more than 26 million citations for biomedical literature from MEDLINE, life science journals and online books. Citations may include links to full-text contents from PubMed Central and publisher web sites”(Dan Corlan, 2012). There is therefore a growing interest for scientists to extract automatically relevant information from these texts. Several informatics tools have been developed for gene interaction (Mallory *et al*., 2016), gene-disease (An *et al*., 2016), disease-drug (Bravo *et al*., 2015) extraction from scientific and medical texts. Several community challenges for text-mining in biology have been organized to cross-evaluate these tools (Krallinger *et al*., 2008; Ananiadou *et al*., 2015; Huang and Lu, 2016).

The annotation of biological entities in texts has emerged over the 1990s (Ananiadou *et al*., 2006). Biological advances, especially in genetics and molecular biology, have greatly contributed to increase publications and biological databases (Kersey *et al*., 2015). This has encouraged the emergence of new biology and\or computer science research fields including biotext mining (Ananiadou *et al*., 2006). In particular, algorithms were developed to respond to the needs for protein and gene names unambiguous identification (Denys *et al*., 1998) and protein function annotation (Andrade and Valencia, 1998). In the 2000s, approaches of exact matching were used for biotext annotation, benefiting from the stable development of databases serving to construct dictionaries. An original method for text alignment with dictionaries of protein and gene names was developed using “BLAST” (Krauthammer *et al*., 2000). These techniques have the disadvantage to be slow and to consume large computing capacity, especially with large dictionaries. However, they ensure good performance. Later, annotation techniques were improved by integrating machine learning and classification algorithms to increase the performance of exact matching (Tsuruoka *et al*., 2005). The integration of rule-based machine learning and classification approaches for the production of hybrid tools has helped to address new issues. This led to identify new biological concepts from the literature (Kim *et al*., 2015), extraction of protein interactions (Huang *et al*., 2004), identification of disease and gene relationships (Bravo *et al*., 2015; Tiffin *et al*., 2005; Kaur *et al*., 2014), or drug interactions (Iyer *et al*., 2014). These approaches highly depend on the training corpus (Jonnalagadda *et al*., 2012) and therefore are difficult to generalize.

In parallel, efforts have been provided for homogenization and standardization of “biological” languages. Several international initiatives, such as the Unified Medical Language System (UMLS), HUGO (Povey *et al*., 2001), Gene Ontology (Ashburner *et al*., 2000) and the OBO (Open Biomedical Ontologies) initiative (Smith *et al*., 2007), helped changing the paradigm of biotext mining. These initiatives have enabled the emergence of terminology shared by the whole community and used as stable dictionaries, recently extended to concept synonyms. Consequently, several annotation tools were developed relying on such standard concept dictionaries. The OBO annotator (Groza *et al*., 2015) is an annotation tool developed on the basis of the “Human Phenotype Ontology” in order to annotate similar characters between patients in different textual sources. The National Center of Biomedical Ontology Annotator (NCBO Annotator) tool uses multiple sources (ontologies, terminologies and databases) for biotext annotation (Jonquet *et al*., 2009). NCBO Annotator, based on a learning and exact matching algorithm “Mgrep”, and trained on limited sources and ontologies, shows average performances and is known as a reference for biotext annotation (Shah *et al*., 2009).

In this paper, we present OntoContext, a new python Package for biotext annotation using Ontologies to define Context. OntoContext was developed to automatically identify genes and gene products involved in a particular context: for example, cell population in a given anatomical localisation related to a physio-pathological state. To this end, we chose to annotate texts for concepts derived from ontologies. The described annotation method is based on natural processing language (NLP) from already structured or semi-structured information sources. First, ontologies are used as structured sources of information to construct specific morphosyntactic label dictionaries of each domain. Second, texts are morphosyntactically labelled and screened for ontology concepts stored in these dictionaries using label exact matching. Third, texts are scanned to identify and extract names of genes and gene products using the geniatagger.0.1 package (Tsuruoka *et al*., 2005). Fourth, texts are grouped according to a set of shared concepts across the three used ontologies. Fifth, the names of genes and gene products are extracted according to the defined context. Evaluation of two reference corpuses shows that OntoContext performs better that NCBO Annotator.

## 2 Material & Methods

### 2.1. Morphosyntactic labelling

OntoContext relies on the Part Of Speech Tagging morphosyntactic labelling technique (Baud *et al*., 1998; Brown, 2010) to identify targets in the literature. This technique allows transforming an explicit text in its syntactic representation. We define the morphosyntactic structure for each term, using the NLTK (Natural Language ToolKit) python package (Bird *et al*., 2009).

Ontologies, databases and construction of dictionaries

According to the context of our research, the OntoContext workflow relies on dictionaries (detailed in Supplementary Table S1) derived from three ontologies: the Cell Ontology for the cellular context (Bard *et al*., 2005), the UBERON Ontology for the anatomical context (Mungall *et al*., 2012), and the Human Disease Ontology for the pathological context (Kibbe *et al*., 2014). We used the available versions of these ontologies downloaded on 13/04/2015. We also used additional dictionaries, derived from previous or reduced versions of the ontologies. For each ontology, we build:

1. a synonym table associating all concepts in the ontology to their synonyms and plural form
2. a term-label table (called “ontology-derived dictionary”) containing all concepts, synonyms and plural form associated to their respective morphosyntactic label
3. a table containing parent concepts and their derived-children.

Note that dictionaries built here are not extended to term-derived adjectives. We also use non-hierarchically organized dictionaries derived from the corpus annotations BioText and CRAFT 1.0, containing only terms and their morphosyntactic label. All these table are grouped into the concept.db database, built using the SQLite database management system (Kreibich, 2010).

### 2.2. Text annotation

Our annotation method is based on the following steps as schematized on Figure 1: We use the NLTK modules to parse and label terms derived from ontologies (Figure 1A) and text sentences (Figure 1B). The implemented method takes text files as Input. After parsing and labelling sentences (step1), OntoContext looks for labels also found in the selected dictionary (step2). From these labels, OntoContext identify terms that match with the term-label table (step3, step4). This process is iterated for each candidate term (step5). For gene name retrieving, we use the geniatagger.0.1 package (Tsuruoka *et al*., 2005). The selected annotations are stored in a new table. These steps are coded into the “Annotation” function in OntoContext.

**Fig. 1.**
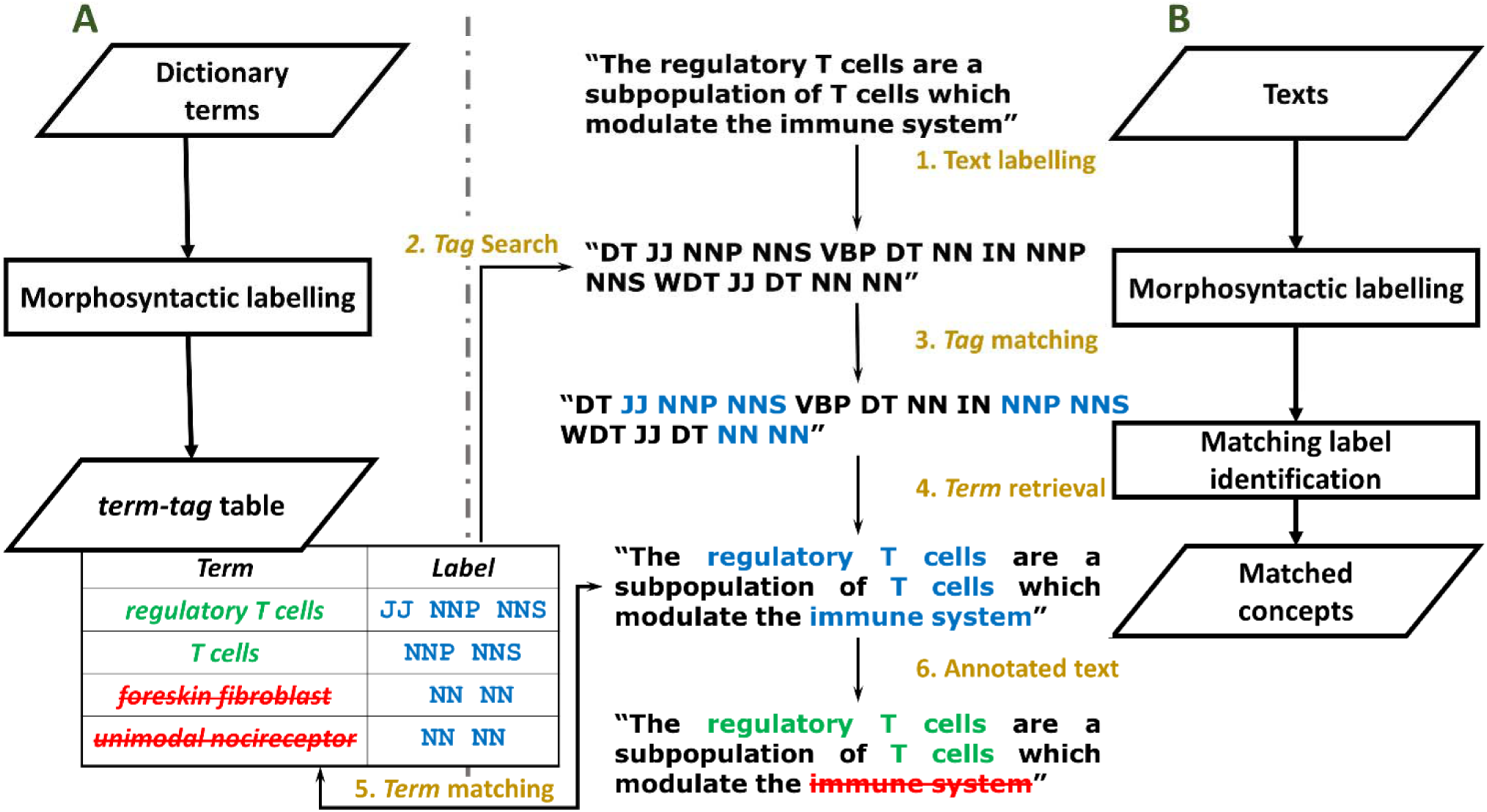
Principle of the OntoContext annotation algorithm. (A) Concepts with their synonyms are extracted from ontologies and plural forms are added to build a dictionary. These terms are labelled morphosyntactically using the NLTK package. This step generates the term-label table. An example of this table is shown with a term and a label column. (B) 1/ Texts of interest are parsed and labelled using the NLTK package. 2-3/ Matching labels with the term-label table are identified (in blue or light grey). 4/ The corresponding expressions are retrieved (in blue or light grey). 5/ These retrieved expressions are used to query the ontology-derived dictionary (term-label table) by exact term matching. 6/ Only expressions matching with the term-label table are kept for the text annotations (in green or light grey). Barred terms (or in red) are not considered as annotations. “DT”, determiner; “JJ”, adjective; “NNP”, singular proper nouns; “NNS”, plural noun; “VBP”, verb in the present tense; “NN”, common singular noun; “WDT”, wh-determiner.

### 2.3. Text contextualization and Gene/RNA/Protein name retrieval

After text annotation and matched concept identification with the dictionaries, children and synonyms are retrieved from each ontology. Concepts found at least once among the annotated texts are listed and offered for multiple selections. This step is guided by the indication of the concept frequency in the annotated texts (defined as the number of texts where the concept and its children are cited). Finally, we recover a list of gene-related names from this group of contextualized texts by querying the “result” table. The user is provided with the OntoContext graphical module “crisscross” developed using the Tkinter Package (Lundh, 1999), to visualize and select terms of interest in order to contextualize texts across ontologies.

### 2.4. Validation methodology

To evaluate OntoContext, we have selected two published text corpuses and have confronted OntoContext annotation performances against manually annotated texts used as reference (Supplementary Table S1). The CRAFT 1.0 Corpus annotated by experts for cell population concepts is a set of 67 full texts (Bada *et al*., 2012). The manual annotations are based on the 2012 Cell Ontology. The BioText corpus annotated by experts for pathological concepts includes 141 abstracts (Rosario and Hearst, 2004). We did not validate OntoContext for anatomical concepts in the absence of a reference corpus.

For each comparison, three performance criteria were calculated:

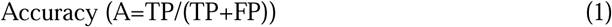

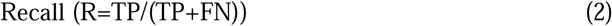

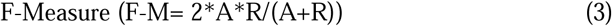

with TP = True Positives, FP = False Positives, FN = False Negatives.

We also designed a validation protocol based on these measures to compare the OntoContext performances against NCBO Annotator (Jonquet *et al*., 2009) (Figure 2).

**Fig. 2.**
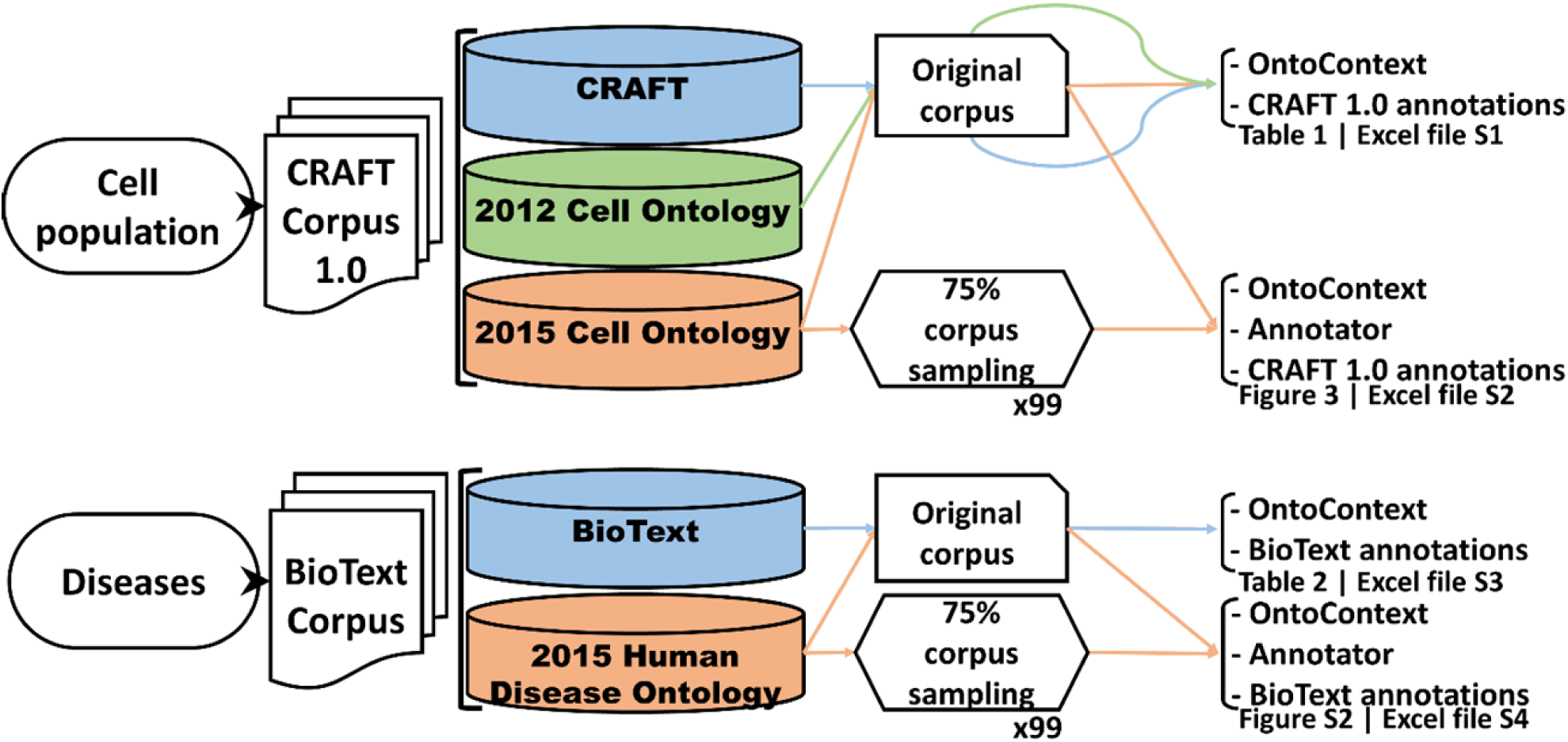
OntoContext validation tests and comparison with NCBO Annotator. The annotation of concepts such as “cell populations” or “diseases” is tested in various corresponding corpuses. Dictionaries derived from manual annotations of this corpus or from ontologies (Supplementary Table S1) are used in the validation process. Either the evaluation of corpus annotation is done with two tests, on the entire corpus or on a random sample of 75% of articles, repeated 99 times (x99). The performance comparison allows crossing the annotation results with those performed manually by expert or automatically by NCBO Annotator and OntoContext. The corresponding measures of the global Accuracy, Recall and F-Measure compare annotation sources and the method to obtain these annotations. The performance results are reported in Tables 1 and 2.

### 2.5. Implementation and availability

OntoContext was implemented in Python 2.7 (Lutz, 2013), a language chosen for its object-oriented ability that allows encapsulating and reusing methods and other libraries. The package and all its dependencies are available at the github (https://github.com/walidbedhiafi/OntoContext1) and pypi (https://pypi.org/project/OntoContext) repositories.

**Table 1.** OntoContext annotation performance of the CRAFT.1.0 corpus*

| Performance | CRAFT | 2012 Cell Ontology | 2015 Cell Ontology |
| --- | --- | --- | --- |
| Accuracy | 94% | 94% | 78% |
| Recall | 89% | 60% | 60% |
| F-Measure | 91% | 73% | 68% |

\*All 67 texts from the Craft 1.0 corpus have been annotated with OntoContext against three dictionaries. OntoContext annotation performance was evaluated compared to the CRAFT 1.0 expert manual annotations used as reference (see Supplementary Excel file S1 for detailed results). Figure 2 provides a schematized description of the evaluation workflow.

**Table 2.**
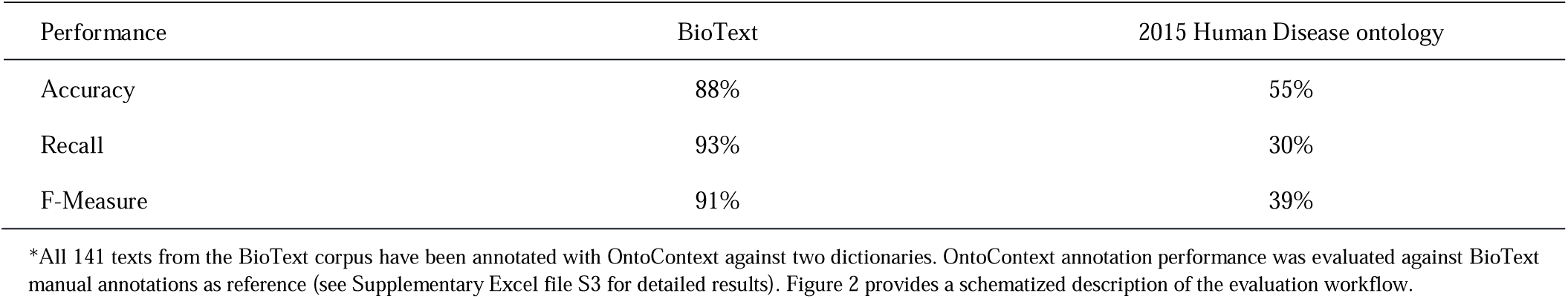
OntoContext annotation performance of the BioText corpus*

| Performance | BioText | 2015 Human Disease ontology |
| --- | --- | --- |
| Accuracy | 88% | 55% |
| Recall | 93% | 30% |
| F-Measure | 91% | 39% |

\*All 141 texts from the BioText corpus have been annotated with OntoContext against two dictionaries. OntoContext annotation performance was evaluated against BioText manual annotations as reference (see Supplementary Excel file S3 for detailed results). Figure 2 provides a schematized description of the evaluation workflow.

## 3 Results

OntoContext allows automatic annotation and contextualisation of biomedical texts in order to recover gene, RNA and protein names (abbreviated as gene-related names below). OntoContext is designed as a two-module package: The “annot” module contains the “annotation” function, based on the combination of the morphosyntactic labelling and the exact matching between target texts and ontology-derived dictionaries. The combination of these methods is faster than exact matching alone (Supplementary Note S1). The presented algorithms and the complexity calculation demonstrate how the “annot” module can be faster than classical exact matching algorithms.

The “annot” module takes successively as input the path to the folder containing the texts to annotate, the path to the geniatagger file (Tsuruoka *et al*., 2005) and Concept.db database, containing the Ontology-derived dictionaries provided by OntoContext. The user provides a name for the output table where the annotation results will be stored. These results consist in a list of concepts found in the annotated texts and their assignment to a given category (cell population, human disease, anatomical localization, gene-related names. Each row contains a concept, the related category and the text Id.

The second module named “crisscross” uses the previous “annot” output as input. “crisscross” contains five main interrelated function and steps, and a friendly interface:

1. Option step: The user can choose to display the annotation sorted either by alphabetic order or by citation frequency.
2. Extension step: The interface displays all terms found in the texts (“annot” output) along with their child terms obtained by automatic extension, classified into three lists, according to their assignment to a given ontology.
3. Selection step: The user can select sorted terms sorted either by alphabetical order or by frequency.
4. Contextualization step: A context of study is defined by three terms belonging to different ontologies. Contextualized texts sharing the same context of study are grouped (Supplementary Figure S1).
5. Gene-related name extraction: the gene-related names are extracted from the grouped texts obtained in contextualization step. Three lists are generated, at the levels of DNA (gene names), RNA (RNA transcript names) and proteins.

### 3.1. Evaluation of OntoContext performances

In order to evaluate OntoContext package, we selected two published corpuses: The CRAFT Corpus1.0 for cell population concepts and the BioText corpus for the disease concepts. Three performance criteria (Accuracy, Recall and F-Measure) were systematically calculated and compared to those obtained using NCBO Annotator (Jonquet *et al*., 2009) (Figure 3).

**Fig. 3.**
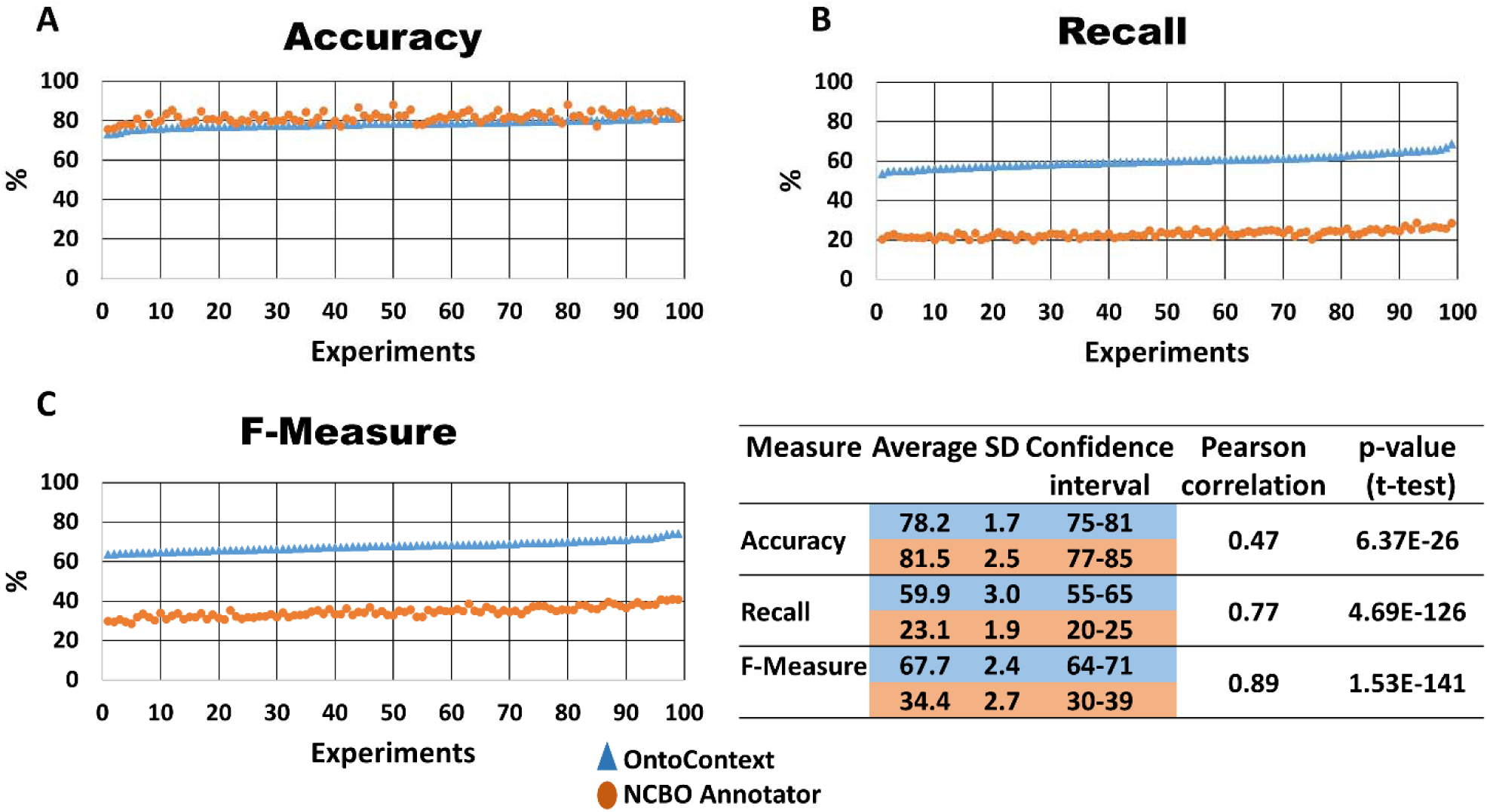
Comparing OntoContext and NCBO Annotator performances. A k-fold derived validation test for the OntoContext vs. NCBO Annotator tool performance comparison using the 2015 Cell Ontology-derived dictionary. We considered the obtained NCBO Annotator and OntoContext annotations for 35 of the 67 articles of the Craft corpus, and for each validation round, we drew 75% (27) of these 35 articles and assessed performances. Each pair of blue or light grey (OntoContext) and orange or dark grey (NCBO Annotator) points represents one of the 99 measures of Accuracy (A), Recall (B) and F-Measure (C) in percentage. Results are sorted by increasing OntoContext performance values. The performances of both tools are represented on the same experience. The table summarizes the mean values, with the standard deviations (SD). The confidence interval was calculated using the 95% quantile. The correlation test is Pearson correlation between the OntoContext values and the NCBO annotator values. The p-value was calculated based on paired t-test. The results of NCBO annotator are presented above the results of OntoContext in the table.

#### 3.1.1. OntoContext annotation of the Craft 1.0 corpus

For the CRAFT 1.0 corpus (Bada *et al*., 2012), we evaluated the performance of OntoContext using three different dictionaries: a reference dictionary based on the manual annotation of CRAFT 1.0 corpus (CRAFT-derived dictionary), a dictionary derived from the 2012 Cell Ontology and a dictionary derived from the 2015 Cell Ontology (Supplementary Table S1). Results are presented in Table 1 and Supplementary Excel file S1. The Accuracy values using the CRAFT-derived and 2012 Cell Ontology-derived dictionaries are the same (94%). Accuracy decreases to 78% when using the 2015 dictionary, indicating lower specificity with more false positives. This can be explained by the Craft expert annotators using the 2012 ontology, that did not include terms that were later added and therefore found in the enriched 2015 Cell Ontology-derived dictionary. The Recall measure is higher for 2015 Cell ontology with an 18-point increase between 2012 and 2015 Cell Ontology-derived dictionaries, but lower when using the CRAFT-derived dictionary.

Looking deeper in the results (Supplementary Excel file S1), some weaknesses in the OntoContext annotation are detected: since the ontology-derived concept dictionaries contain nouns but not derived adjectives, the latter are ignored (e.g. Cell / Cellular, Neuron / Neuronal…), at variance to manual extracted dictionary (CRAFT-derived dictionary). Another weakness consists in citation of concepts by the manual annotators in two parts of a sentence with other unrelated words in-between such as “CELLS IN […] MESENCHYMAL REGION”. When we ignore such interrupted concepts and recalculate the performances, Recall and F-Measure values slightly improve: For the manual CRAFT-derived dictionary, F-Measure increases from 91% to 94%, from 73% to 76% for the 2012 Cell Ontology-derived dictionary, and from 68% to 70% for the 2015 Cell-derived dictionary (not shown).

We then compared the OntoContext and NCBO Annotator performance using 35 out of 67 texts (50%) among the CRAFT.1.0 Corpus. NCBO Annotator is more specific than OntoContext (Accuracy of 82% vs 79%) but less sensitive, more terms being recognized by OntoContext (Recall of 23% vs 59%) (Supplementary Excel file S2). We performed a 99-round validation by taking randomly at each round a subset of 27 texts among the 35 texts (75%), annotating each subset by OntoContext and NCBO Annotator and measuring the annotation performances (Figure 3). OntoContext shows significantly higher F-Measures than Annotator (p-value<0.001). OntoContext systematically outperforms NCBO Annotator for Recall, the Accuracy values being closer.

#### 3.1.2. OntoContext annotation of the BioText corpus

In order to evaluate OntoContext for disease annotations, we used the BioText Medical corpus (141 abstracts) (Rosario and Hearst, 2004) and assessed its performances using two dictionaries: the BioText-derived dictionary established in 2004 by experts and used as reference for annotation and the 2015 Human Disease-derived dictionary (Supplementary Table S1). The BioText-derived dictionary not being generated from an ontology and thus not organized hierarchically, the extension step of the “crisscross” module cannot be performed. OntoContext applied to the BioText-derived dictionary reveals that some annotations include two disease concepts: for example, “PROSTATE CANCER” is a disease but the included term “CANCER” is also a disease. Therefore, OntoContext will annotate this term twice, as “CANCER” and “PROSTATE CANCER”. This explains, in part, that Accuracy is not optimum though the dictionary used to annotate the BioText Medical corpus, derived from this corpus.

Results for assessment of the OntoContext annotations of this corpus according to both dictionaries are presented in Table 2. Again, performances depend on the dictionary used. With the 2015 Human Disease-derived dictionary, we lose 33 points of Accuracy and 63 points of Recall as compared to the BioText-derived dictionary. This causes a 52-point loss for the F-Measure. This is likely due to significant differences with the Bio-Text-derived dictionary content that was originally used to annotate manually the BioText corpus in 2004. We then compared OntoContext to NCBO Annotator performances taking into account the 100 first texts among 141 texts of the BioText corpus along with the 2015 Human Disease-derived dictionary, despite of the relative weak performances obtained with this dictionary. Indeed, we were limited in our choice by the fact that NCBO annotator uses the 2015 Human Disease Ontology. In supplementary Excel file S4, we present the comparative annotation performances of these tools.

OntoContext recognizes more concepts (102) than NCBO Annotator (77 concepts) and, OntoContext is more specific than the NCBO Annotator (Accuracy of 60% vs 57%) and more sensitive, less false negative terms for OntoContext (Recall of 30% vs 21%) (Supplementary Excel file S4).

We then performed a 99-round validation by taking randomly at each round a subset of 75 texts among the 100 texts already analysed and measuring the annotation performances using OntoContext and NCBO annotator. OntoContext has significantly higher overall performances for all criteria considered than NCBO Annotator (p-value<0.001) and systematically outperforms NCBO Annotator for Recall and F-Measure. For Accuracy, values are close with OntoContext being better in 79 validation rounds out of 99 (Supplementary Figure S2).

### 3.2. Real Case study

In order to further validate our method, we applied OntoContext to a corpus of abstracts generated from PubMed in order to identify genes, RNA and proteins associated with a specific context. We queried PubMed with “(aging OR longevity) AND immune system”, restricted to MeSH terms and filtered for abstracts since 1980. This query was launched on August 1st, 2016 and retrieved 9,930 abstracts composed by 142,613 sentences representing 2 642,035 words. Abstracts were downloaded and annotated using the OntoContext “annot” module with the 2015 Cell-derived dictionary, 2015 Human Disease-derived dictionary and the 2015 UBERON-derived dictionary along with the geniatagger.1.0 package. The results are summarized in Figure 4A. OntoContext detected 34,350 cell population concepts, 17,260 anatomical concepts and 81,160 pathologies, and 3,063 gene-related names. Intriguingly, no gene-related names were retrieved for more than 7,000 abstracts. Looking closer at the annotations, we found that concepts such as “CD4-POSITIVE, CD25-POSITIVE, ALPHA-BETA REGULATORY T CELL” are only annotated as cell populations even though they contain molecular markers (“CD4” and “CD25”) that should also be recognized as gene-related names. This is due to geniatagger parsing all the “CD4-POSITIVE” or “CD25-POSITIVE” as a whole word. We thus performed a second round of annotation after removing all the “+” and “-“characters. This improved slightly the gene-extraction performance with more than 100 new texts being annotated with gene-related names, for a total of 3,988 gene-related names (Figure 4B).

**Fig. 4.**
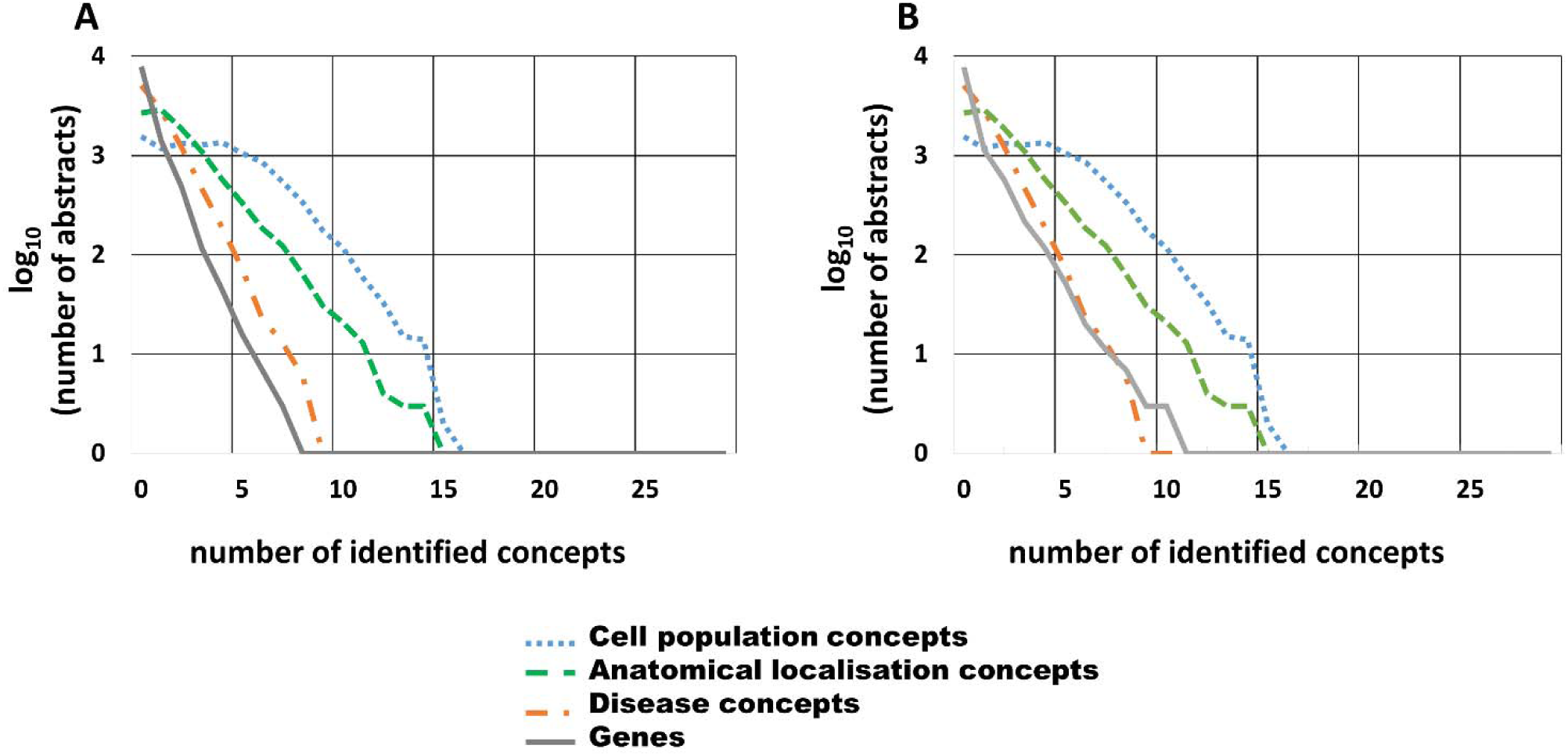
Cumulative distribution of the number of OntoContext-identified concept category terms per text analysed in a 9,930 corpus of PubMed abstracts. Cumulative distribution are expressed as log10 of the number of abstracts including terms related to cell populations, anatomical localisations, diseases and gene-related names for (A) a first round of standard OntoContext annotation results and (B) a second round of gene annotation removing the “+” and “-“ signs (see Results for details). The 9,930 PubMed abstract corpus was obtained by a query on the “immune system” and “aging” MeSH terms.

## 4 Discussion

In this paper, we present OntoContext, a python package developed for the automatic annotation of biomedical texts according to ontology-derived dictionaries describing a biomedical context such as cell population, anatomical location and diseases, in addition to protein, RNA or gene name information. Although, specialized ontologies and databases are available, little interrelations between them in terms of biological meaning is provided. Our aim was to sort texts based first on a PubMed search with MeSH terms, then their contexts of study derived from ontology concept labelling and to retrieve gene-related names cited in these contexts.

The two modules “annot” and “crisscross” developed here allow the annotation of a text corpus against a concept database and the intersection between texts annotated by these concepts, respectively. These two modules can be used separately or successively. Noteworthy, “crisscross” depends on the output of “annot”, the performance of which is crucial for the global performance of OntoContext. For that reason, we focused our performance assessment on “annot”. With our original objective in mind, we needed an organized and controlled vocabulary and thus chose to build dictionaries derived from ontologies. Ontologies represent standard and controlled vocabularies shared by the scientific community and hierarchically organized, hence allowing clustering. For gene name recognition, we selected the already available geniatagger.0.1 package (Tsuruoka *et al*., 2005). The exact matching method has the advantage to easily annotate with controlled vocabulary. By implementing an exact matching algorithm based on the morphosyntactic labelling of text and dictionary terms, we have here overcome the relative slowness of this method (Navarro and Fredriksson, 2004), and OntoContext appears faster than a classic exact matching algorithm, as demonstrated in Supplementary Note S1. For the morphosyntactic labelling, we used the NLTK python package (Bird *et al*., 2009) included into the “Annotation” function to label the dictionaries terms and texts. These labels may differ between the database and the sentences depending on several criteria such as the uppercase and the lowercase writing and the plural forms. For that reason, a pre-treatment step of data consisting in adding plural, synonyms, considering lower and uppercase improves OntoContext that has better performances as compared to NCBO annotator (Jonquet *et al*., 2009). However, both tools are still unable to recognize adjectives, contrarily to experts. The most important performance criteria seem related to the date at which the dictionary used for annotation relatively to the text corpus is generated. Indeed, new concepts are created along with the evolution of knowledge, leading to changes in vocabularies and creating difficulties for annotation. These difficulties are not related to the method but rather to the quality of the vocabulary used comparatively to the reference. Indeed, we observed variations in OntoContext annotation results when we used different dictionaries to analyse the same text corpus.

For the evaluation of an annotation method, it is important to use the same dictionary as a reference. We indeed assessed OntoContext using CRAFT-derived dictionary to annotate CRAFT.1.0 corpus (Bada *et al*., 2012) and obtained more than 90% Accuracy. This result indicates good performance of our “annot” module compared to other methods, similar with that reported by Kim et al. (Kim *et al*., 2015), when geniatagger accuracy is about 54% for cell population identification (Tsuruoka *et al*., 2005). We have to notice that previous tools did not use the same corpus for evaluation as OntoContext. These tools being developed on a training dataset, their performances decrease when corpuses are changed. For OntoContext, we have developed a method that overcomes these drawbacks since it is based on exact matching and morphosyntactic labelling, using ontology-derived dictionaries.

For gene-related name annotation, many databases were developed but they are species dependent: the “gene name database” concerns human (Povey *et al*., 2001), the “mouse genome informatics database” concerns mouse (Bult *et al*., 2016) and the “WormBase” concerns nematode (Harris *et al*., 2004). Other limits may exist such as for the Protein Ontology that contains only coding gene names (Natale *et al*., 2014). The inclusion in the “annot” module of the geniatagger package (Tsuruoka *et al*., 2005) overcomes these limits allowing to annotate genes, RNA and protein names from any species with good performances.

As a whole, OntoContext performance is influenced essentially by the quality and date of the dictionaries. Using newest version of ontologies is a guarantee for better annotation results, whatever the text corpus considered for annotation.

In its present stage of development, the OntoContext package can be used for different purposes. Here we present an example of annotation of an abstract corpus from PubMed. We used PubMed abstracts and not full texts here because they are freely available and easily extractable, although we realize that using abstracts only imposes a limitation on the number of identifiable gene-related names. OntoContext can also be applied to annotate information from clinical texts or from specialized databases (transcriptomic, protein, genomic…) in order to generate new knowledge and life process modelling. This new tool needs further evaluation for different use and improvement for better performance such as the speed and accuracy of the annotation method.

## Supporting information

Supplementary_Excel_file_S1 2012_Cell Ontology derived

Supplementary_Excel_file_S1 2015_Cell Ontology derived

Supplementary_Excel_file_S1 CRAFT 1.0 derived

Supplementary_Excel_file_S2 Same_dictionary_OntoC_Annot

Supplementary_Excel_file_S3 2015 Human Disease derived

Supplementary_Excel_file_S3 BioText derived

Supplementary_Excel_file_S4 Annotator_vs_OntoC

## Acknowledgements

We thank Dr. Ben Miled Slimen and Dr. Mhamdi Faouzi for their help.

