## Supplementary_Excel_file_S1 2012_Cell Ontology derived for "OntoContext, a new python package for gene contextualization based on the annotation of biomedical texts"

| Num file | Reference CRAFT 1.0 Annotation | OntoContext Annotation | TP | FP | FN |
|---|---|---|---|---|---|
| file1 | CONES | CELL | | 1 | 1 |
| | MACROPHAGES | CELLS | 1 | | |
| | CELLS | RETINAL GANGLION CELL | 1 | | |
| | RETINAL GANGLION CELL | MACROPHAGES | 1 | | |
| | ROD ... PHOTORECEPTORS | | | | 1 |
| | RODS | | | | 1 |
| | CONE PHOTORECEPTORS | | | | 1 |
| file2 | CELL | CELL | 1 | | |
| | STEM ... CELLS | CELLS | 1 | | 1 |
| | CELLS | | | | |
| file3 | OLIGODENDROCYTE | | | | 1 |
| | CELLULAR | | | | 1 |
| file4 | CELLS | CELLS | 1 | | |
| | CELL | CELL | 1 | | |
| | BLASTOMERES | BLASTOMERES | 1 | | |
| | ENDODERM CELLS | ENDODERM CELLS | 1 | | |
| | STEM CELLS | STEM CELLS | 1 | | |
| | EUKARYOTIC CELLS | EUKARYOTIC CELLS | 1 | | |
| file5 | CELLULAR | EPITHELIAL CELLS | 1 | | 1 |
| | EPITHELIAL CELLS | CELLS | 1 | | |
| | CELLS | CELL | 1 | | |
| | CELL | OSTEOBLASTS | 1 | | |
| | B LYMPHOCYTES | EPITHELIAL CELL | 1 | | 1 |
| | EPITHELIAL CELL | LYMPHOCYTES | | 1 | |
| | OSTEOBLASTS | MUSCLE CELL | 1 | | |
| | MUSCLE CELL | MACROPHAGE | 1 | | |
| | MACROPHAGE | | | | |
| file6 | SUPPORTING CELL | HAIR CELLS | 1 | | 1 |
| | NEURON | MOTONEURONS | 1 | | |
| | SUPPORTING CELLS | INNER HAIR CELLS | 1 | | 1 |
| | NEURON AFFERENTS | HAIR CELL | 1 | | 1 |
| | OUTER HAIR CELL | OUTER HAIR CELLS | 1 | | |
| | SENSORY NEURONS | CELLS | 1 | | |

| | |
|---|---|
| Accuracy | 0,94 |
| Recall | 0,60 |
| F-Measure | 0,73 |

| | | | | | |
|---|---|---|---|---|---|
| | PRIMARY NEURONS | NEURON | 1 | | |
| | INNER ... HAIR CELLS | SENSORY NEURON | 1 | | 1 |
| | VESTIBULAR HAIR CELL | PILLAR CELLS | 1 | | 1 |
| | CELL | CELL | 1 | | |
| | NEURONS | NEURONS | 1 | | |
| | NEURONAL | OUTER HAIR CELL | 1 | | 1 |
| | INNER HAIR CELL | SENSORY NEURONS | 1 | | |
| | HAIR CELL | PRIMARY NEURONS | 1 | | |
| | OUTER HAIR CELLS | | | | |
| | CELLS | | | | |
| | SENSORY NEURON | | | | |
| | CELLULAR | | | | 1 |
| | PILLAR CELLS | | | | |
| | HAIR CELLS | | | | |
| | MOTONEURONS | | | | |
| | INNER HAIR CELLS | | | | |
| | DEITERS' CELLS | | | | 1 |
| file7 | EOSINOPHILES | CARDIOMYOCYTES | 1 | | 1 |
| | PLATELETS | LEUCOCYTES | 1 | | |
| | CELLULAR | RED BLOOD CELLS | 1 | | |
| | LEUCOCYTES | CELLS | 1 | | |
| | RED BLOOD CELLS | PLATELETS | 1 | | |
| | CELLS | ERYTHROCYTES | 1 | | |
| | CARDIOMYOCYTES | BLOOD CELL | | 1 | |
| | BASOPHILES | ERYTHROCYTE | 1 | | 1 |
| | ERYTHROCYTES | CELL | 1 | | |
| | ERYTHROCYTE | PLATELET | 1 | | |
| | CELL | BLOOD CELLS | | 1 | |
| | PLATELET | MONOCYTES | 1 | | |
| | MONOCYTES | RED BLOOD CELL | 1 | | |
| | RED BLOOD CELL | LYMPHOCYTES | 1 | | |
| | LYMPHOCYTES | LEUKOCYTES | 1 | | |
| | NEUTROPHILES | | | | 1 |
| | LEUKOCYTES | | | | |
| file8 | CELL | CELL | 1 | | |
| | NEURONAL | NEURONAL | | | 1 |

| | | | | |
|---|---|---|---|---|
| file9 | SPERM | SPERM | 1 | |
| | SPERMATID | SPERMATID | 1 | |
| | CELLS | CELLS | 1 | |
| | NEURON | NEURON | 1 | |
| | T-CELL | T-CELL | 1 | |
| | CELL | CELL | 1 | |
| | NEURONS | NEURONS | 1 | |
| | ERYTHROID CELLS | | | 1 |
| | NEURONAL | | | 1 |
| | OLFACTORY NEURON | | | 1 |
| | OLFACTORY NEURONS | | | 1 |
| file10 | NEURON | CELL | 1 | |
| | CELLS | CELLS | 1 | |
| | NEURONAL CELLS | NEURONS | 1 | 1 |
| | ASTROCYTES | NEURON | 1 | |
| | CELL | ASTROCYTES | 1 | |
| | NEURONS | | | |
| | CELLULAR | | | 1 |
| | NEURONAL CELL | | | 1 |
| file11 | PIGMENT CELL | PIGMENT CELL | 1 | |
| | MELANOCYTE | MELANOCYTE | 1 | |
| | STEM CELLS | CELLS | 1 | 1 |
| | DERMAL PAPILLA CELLS | MELANOBLAST | 1 | 1 |
| | CELLS | CELL | 1 | |
| | MELANOBLAST | KERATINOCYTES | 1 | |
| | DERMAL PAPILLAE CELLS | MELANOCYTES | 1 | 1 |
| | PIGMENT-CELL | PIGMENT CELLS | 1 | 1 |
| | CELL | MESENCHYMAL CELLS | 1 | |
| | KERATINOCYTES | MELANOBLASTS | 1 | |
| | CELLULAR | | | 1 |
| | MELANOCYTES | | | |
| | MESENCHYMAL CELLS | | | |
| | PIGMENT CELLS | | | |
| | MELANOBLASTS | | | |
| file12 | FIBROBLASTS | FIBROBLASTS | 1 | |
| | ENDOTHELIAL CELL | ENDOTHELIAL CELL | 1 | |

| | | | | | |
|---|---|---|---|---|---|
| | CELLULAR | EPITHELIAL CELLS | 1 | | 1 |
| | EPITHELIAL CELLS | CELLS | 1 | | |
| | CELLS IN ... MESENCHYMAL REGIONS | CELL | 1 | | 1 |
| | CELLS | PLATELET | 1 | | |
| | CELLS IN ... EPITHELIAL ... REGIONS | | | | 1 |
| | EPITHELIAL CELLS ... IN ... LUNGS | | | | 1 |
| | CELL | | | | |
| | PLATELET | | | | |
| | EPITHELIAL ... CELLS | | | | 1 |
| | MESENCHYME CELLS | | | | 1 |
| | EPITHELIAL CELLS IN ... LUNGS | | | | 1 |
| | LUNG EPITHELIAL CELL | | | | 1 |
| | LUNG EPITHELIAL CELLS | | | | 1 |
| file13 | CELL | CELL | 1 | | |
| | EPITHELIAL CELLS | FIBROBLASTS | 1 | | 1 |
| | CELLS | CELLS | 1 | | |
| | FIBROBLASTS | EPITHELIAL CELLS | 1 | | |
| | CELLULAR | ENTEROCYTES | 1 | | |
| | ENTEROCYTES | | | | |
| file14 | NEURONAL | CELL | | 1 | |
| | DOPAMINE CELLS | CELLS | 1 | | 1 |
| | CELLS | | 1 | | |
| | CELLULAR | | | | 1 |
| | DOPAMINERGIC CELL | | | | 1 |
| | DOPAMINERGIC CELLS | | | | 1 |
| file15 | CELL | CELL | 1 | | |
| | OOCYTE | OOCYTE | 1 | | |
| | CELLS | CELLS | 1 | | |
| | CELLULAR | | | | 1 |
| file16 | PROERYTHROBLAST | PROERYTHROBLAST | 1 | | |
| | RBCS | RBCS | 1 | | |
| | STEM CELL | STEM CELL | 1 | | |
| | ERYTHROCYTES | ERYTHROCYTES | 1 | | |
| | STEM CELLS | STEM CELLS | 1 | | |
| | HEMATOPOIETIC CELLS | RETICULOCYTE | 1 | | 1 |

| | | | | | |
|---|---|---|---|---|---|
| | RETICULOCYTE | HEMATOPOIETIC STEM CELL | 1 | | |
| | WHITE BLOOD CELLS | RED BLOOD CELL | 1 | | |
| | STEM ... CELLS | T-CELLS | 1 | | 1 |
| | HEMATOPOIETIC STEM CELL | CELL | 1 | | |
| | RED BLOOD CELL | HEMATOPOIETIC PROGENITOR CELLS | 1 | | |
| | T-CELLS | MESENCHYMAL CELLS | 1 | | |
| | CELL | EGGS | 1 | | |
| | RED ... BLOOD CELLS | WHITE BLOOD CELLS | 1 | | 1 |
| | HEMATOPOIETIC PROGENITOR CELLS | MYELOID CELLS | 1 | | |
| | HEMATOPOIETIC PROGENITORS | CELLS | 1 | | 1 |
| | ERYTHROID CELLS | FIBROBLASTS | 1 | | 1 |
| | MESENCHYMAL CELLS | CIRCULATING CELLS | 1 | | |
| | CELLS | FIBROBLAST | 1 | | |
| | ERYTHROID CELL | LYMPHOCYTES | 1 | | 1 |
| | APOPTOTIC CELLS | HEMATOPOIETIC STEM CELLS | 1 | | 1 |
| | MYELOID CELLS | MESODERMAL CELLS | 1 | | |
| | EGGS | ERYTHROID PROGENITOR CELLS | 1 | | |
| | ERYTHROID | MACROPHAGES | 1 | | 1 |
| | FIBROBLASTS | RED BLOOD CELLS | 1 | | |
| | CIRCULATING CELLS | BLOOD CELL | 1 | | |
| | FIBROBLAST | BLOOD CELLS | 1 | | |
| | MYELOID | PROERYTHROBLASTS | 1 | | 1 |
| | LYMPHOCYTES | | | | |
| | FIBROBLAST CELL | | | | 1 |
| | HEMATOPOIETIC STEM CELLS | | | | |
| | MESODERMAL CELLS | | | | |
| | ERYTHROID PROGENITOR CELLS | | | | |
| | MACROPHAGES | | | | |
| | RED BLOOD CELLS | | | | |
| | ERYTHROID PROGENITORS | | | | 1 |
| | BLOOD CELL | | | | |
| | BLOOD CELLS | | | | |
| | PROERYTHROBLASTS | | | | |
| file17 | CELL | CELL | 1 | | |
| | CELLS | APCS | | 1 | |
| | CELLS IN VIVO | CELLS | 1 | | 1 |
| | STEM CELLS | EMBRYONIC STEM CELLS | | 1 | |
| | APOPTOTIC CELLS | STEM CELLS | 1 | | 1 |

| | | | | | |
|---|---|---|---|---|---|
| file18 | FIBROCYTES | FIBROCYTES | 1 | | |
| | VESTIBULAR DARK CELLS | HAIR CELLS | 1 | | |
| | BASAL CELLS | BASAL CELLS | 1 | | |
| | HAIR CELLS | HAIR CELL | | 1 | |
| | EPITHELIAL CELLS | EPITHELIAL CELLS | 1 | | |
| | CELLS | CELLS | 1 | | |
| | STEM CELL | STEM CELL | 1 | | |
| | CELL | CELL | 1 | | |
| | HAIR CELL | BASAL CELL | | 1 | |
| | VESTIBULAR HAIR CELLS | VESTIBULAR DARK CELLS | 1 | | 1 |
| | TRANSITIONAL CELLS | | | | 1 |
| file19 | MOTOR NEURON | MOTOR NEURON | 1 | | |
| | ZYGOTES | NEURON | | 1 | |
| | ADIPOCYTE | ZYGOTES | 1 | | |
| | CELL | ADIPOCYTE | 1 | | |
| | MOTONEURON | CELL | 1 | | |
| | CELLULAR | MOTONEURON | 1 | | 1 |
| file20 | APOPTOTIC ... CELLS | CELL | 1 | | 1 |
| | APOPTOTIC CELL | CELLS | 1 | | 1 |
| | APOPTOTIC CELLS | EARLY ERYTHROBLAST | 1 | | 1 |
| | CELL | EPITHELIAL CELLS | 1 | | |
| | CELLS | ERYTHROBLAST | 1 | | |
| | CELLS IN ... MESENCHYME | ERYTHROCYTES | 1 | | 1 |
| | CELLULAR | FIBROBLASTS | 1 | | 1 |
| | EARLY ERYTHROBLAST | MACROPHAGE | 1 | | |
| | EPITHELIAL CELLS | MACROPHAGES | 1 | | |
| | ERYTHROCYTES | MEGAKARYOCYTES | 1 | | |
| | FIBROBLASTS | MONOCYTE | 1 | | |
| | HEMATOPOIETIC STEM CELLS | NEUTROPHILS | 1 | | 1 |
| | MACROPHAGE | PERITONEAL MACROPHAGES | 1 | | |
| | MACROPHAGES | PHAGOCYTES | 1 | | |
| | MEGAKARYOCYTES | PHOTORECEPTOR CELLS | 1 | | |
| | MESENCHYMAL ... CELLS | PIGMENTED EPITHELIAL CELLS | 1 | | 1 |
| | MONOCYTE | THYMOCYTES | 1 | | |
| | NEURONAL | | | | 1 |

| | | | | |
|---|---|---|---|---|
| | NEURONAL CELLS | | | 1 |
| | NEUTROPHILS | | | |
| | PERITONEAL MACROPHAGES | | | |
| | PHAGOCYTES | | | |
| | PHOTORECEPTOR CELLS | | | |
| | PIGMENTED CELLS | | | 1 |
| | PIGMENTED EPITHELIAL CELLS | | | |
| | PLATELET | | | 1 |
| | STEM ... CELLS | | | 1 |
| file21 | CARTILAGE CELLS | CELL | 1 | 1 |
| | CELL | CELLS | 1 | |
| | CELLS | CHONDROCYTES | 1 | |
| | CELLULAR | EGGS | 1 | 1 |
| | CHONDROCYTES | MACROPHAGES | 1 | |
| | EGGS | MONONUCLEAR CELLS | 1 | |
| | FIBROBLAST | NEUTROPHILS | 1 | |
| | HYPERTROPHIC CHONDROCYTES | OSTEOBLASTS | 1 | 1 |
| | MACROPHAGES | SYNOVIAL CELLS | 1 | |
| | MESENCHYMAL CELLS | | 1 | 1 |
| | MONONUCLEAR CELLS | | 1 | |
| | NEUTROPHILS | | | |
| | OSTEOBLASTS | | | |
| | SYNOVIAL CELLS | | | |
| | SYNOVIOCYTES | | | 1 |
| file22 | CELL | CELL | 1 | |
| | CELLS | CELLS | 1 | |
| | GLIA | MOTOR NEURON | 1 | 1 |
| | MOTOR NEURON | MOTOR NEURONS | 1 | |
| | MOTOR NEURONS | NEURON | 1 | |
| | NEURON | NEURONS | 1 | |
| | NEURONS | SENSORY NEURONS | 1 | |
| | SENSORY NEURONS | | | |
| file23 | CELL | CELL | 1 | |
| | CELLS | CELLS | 1 | |
| | CELLULAR | CNS NEURONS | 1 | 1 |
| | CNS NEURONS | NEURON | 1 | |

| | | | | |
|---|---|---|---|---|
| | GRANULE NEURON | NEURONS | 1 | 1 |
| | GRANULE NEURONS | | 1 | 1 |
| | NEURON | | 1 | |
| | NEURONAL | | 1 | 1 |
| | NEURONAL CELL | | | 1 |
| | NEURONAL CELLS | | | 1 |
| | NEURONS | | | |
| | PURKINJE NEURON | | | 1 |
| file24 | BASAL ... CELLS | CELL | 1 | 1 |
| | CELL | CELLS | 1 | |
| | CELLS | EPIDERMAL CELLS | 1 | |
| | CELLS OF ... ECTODERM | EPITHELIAL CELLS | 1 | 1 |
| | CELLS WITHIN ... EPIDERMIS | FIBROBLAST | 1 | 1 |
| | CELLULAR | HAIR CELL | 1 | 1 |
| | ECTODERMAL CELLS | HEPATOCYTES | 1 | 1 |
| | EPIDERMAL CELLS | KERATINOCYTE | 1 | |
| | EPITHELIAL ... CELL | KERATINOCYTES | 1 | |
| | EPITHELIAL CELLS | MESENCHYMAL CELLS | 1 | |
| | FIBROBLAST | T CELL | 1 | |
| | GRANULAR LAYER CELLS | | | 1 |
| | HAIR CELL | | | |
| | HEPATOCYTES | | | 1 |
| | KERATINOCYTE | | | |
| | KERATINOCYTES | | | 1 |
| | MESENCHYMAL CELLS | | | |
| | MOTILE ... CELLS | | | 1 |
| | SQUAMOUS CELL | | | 1 |
| | T CELL | | | |
| file25 | BIPOLAR CELL | CELL | 1 | 1 |
| | BIPOLAR CELLS | CELLS | 1 | 1 |
| | CELL | HORIZONTAL CELLS | 1 | |
| | CELLS | NEURONS | 1 | |
| | CONE | PHOTORECEPTOR CELLS | 1 | 1 |
| | HORIZONTAL ... CELL | ROD BIPOLAR CELLS | 1 | 1 |
| | HORIZONTAL ... CELLS | SENSORY NEURONS | 1 | |
| | HORIZONTAL CELL | | | 1 |
| | HORIZONTAL CELLS | | | |

| | | | | |
|---|---|---|---|---|
| | MÃ¼LLER CELLS | | | 1 |
| | NEURONAL | | | 1 |
| | NEURONS | | | |
| | OFF ... BIPOLAR CELLS | | | 1 |
| | ON ... BIPOLAR CELLS | | | 1 |
| | PHOTORECEPTOR | | | 1 |
| | PHOTORECEPTOR CELLS | | | |
| | PHOTORECEPTORS | | | 1 |
| | PHOTORECEPTORS ... R6 | | | 1 |
| | PHOTORECEPTORS R1 | | | 1 |
| | R1 ... PHOTORECEPTORS | | | 1 |
| | R6 PHOTORECEPTORS | | | 1 |
| | ROD BIPOLAR | | | 1 |
| | ROD BIPOLAR CELLS | | | |
| | ROD PHOTORECEPTORS | | | 1 |
| | RODS | | | 1 |
| | SENSORY NEURONS | | | |
| file26 | ASTROCYTES | ASTROCYTES | 1 | |
| | ASTROCYTIC | CARDIAC MYOCYTES | 1 | 1 |
| | CARDIAC MYOCYTES | CELL | 1 | |
| | CELL | CELLS | 1 | |
| | CELLS | EMBRYONIC STEM CELLS | 1 | |
| | CELLULAR | GRANULE CELL | 1 | 1 |
| | GLIAL | HEPATOCYTE | 1 | 1 |
| | GRANULE CELL | HEPATOCYTES | 1 | |
| | HEPATOCYTE | MYOCYTES | 1 | |
| | HEPATOCYTES | NEURON | 1 | |
| | LYMPHOCYTIC | NEURONS | 1 | 1 |
| | NEURON | OOCYTES | 1 | |
| | NEURONAL | PYRAMIDAL NEURONS | 1 | 1 |
| | NEURONS | STEM CELLS | 1 | |
| | OOCYTES | | | |
| | PHAGOCYTIC CELLS | | | 1 |
| | PURKINJE ... CELL | | | 1 |
| | PYRAMIDAL NEURONS | | | |
| | STEM CELLS | | | |
| file27 | ASTROCYTE | ASTROCYTES | 1 | 1 |

| | | | | | |
|---|---|---|---|---|---|
| | ASTROCYTES | BLOOD CELLS | | 1 | |
| | ASTROCYTIC | CELL | 1 | | 1 |
| | ASTROCYTIC CELL | CELLS | 1 | | 1 |
| | ASTROGLIAL ... CELL | CHROMAFFIN CELLS | 1 | | 1 |
| | ASTROGLIAL CELL | ENDODERM CELLS | 1 | | 1 |
| | ASTROGLIAL CELLS | FIBROBLASTS | 1 | | 1 |
| | CELL | GLIAL CELL | 1 | | |
| | CELLS | GLIAL CELLS | 1 | | |
| | CELLS OF ... STRATUM GRANULOSUM | MESODERMAL CELLS | 1 | | 1 |
| | CELLULAR | MYOBLAST | 1 | | 1 |
| | CHROMAFFIN CELLS | NEURON | 1 | | |
| | ENDODERM CELLS | NEURONS | 1 | | |
| | ENDODERMAL ... CELLS | PLATELET | 1 | | |
| | FIBROBLASTS | PURKINJE CELLS | 1 | | |
| | GLIAL | PYRAMIDAL NEURON | 1 | | 1 |
| | GLIAL CELL | PYRAMIDAL NEURONS | 1 | | |
| | GLIAL CELLS | RADIAL GLIAL CELLS | 1 | | |
| | MESODERMAL CELLS | RED BLOOD CELLS | 1 | | |
| | MYOBLAST | | | | |
| | NEURON | | | | |
| | NEURONAL | | | | 1 |
| | NEURONAL ... CELL | | | | 1 |
| | NEURONAL ... CELLS | | | | 1 |
| | NEURONAL CELLS | | | | 1 |
| | NEURONS | | | | |
| | PLATELET | | | | |
| | PURKINJE CELLS | | | | |
| | PURKINJE-CELL | | | | 1 |
| | PURKINJE-CELLS | | | | 1 |
| | PYRAMIDAL NEURON | | | | |
| | PYRAMIDAL NEURONS | | | | |
| | RADIAL GLIAL CELLS | | | | |
| | RED BLOOD CELLS | | | | |
| file28 | AFFERENT CELL | CELL | 1 | | 1 |
| | AFFERENT SENSORY NEURONS | CELLS | 1 | | 1 |
| | AFFERENTS ... NEURONS | MOTOR NEURON | 1 | | 1 |
| | APOPTOTIC CELLS | MOTOR NEURONS | 1 | | 1 |
| | CELL | NEUROBLAST | 1 | | |

| | | | | |
|---|---|---|---|---|
| | CELLS | NEUROBLASTS | 1 | |
| | CELLULAR | NEURON | 1 | 1 |
| | MOTOR NEURON | NEURONS | 1 | |
| | MOTOR NEURONS | PEPTIDERGIC NEURONS | 1 | |
| | NEUROBLAST | SENSORY NEURON | 1 | |
| | NEUROBLASTS | SENSORY NEURONS | 1 | |
| | NEURON | | | |
| | NEURONAL | | | 1 |
| | NEURONAL CELL | | | 1 |
| | NEURONS | | | |
| | PEPTIDERGIC NEURONS | | | |
| | SENSORY ... NEURONS | | | |
| | SENSORY NEURON | | | |
| | SENSORY NEURONS | | | |
| file29 | CELL | CELL | 1 | |
| | CELLS | CELLS | 1 | |
| | EGGS | EGGS | 1 | |
| | GRANULE CELL | GRANULE CELL | 1 | |
| | GRANULE CELLS | GRANULE CELLS | 1 | |
| | NEURON | NEURONS | 1 | |
| | NEURONAL | PURKINJE CELL | 1 | 1 |
| | NEURONAL CELL | SCHWANN CELL | 1 | 1 |
| | NEURONS | SCHWANN CELLS | 1 | |
| | PURKINJE CELL | | | |
| | SCHWANN CELL | | | |
| | SCHWANN CELLS | | | |
| | SCHWANN-CELL | | | 1 |
| | SPERMATOZOA | | | 1 |
| | STEM ... CELLS | | | 1 |
| file30 | CELL | CELL | 1 | |
| | CELLS | CELLS | 1 | |
| file31 | BLOOD CELL | BLOOD CELL | 1 | |
| | CELL | CELL | 1 | |
| | CELLS | CELLS | 1 | |
| | ENDOTHELIAL CELL | ENDOTHELIAL CELL | 1 | |
| | EPITHELIAL CELLS | EPITHELIAL CELLS | 1 | |

| | | | | | |
|---|---|---|---|---|---|
| | GOBLET CELL | GOBLET CELL | 1 | | |
| | GOBLET CELLS | GOBLET CELLS | 1 | | |
| | HEMATOPOIETIC CELL | IMMUNE CELL | 1 | | 1 |
| | IMMUNE CELL | IMMUNE CELLS | 1 | | |
| | IMMUNE CELLS | LYMPHOCYTE | 1 | | |
| | LYMPHOCYTE | LYMPHOCYTES | 1 | | |
| | LYMPHOCYTES | MAST CELL | 1 | | |
| | MAST CELL | MAST CELLS | 1 | | |
| | MAST CELLS | MONONUCLEAR CELL | 1 | | |
| | MONONUCLEAR CELL | NEUTROPHIL | 1 | | |
| | NEUTROPHIL | NEUTROPHILS | 1 | | |
| | NEUTROPHILS | PANETH CELL | 1 | | |
| | PANETH CELL | STEM CELL | 1 | | |
| | STEM CELL | T CELL | 1 | | |
| | T CELL | | | | |
| file32 | CELLS | CELL | | 1 | |
| | CELLS ... OF ... EPITHELIUM | CELLS | 1 | | 1 |
| | CELLULAR | NATURAL KILLER CELL | 1 | | 1 |
| | NATURAL KILLER CELL | RECEPTOR CELLS | | 1 | |
| | TASTE CELLS | TASTE RECEPTOR CELLS | 1 | | 1 |
| | TASTE RECEPTOR CELLS | | 1 | | |
| | TRCS | | | | 1 |
| file33 | CELL | CELL | 1 | | |
| | CELLS | CELLS | 1 | | |
| | GLIA | NEURONS | 1 | | 1 |
| | NEURONAL | | | | 1 |
| | NEURONES | | | | 1 |
| | NEURONS | | | | |
| file34 | CELLS | CELL | | | 1 |
| | HEPATOCYTE | CELLS | 1 | | |
| | MUSCLE PRECURSOR CELL | HEPATOCYTE | 1 | | |
| | MUSCLE PRECURSOR CELLS | MUSCLE PRECURSOR CELL | 1 | | |
| | MUSCLE PRECURSORS | MUSCLE PRECURSOR CELLS | 1 | | 1 |
| | MYOGENIC PRECURSORS | | | | 1 |
| file35 | CELL | CELL | 1 | | |

| | | | | | |
|---|---|---|---|---|---|
| | CELLS | CELLS | 1 | | |
| | CELLULAR | EMBRYONIC STEM CELLS | 1 | | 1 |
| | ENDODERM ... CELLS | ENDODERM CELLS | 1 | | |
| | ENDODERM CELLS | EXTRAEMBRYONIC CELLS | 1 | | |
| | EXTRAEMBRYONIC CELLS | GERM CELLS | | 1 | |
| | NEURONAL | NEURONS | 1 | | 1 |
| | NEURONS | PRIMORDIAL GERM CELLS | 1 | | |
| | PCGS | STEM CELLS | 1 | | 1 |
| | PGCS | | | | 1 |
| | PRIMITIVE GERM CELLS ... PCGS | | | | 1 |
| | PRIMORDIAL GERM CELLS | | | | |
| | STEM CELLS | | | | |
| file36 | ASTROCYTES | ASTROCYTES | 1 | | |
| | BIPOLAR CELLS | CELL | 1 | | 1 |
| | CELL | CELLS | 1 | | |
| | CELLS | CONE CELL | 1 | | |
| | CONE | CONE CELLS | 1 | | 1 |
| | CONE CELL | GLIAL CELL | 1 | | |
| | CONE CELLS | PHOTORECEPTOR CELL | 1 | | |
| | CONE PHOTORECEPTORS | PHOTORECEPTOR CELLS | 1 | | 1 |
| | CONES | | | | 1 |
| | GLIAL CELL | | | | |
| | MÃ¼LLER GLIA | | | | 1 |
| | MÃ¼LLER GLIAL | | | | 1 |
| | PHOTORECEPTOR | | | | 1 |
| | PHOTORECEPTOR CELL | | | | |
| | PHOTORECEPTOR CELLS | | | | |
| | PHOTORECEPTORS | | | | 1 |
| | PHOTORECEPTORS IN ... RETINA | | | | 1 |
| | RADIAL GLIA | | | | 1 |
| | ROD ... PHOTORECEPTORS | | | | 1 |
| | ROD CELL | | | | 1 |
| | ROD PHOTORECEPTOR | | | | 1 |
| | ROD PHOTORECEPTOR CELLS | | | | 1 |
| | RODS | | | | 1 |
| file37 | CELL | CELL | 1 | | |
| | OOCYTES | OOCYTES | 1 | | |

| | | | | | |
|---|---|---|---|---|---|
| | CELLS | CELLS | 1 | | |
| file38 | CELL | CELL | 1 | | |
| | CELLS | CELLS | 1 | | |
| | MYOCYTE | MYOCYTE | 1 | | |
| | MYOCYTES | MYOCYTES | 1 | | |
| | SPERM | SPERM | 1 | | |
| | STEM ... CELLS | | | | 1 |
| | STEM CELLS | | | | 1 |
| file39 | CELL | CELL | 1 | | |
| | CELLS | CELLS | 1 | | |
| | CELLULAR | GRANULE CELLS | 1 | | 1 |
| | GRANULE CELLS | NEURONS | 1 | | |
| | NEURONAL | OLFACTORY SENSORY NEURONS | 1 | | 1 |
| | NEURONS | OOCYTES | 1 | | |
| | OLFACTORY SENSORY NEURONS | OUTPUT NEURONS | 1 | | 1 |
| | OOCYTES | PYRAMIDAL CELLS | 1 | | |
| | OUTPUT NEURONS | PYRAMIDAL NEURONS | 1 | | |
| | PRINCIPAL ... NEURONS | SENSORY NEURONS | | 1 | 1 |
| | PRINCIPAL NEURONS | | | | 1 |
| | PYRAMIDAL CELLS | | | | |
| | PYRAMIDAL NEURONS | | | | |
| file40 | CELL | CELL | 1 | | |
| | CELLS | CELLS | 1 | | |
| | CELLULAR | FIBROBLAST | | 1 | 1 |
| | FIBROBLAST CELL | FIBROBLASTS | 1 | | 1 |
| | FIBROBLASTS | GLIAL CELLS | 1 | | |
| | GLIAL CELLS | HEPATOCYTE | 1 | 1 | |
| | HEPATOCYTE | | | | |
| file41 | B CELL | CELL | 1 | | 1 |
| | CELL | CELLS | 1 | | |
| | CELLS | EPIDERMAL CELLS | 1 | | |
| | CELLULAR | EPITHELIAL CELLS | 1 | | 1 |
| | EPIDERMAL CELLS | FIBROBLAST | 1 | | |
| | EPITHELIAL CELLS | FIBROBLASTS | 1 | | |
| | FIBROBLAST | | | | |

| | | | | |
|---|---|---|---|---|
| | FIBROBLASTS | | | |
| file42 | ASTROCYTES | ASTROCYTES | 1 | |
| | CELL | CELL | 1 | |
| | CELLS | CELLS | 1 | |
| | EGGS | EGGS | 1 | |
| | GLIAL | MICROGLIA | 1 | 1 |
| | MICROGLIA | NEURONS | 1 | |
| | MICROGLIAL | | | 1 |
| | NEURONAL | | | 1 |
| | NEURONS | | | |
| file43 | ADIPOCYTE | ADIPOCYTE | 1 | |
| | ADIPOCYTES | ADIPOCYTES | 1 | |
| | CELL | CELL | 1 | |
| | CELLS | CELLS | 1 | |
| | CELLULAR | CHONDROCYTES | 1 | 1 |
| | CHONDROCYTES | FIBROBLAST | 1 | |
| | FIBROBLAST | FIBROBLASTS | 1 | |
| | FIBROBLASTS | GLIAL CELLS | 1 | |
| | GLIAL CELLS | HYPERTROPHIC CHONDROCYTES | 1 | |
| | HYPERTROPHIC CHONDROCYTES | MESENCHYMAL CELL | 1 | |
| | MESENCHYMAL CELL | MESENCHYMAL CELLS | 1 | |
| | MESENCHYMAL CELLS | NEURONS | 1 | |
| | MULTIPOTENTIAL PROGENITOR CELLS | OLIGODENDROCYTES | 1 | 1 |
| | NEURONS | OSTEOBLAST | 1 | |
| | OLIGODENDROCYTES | OSTEOBLASTS | 1 | |
| | OSTEOBLAST | OSTEOCLAST | 1 | |
| | OSTEOBLASTS | OSTEOCLASTS | 1 | |
| | OSTEOCLAST | STEM CELL | 1 | |
| | OSTEOCLASTS | STEM CELLS | 1 | |
| | STEM ... CELLS | STROMAL CELLS | | |
| | STEM CELL | | | |
| | STEM CELLS | | | |
| | STROMAL CELLS | | | |
| file44 | CELL | CELL | 1 | |
| | CELLS | CELLS | 1 | |
| | CELLULAR | ERYTHROCYTES | 1 | 1 |

| | | | | | |
|---|---|---|---|---|---|
| | ERYTHROCYTES | FIBROBLAST | 1 | | |
| | EUKARYOTIC CELLS | FIBROBLASTS | 1 | | 1 |
| | FIBROBLAST | KERATINOCYTES | 1 | | |
| | FIBROBLASTS | OOCYTE | 1 | | |
| | NEURONAL | OOCYTES | 1 | | |
| | OOCYTE | PLATELET | 1 | | |
| | OOCYTES | PLATELETS | 1 | | |
| | PLATELET | SPERM | 1 | | |
| | PLATELETS | | | | |
| | SPERM | | | | |
| | STEMS CELLS | | | | 1 |
| file45 | | | | | |
| file46 | CELL | CELL | 1 | | |
| | CELLS | CELLS | 1 | | |
| | EGGS | EGGS | 1 | | |
| | GLIAL CELL | GRANULE CELLS | | | 1 |
| | GRANULAR CELLS | NEURON | | | 1 |
| | GRANULE CELLS | PYRAMIDAL CELLS | | | |
| | NEURON | | | | |
| | PYRAMIDAL CELLS | | | | |
| | SPERMATOZOA | | | | 1 |
| | STEM ... CELLS | | | | 1 |
| file47 | CELL | CELL | 1 | | |
| | CELLS | CELLS | 1 | | |
| | CELLULAR | GERM CELL | 1 | | 1 |
| | GERM CELL | GERM CELLS | 1 | | |
| | GERM CELLS | GERMLINE STEM CELLS | 1 | | |
| | GERMLINE STEM CELLS | STEM CELL | 1 | | |
| | MULTIPOTENT ... STEM CELLS | STEM CELLS | | 1 | 1 |
| | STEM CELL | | | | |
| file48 | | | | | |
| file49 | AMACRINE | AMACRINE CELLS | 1 | | 1 |
| | AMACRINE CELLS | CELL | 1 | | |
| | BIPOLAR CELLS | CELLS | 1 | | 1 |

| | | | | | |
|---|---|---|---|---|---|
| | CELL | HORIZONTAL CELLS | 1 | | |
| | CELLS | NEURONS | 1 | | |
| | CONE | PHOTORECEPTOR CELL | 1 | | 1 |
| | CONE PHOTORECEPTOR | PHOTORECEPTOR CELLS | 1 | | 1 |
| | CONE PHOTORECEPTORS | PINEALOCYTES | 1 | | 1 |
| | CONES | | | | 1 |
| | GANGLION CELL | | | | 1 |
| | GANGLION CELLS | | | | 1 |
| | HORIZONTAL CELLS | | | | |
| | MÃ¼LLER GLIA | | | | 1 |
| | MULTIPOTENT ... CELLS | | | | 1 |
| | NEURONS | | | | |
| | PHOTORECEPTOR | | | | 1 |
| | PHOTORECEPTOR CELL | | | | |
| | PHOTORECEPTOR CELLS | | | | |
| | PHOTORECEPTOR CELLS IN ... RETINA | | | | 1 |
| | PHOTORECEPTORS | | | | 1 |
| | PHOTORECEPTORS IN ... RETINA | | | | 1 |
| | PINEALOCYTES | | | | |
| | RETINAL PHOTORECEPTOR | | | | 1 |
| | RETINAL PHOTORECEPTOR CELLS | | | | 1 |
| | RETINAL PHOTORECEPTORS | | | | 1 |
| | ROD ... PHOTORECEPTORS | | | | 1 |
| | ROD PHOTORECEPTOR | | | | 1 |
| | ROD PHOTORECEPTORS | | | | 1 |
| | RODS | | | | 1 |
| file50 | CELL | CELL | 1 | | |
| | CELLS | CELLS | 1 | | |
| | CELLULAR | EMBRYONIC STEM CELL | | 1 | 1 |
| | FIBROBLAST | EMBRYONIC STEM CELLS | | 1 | |
| | FIBROBLASTS | FIBROBLAST | 1 | | |
| | HEPATOCYTES | FIBROBLASTS | 1 | | |
| | STEM CELL | HEPATOCYTES | 1 | | |
| | STEM CELLS | STEM CELL | 1 | | |
| | | STEM CELLS | 1 | | |
| file51 | APOPTOTIC CELLS | CELL | 1 | | 1 |
| | CELL | CELLS | 1 | | |

| | | | | | |
|---|---|---|---|---|---|
| | CELL IN ... EPIBLAST ... REGION | LYMPHOCYTES | 1 | | 1 |
| | CELLS | NEURONS | 1 | | |
| | CELLS IN ... EXTRAEMBRYONIC TISSUES | STEM CELL | 1 | | 1 |
| | CELLS OF ... ECTODERM | STEM CELLS | 1 | | 1 |
| | CELLULAR | TROPHOBLAST CELLS | 1 | | 1 |
| | DIPLOID ... CELL | | | | 1 |
| | EXTRAEMBRYONIC CELLS | | | | 1 |
| | LYMPHOCYTES | | | | |
| | NEURONAL | | | | 1 |
| | NEURONS | | | | |
| | RED BLOOD CELLS | | | | 1 |
| | STEM CELL | | | | |
| | STEM CELLS | | | | |
| | TROPHOBLAST ... CELL | | | | |
| | TROPHOBLAST ... CELLS | | | | |
| | TROPHOBLAST CELLS | | | | |
| file52 | PANCREATIC BETA CELL | CELL | | 1 | |
| | | BETA CELL | | 1 | |
| | | PANCREATIC BETA CELL | 1 | | |
| file53 | APOPTOTIC CELL | CELL | 1 | | 1 |
| | APOPTOTIC CELLS | CELLS | 1 | | 1 |
| | BLOOD CELLS | CHONDROCYTE | 1 | | 1 |
| | CELL | ENDOTHELIAL CELLS | 1 | | |
| | CELLS | FIBROBLAST | 1 | | |
| | CELLULAR | FOLLICLE CELLS | 1 | | |
| | CHONDROCYTE | GERM CELL | 1 | | |
| | ENDOTHELIAL ... CELLS | GERM CELLS | 1 | | |
| | ENDOTHELIAL CELL | SERTOLI CELL | 1 | | 1 |
| | FIBROBLAST | SERTOLI CELLS | 1 | | |
| | FOLLICLE CELLS | | | | |
| | GERM CELL | | | | |
| | GERM CELLS | | | | |
| | PLATELET | | | | 1 |
| | POLYGONAL CELLS | | | | 1 |
| | SERTOLI | | | | 1 |
| | SERTOLI CELL | | | | |
| | SERTOLI CELLS | | | | |

| | | | | | |
|---|---|---|---|---|---|
| | SUPPORTING CELL | | | | 1 |
| | SUPPORTING CELLS | | | | 1 |
| | VASCULAR ENDOTHELIAL CELLS | | | | 1 |
| file54 | CELL | CELL | 1 | | |
| | CELLS | CELLS | 1 | | |
| | FIBROBLASTIC CELL | FIBROBLASTS | 1 | | 1 |
| | FIBROBLASTS | | | | |
| | STEM ... CELLS | | | | 1 |
| file55 | AMELOBLASTS | AMELOBLASTS | 1 | | |
| | BASAL CELLS | BASAL CELLS | 1 | | |
| | CELL | CELL | 1 | | |
| | CELLS | CELLS | 1 | | |
| | CELLS OF ... STRATIFIED EPITHELIA | ECTODERMAL CELLS | 1 | | 1 |
| | CELLS OF EPIDERMIS | EPIDERMAL CELL | 1 | | 1 |
| | CELLULAR | EPITHELIAL CELL | 1 | | 1 |
| | ECTODERMAL CELLS | EPITHELIAL CELLS | 1 | | |
| | EPIDERMAL CELL | KERATINOCYTES | 1 | | |
| | EPITHELIAL CELL | LYMPHOCYTES | 1 | | |
| | EPITHELIAL CELLS | MACROPHAGES | 1 | | |
| | KERATINOCYTES | NEUTROPHILS | 1 | | |
| | LYMPHOCYTES | OOCYTES | 1 | | |
| | MACROPHAGES | STEM CELL | 1 | | |
| | NEUTROPHILS | STEM CELLS | 1 | | |
| | OOCYTES | THYMOCYTE | | 1 | |
| | STEM CELL | THYMOCYTES | | 1 | |
| | STEM CELLS | | | | |
| file56 | BLOOD CELLS | BLOOD CELLS | 1 | | |
| | CELL | CELL | 1 | | |
| | CELLS | CELLS | 1 | | |
| | CELLULAR | EMBRYONIC STEM CELL | | 1 | 1 |
| | ENUCLEATING ... CELLS | EMBRYONIC STEM CELLS | | 1 | 1 |
| | FIBROBLASTS | FIBROBLASTS | 1 | | |
| | STEM CELL | STEM CELL | 1 | | |
| | STEM CELLS | STEM CELLS | 1 | | |
| file57 | PLATELET | PLATELET | 1 | | |

| | | | | | |
|---|---|---|---|---|---|
| file58 | CELL | CELL | 1 | | |
| | CELLS | CELLS | 1 | | |
| | CONE | GLIAL CELLS | 1 | | 1 |
| | CONE PHOTORECEPTOR | INTERNEURONS | 1 | | 1 |
| | CONE PHOTORECEPTORS | NEURON | 1 | | 1 |
| | DOPAMINERGIC NEURONS | NEURONS | 1 | | 1 |
| | GANGLION CELL | OOCYTES | 1 | | 1 |
| | GANGLION CELLS | | 1 | | 1 |
| | GANGLION CELLS OF ... RETINA | | | | 1 |
| | GLIA | | | | 1 |
| | GLIAL CELLS | | | | 1 |
| | INTERNEURONS | | | | |
| | NEURON | | | | |
| | NEURONAL | | | | 1 |
| | NEURONS | | | | |
| | OOCYTES | | | | |
| | PHOTORECEPTOR | | | | 1 |
| | PHOTORECEPTOR ... NEURONS OF ... RETINA | | | | 1 |
| | PHOTORECEPTOR NEURONS | | | | 1 |
| | PHOTORECEPTORS | | | | 1 |
| | PHOTOSENSORY NEURON | | | | 1 |
| | PHOTOSENSORY NEURONS | | | | 1 |
| | PHOTOSENSORY NEURONS IN ... RETINA | | | | 1 |
| | PHOTOSENSORY NEURONS OF ... RETINA | | | | 1 |
| | PRIMARY ... NEURON | | | | 1 |
| | PRIMARY ... NEURONS | | | | 1 |
| | RECEPTORAL ... NEURONS | | | | 1 |
| | RECEPTORAL NEURONS | | | | 1 |
| | ROD PHOTORECEPTOR | | | | 1 |
| | ROD PHOTORECEPTOR NEURONAL | | | | 1 |
| | ROD PHOTOSENSORY NEURONS | | | | 1 |
| file59 | APOPTOTIC CELLS | CELL | 1 | | 1 |
| | CELL | CELLS | 1 | | |
| | CELLS | MESENCHYMAL CELL | 1 | | |
| | EPITHELIAL ... CELL | MUSCLE CELL | | 1 | 1 |
| | MESENCHYMAL CELL | MUSCLE CELLS | | 1 | |
| | NC ... CELL | NEURAL CREST CELL | 1 | | 1 |

| | | | | | |
|---|---|---|---|---|---|
| | NCCS | NEURAL CREST CELLS | 1 | | 1 |
| | NEURAL CREST CELL | SMOOTH MUSCLE CELL | 1 | | |
| | NEURAL CREST CELLS | SMOOTH MUSCLE CELLS | 1 | | |
| | SMOOTH MUSCLE CELL | STEM CELLS | 1 | | |
| | SMOOTH MUSCLE CELLS | | | | |
| | STEM CELLS | | | | |
| file60 | CELL | CELL | 1 | | |
| | CELLS | CELLS | 1 | | |
| | CELLULAR | EMBRYONIC STEM CELL | | 1 | 1 |
| | FIBROBLAST | EMBRYONIC STEM CELLS | | 1 | |
| | FIBROBLASTS | FIBROBLAST | 1 | | |
| | PROMYELOCYTIC | FIBROBLASTS | 1 | | 1 |
| | STEM CELL | STEM CELL | 1 | | |
| | STEM CELLS | STEM CELLS | 1 | | |
| file61 | APOPTOTIC CELLS | CELL | 1 | | 1 |
| | CELL | CELLS | 1 | | |
| | CELLS | CHONDROCYTE | 1 | | |
| | CHONDROCYTE | CHONDROCYTES | 1 | | |
| | CHONDROCYTES | FIBROBLAST | 1 | | |
| | FIBROBLAST | HYPERTROPHIC CHONDROCYTES | 1 | | 1 |
| | FIBROBLASTIC | OSTEOBLAST | 1 | | 1 |
| | FIBROBLASTIC CELLS | OSTEOBLASTS | 1 | | |
| | HYPERTROPHIC CHONDROCYTES | OSTEOCLAST | 1 | | |
| | OSTEOBLAST | OSTEOCLASTS | 1 | | |
| | OSTEOBLAST CELLS | OSTEOPROGENITOR CELLS | 1 | | 1 |
| | OSTEOBLASTIC | | | | 1 |
| | OSTEOBLASTS | | | | |
| | OSTEOCLAST | | | | |
| | OSTEOCLASTS | | | | |
| | OSTEOPROGENITOR CELLS | | | | |
| | OSTEOPROGENITORS | | | | |
| file62 | CELLS | CELLS | 1 | | |
| | REGULATORY T CELL | REGULATORY T CELL | 1 | | |
| | T CELL | T CELL | 1 | | |
| | CELL | CELL | 1 | | |
| | BLOOD CELLS | BLOOD CELLS | 1 | | |

| | | | | |
|---|---|---|---|---|
| | LYMPHOCYTES | LYMPHOCYTES | 1 | |
| | LYMPHOCYTE | LYMPHOCYTE | 1 | |
| file63 | CELL | CELL | 1 | |
| | CELLS | CELLS | 1 | |
| | EPITHELIAL CELLS | GLIAL CELL | 1 | 1 |
| | GLIAL CELL | STROMAL CELL | 1 | |
| | STROMAL CELL | STROMAL CELLS | 1 | |
| | STROMAL CELLS | | | |
| file64 | APOPTOTIC CELLS | CELL | 1 | 1 |
| | CELL | CELLS | 1 | |
| | CELLS | GAMETES | 1 | |
| | CELLULAR | GERM CELL | 1 | 1 |
| | GAMETES | GERM CELLS | 1 | |
| | GERM CELL | MALE GERM CELL | 1 | |
| | GERM CELLS | MALE GERM CELLS | 1 | |
| | GERM-CELL | MULTINUCLEATE CELLS | 1 | 1 |
| | GONOCYTES | OOCYTE | 1 | 1 |
| | LEYDIG CELLS | OOCYTES | 1 | 1 |
| | MALE GERM CELL | SERTOLI CELL | 1 | |
| | MALE GERM CELLS | SERTOLI CELLS | 1 | |
| | MULTINUCLEATE CELLS | SPERM | 1 | |
| | MULTINUCLEATED CELLS | SPERMATID | 1 | 1 |
| | OOCYTE | SPERMATIDS | 1 | |
| | OOCYTES | SPERMATOCYTE | 1 | |
| | PRIMARY SPERMATOCYTES | SPERMATOCYTES | 1 | |
| | SERTOLI | ZYGOTE | 1 | 1 |
| | SERTOLI CELL | | | |
| | SERTOLI CELLS | | | |
| | SPERM | | | |
| | SPERMATID | | | |
| | SPERMATIDS | | | |
| | SPERMATOCYTE | | | |
| | SPERMATOCYTES | | | |
| | SPERMATOGONIA | | | 1 |
| | SPERMATOZOA | | | 1 |
| | STEM ... CELLS | | | 1 |
| | ZYGOTE | | | |

| | | | | | |
|---|---|---|---|---|---|
| file65 | LYMPHOCYTES | LYMPHOCYTES | 1 | | |
| | CELLS | CELLS | 1 | | |
| | PURKINJE CELLS | PURKINJE CELLS | 1 | | |
| | | LYMPHOBLASTS | | 1 | |
| file66 | ADIPOCYTES | ADIPOCYTES | 1 | | |
| | AMACRINE | AMACRINE CELL | 1 | | 1 |
| | AMACRINE ... CELLS | AMACRINE CELLS | 1 | | |
| | AMACRINE CELL | AMACRINE NEURONS | 1 | | |
| | AMACRINE CELLS | BIPOLAR NEURONS | 1 | | |
| | AMACRINE NEURONS | CELL | 1 | | |
| | APOPTOTIC CELLS | CELLS | 1 | | 1 |
| | BIPOLAR | CHOLINERGIC NEURONS | 1 | | 1 |
| | BIPOLAR CELL | CNS NEURONS | 1 | | 1 |
| | BIPOLAR CELLS | ENTEROCYTES | 1 | | 1 |
| | BIPOLAR CELLS IN ... RETINA | FIBROBLAST | 1 | | 1 |
| | BIPOLAR NEURONS | FIBROBLASTS | 1 | | |
| | BIPOLARS | INTERNEURONS | 1 | | 1 |
| | CELL | NEURON | 1 | | |
| | CELLS | NEURONS | 1 | | |
| | CHOLINERGIC NEURONS | NULL CELLS | 1 | | |
| | CNS NEURONS | ROD BIPOLAR CELLS | 1 | | |
| | CONE | | | | 1 |
| | CONE BIPOLAR CELLS | | | | 1 |
| | CONES | | | | 1 |
| | ENTEROCYTES | | | | |
| | FIBROBLAST | | | | |
| | FIBROBLASTS | | | | |
| | GANGION CELLS | | | | 1 |
| | GANGLION | | | | 1 |
| | GANGLION ... CELL | | | | 1 |
| | GANGLION ... CELLS | | | | 1 |
| | GANGLION ... CELLS IN ... RETINA | | | | 1 |
| | GANGLION CELL | | | | 1 |
| | GANGLION CELLS | | | | 1 |
| | GANGLION NEURONS | | | | 1 |
| | GLIAL | | | | 1 |
| | HORIZONTAL ... CELLS | | | | 1 |

| | | | | | |
|---|---|---|---|---|---|
| | HORIZONTAL ... NEURONS | | | | 1 |
| | INTERNEURONS | | | | |
| | MÃ¼LLER CELLS | | | | 1 |
| | MÃ¼LLER GLIA CELL | | | | 1 |
| | NEURON | | | | |
| | NEURONAL | | | | 1 |
| | NEURONS | | | | |
| | ON BIPOLAR CELLS | | | | 1 |
| | PHOTORECEPTOR | | | | 1 |
| | PHOTORECEPTORS | | | | 1 |
| | RETINAL ... GANGLION ... CELLS | | | | 1 |
| | RETINAL ... ROD CELLS | | | | 1 |
| | RETINAL BIPOLAR ... CELLS | | | | 1 |
| | ROD ... BIPOLAR CELLS | | | | |
| | ROD ... CELL | | | | 1 |
| | ROD ... CELLS | | | | 1 |
| | ROD ... CELLS IN ... RETINA | | | | 1 |
| | ROD BIPOLAR ... CELLS | | | | |
| | ROD BIPOLAR CELLS | | | | |
| | ROD PHOTORECEPTORS | | | | 1 |
| | RODS | | | | 1 |
| file67 | CELL | CELL | 1 | | |
| | CELLS | CELLS | 1 | | |
| | EGGS | EGGS | 1 | | |
| | GAMETE | EMBRYONIC STEM CELL | | 1 | |
| | GAMETES | GAMETES | 1 | | |
| | GERM CELL | GERM CELL | 1 | | |
| | OOCYTE | OOCYTE | 1 | | |
| | OOCYTES | OOCYTES | 1 | | |
| | SPERM | SPERM | 1 | | |
| | SPERMATIDS | SPERMATIDS | 1 | | |
| | SPERMATOCYTE | SPERMATOCYTE | 1 | | |
| | SPERMATOCYTES | SPERMATOCYTES | 1 | | |
| | SPERMATOGONIA | STEM CELL | 1 | | 1 |
| | SPORE | | | | 1 |
| | STEM CELL | | | | |
| | STEM CELLS | | | | 1 |

| | | | | |
|---|---|---|---|---|
| Total | | 515 | 32 | 340 |
| | Accuracy | 0,94 | | |
| | Recall | 0,60 | | |
| | F-Measure | 0,73 | | |
