## Supplementary_Excel_file_S1 2015_Cell Ontology derived for "OntoContext, a new python package for gene contextualization based on the annotation of biomedical texts"

| Num file | Reference CRAFT 1.0 Annotation | Annotation OntoContext | TP | FP | FN |
|---|---|---|---|---|---|
| file1 | CELLS | CELL | | 1 | |
| | CONE PHOTORECEPTORS | CELLS | 1 | | 1 |
| | CONES | MACROPHAGES | 1 | | 1 |
| | MACROPHAGES | RETINAL GANGLION CELL | 1 | | |
| | RETINAL GANGLION CELL | | | | |
| | ROD ... PHOTORECEPTORS | | | | 1 |
| | RODS | | | | 1 |
| file2 | CELL | CELL | 1 | | |
| | STEM ... CELLS | CELLS | 1 | | 1 |
| | CELLS | | | | |
| file3 | OLIGODENDROCYTE | | | | 1 |
| | CELLULAR | | | | 1 |
| file4 | CELLS | CELLS | 1 | | |
| | CELL | CELL | 1 | | |
| | BLASTOMERES | BLASTOMERES | 1 | | |
| | ENDODERM CELLS | ENDODERM CELLS | 1 | | |
| | STEM CELLS | STEM CELLS | 1 | | |
| | EUKARYOTIC CELLS | EUKARYOTIC CELLS | 1 | | |
| file5 | B LYMPHOCYTES | CELL | 1 | | 1 |
| | CELL | CELLS | 1 | | |
| | CELLS | EPITHELIAL CELL | 1 | | |
| | CELLULAR | EPITHELIAL CELLS | 1 | | 1 |
| | EPITHELIAL CELL | LENS FIBER CELLS | | 1 | |
| | EPITHELIAL CELLS | LYMPHOCYTES | | 1 | |
| | MACROPHAGE | MACROPHAGE | 1 | | |
| | MUSCLE CELL | MUSCLE CELL | 1 | | |
| | OSTEOBLASTS | OSTEOBLASTS | 1 | | |
| file6 | CELL | CELL | 1 | | |
| | CELLS | CELLS | 1 | | |
| | CELLULAR | EAR HAIR CELL | 1 | | 1 |
| | DEITERS' CELLS | HAIR CELL | 1 | | 1 |
| | HAIR CELL | HAIR CELLS | 1 | | |
| | HAIR CELLS | INNER EAR HAIR CELL | 1 | | |

| | |
|---|---|
| Accuracy | 0,78 |
| Recall | 0,60 |
| F-Measure | 0,68 |

| | | | | | |
|---|---|---|---|---|---|
| | INNER ... HAIR CELLS | INNER HAIR CELLS | 1 | | |
| | INNER HAIR CELL | MOTONEURONS | 1 | | |
| | INNER HAIR CELLS | NEURON | 1 | | |
| | MOTONEURONS | NEURONS | 1 | | |
| | NEURON | OUTER HAIR CELL | 1 | | |
| | NEURON AFFERENTS | OUTER HAIR CELLS | 1 | | 1 |
| | NEURONAL | PILLAR CELLS | 1 | | 1 |
| | NEURONS | PRIMARY NEURONS | 1 | | |
| | OUTER HAIR CELL | SENSORY NEURON | 1 | | |
| | OUTER HAIR CELLS | SENSORY NEURONS | 1 | | |
| | PILLAR CELLS | | | | |
| | PRIMARY NEURONS | | | | |
| | SENSORY NEURON | | | | |
| | SENSORY NEURONS | | | | |
| | SUPPORTING CELL | | | | 1 |
| | SUPPORTING CELLS | | | | 1 |
| | VESTIBULAR HAIR CELL | | | | 1 |
| file7 | BASOPHILES | BLOOD CELL | | 1 | 1 |
| | CARDIOMYOCYTES | BLOOD CELLS | | 1 | |
| | CELL | CARDIOMYOCYTES | 1 | | |
| | CELLS | CELL | 1 | | |
| | CELLULAR | CELLS | 1 | | 1 |
| | EOSINOPHILES | ERYTHROCYTE | 1 | | 1 |
| | ERYTHROCYTE | ERYTHROCYTES | 1 | | |
| | ERYTHROCYTES | FRACTION AS | | 1 | |
| | LEUCOCYTES | LEUCOCYTES | 1 | | |
| | LEUKOCYTES | LEUKOCYTES | 1 | | |
| | LYMPHOCYTES | LYMPHOCYTES | 1 | | |
| | MONOCYTES | MONOCYTES | 1 | | |
| | NEUTROPHILES | PLATELET | 1 | | 1 |
| | PLATELET | PLATELETS | 1 | | |
| | PLATELETS | RED BLOOD CELL | 1 | | |
| | RED BLOOD CELL | RED BLOOD CELLS | 1 | | |
| | RED BLOOD CELLS | | | | |
| file8 | CELL | CELL | 1 | | |
| | NEURONAL | | | | 1 |

| | | | | | |
|---|---|---|---|---|---|
| file9 | CELL | CELL | 1 | | |
| | CELLS | CELLS | 1 | | |
| | ERYTHROID CELLS | NEURON | 1 | | 1 |
| | NEURON | NEURONS | 1 | | |
| | NEURONAL | OLFACTORY BULB | | 1 | 1 |
| | NEURONS | SPERM | 1 | | |
| | OLFACTORY NEURON | SPERMATID | 1 | | 1 |
| | OLFACTORY NEURONS | T-CELL | 1 | | 1 |
| | SPERM | | | | |
| | SPERMATID | | | | |
| | T-CELL | | | | |
| file10 | ASTROCYTES | ASTROCYTES | 1 | | |
| | CELL | CELL | 1 | | |
| | CELLS | CELLS | 1 | | |
| | CELLULAR | HIPPOCAMPAL NEURONS | | 1 | 1 |
| | NEURON | HIPPOCAMPUS | | 1 | |
| | NEURONAL CELL | NEURON | 1 | | 1 |
| | NEURONAL CELLS | NEURONS | 1 | | 1 |
| | NEURONS | | | | |
| file11 | CELL | CELL | 1 | | |
| | CELLS | CELLS | 1 | | |
| | CELLULAR | EMBRYONIC CELLS | | 1 | 1 |
| | DERMAL PAPILLA CELLS | KERATINOCYTES | 1 | | 1 |
| | DERMAL PAPILLAE CELLS | MELANOBLAST | 1 | | 1 |
| | KERATINOCYTES | MELANOBLASTS | 1 | | |
| | MELANOBLAST | MELANOCYTE | 1 | | 1 |
| | MELANOBLASTS | MELANOCYTES | 1 | | 1 |
| | MELANOCYTE | MESENCHYMAL CELLS | 1 | | |
| | MELANOCYTES | PIGMENT CELL | 1 | | |
| | MESENCHYMAL CELLS | PIGMENT CELLS | 1 | | |
| | PIGMENT CELL | PRECURSOR CELLS | | 1 | |
| | PIGMENT CELLS | | | | |
| | PIGMENT-CELL | | | | 1 |
| | STEM CELLS | | | | 1 |
| file12 | CELL | ALVEOLAR EPITHELIAL CELLS | | 1 | |
| | CELLS | CELL | 1 | | |

| | | | | | |
|---|---|---|---|---|---|
| | CELLS IN ... EPITHELIAL ... REGIONS | CELLS | 1 | | 1 |
| | CELLS IN ... MESENCHYMAL REGIONS | EMBRYONIC FIBROBLASTS | | 1 | 1 |
| | CELLULAR | ENDOTHELIAL CELL | 1 | | 1 |
| | ENDOTHELIAL CELL | EPITHELIAL CELLS | 1 | | |
| | EPITHELIAL ... CELLS | FIBROBLASTS | 1 | | |
| | EPITHELIAL CELLS | LUNG EPITHELIAL CELL | 1 | | |
| | EPITHELIAL CELLS ... IN ... LUNGS | LUNG EPITHELIAL CELLS | 1 | | 1 |
| | EPITHELIAL CELLS IN ... LUNGS | MESENCHYME CELLS | 1 | | |
| | FIBROBLASTS | PLATELET | 1 | | |
| | LUNG EPITHELIAL CELL | TYPE I ALVEOLAR EPITHELIAL CELLS | | 1 | |
| | LUNG EPITHELIAL CELLS | TYPE II ALVEOLAR EPITHELIAL CELLS | | 1 | |
| | MESENCHYME CELLS | | | | |
| | PLATELET | | | | |
| file13 | CELL | CELL | 1 | | |
| | CELLS | CELLS | 1 | | |
| | CELLULAR | ENTEROCYTES | 1 | | 1 |
| | ENTEROCYTES | EPITHELIAL CELLS | 1 | | |
| | EPITHELIAL CELLS | FIBROBLASTS | 1 | | |
| | FIBROBLASTS | | | | |
| file14 | CELLS | CELL | | 1 | |
| | CELLULAR | CELLS | 1 | | 1 |
| | DOPAMINE CELLS | DOPAMINERGIC CELL | 1 | 1 | 1 |
| | DOPAMINERGIC CELL | DOPAMINERGIC CELLS | 1 | | 1 |
| | DOPAMINERGIC CELLS | MITRAL CELLS | | 1 | |
| | NEURONAL | OLFACTORY BULB | | 1 | 1 |
| file15 | CELL | OOCYTE | 1 | | |
| | OOCYTE | CELLS | 1 | | |
| | CELLS | CELL | 1 | | |
| | CELLULAR | | | | 1 |
| file16 | APOPTOTIC CELLS | BLOOD CELL | 1 | | 1 |
| | BLOOD CELL | BLOOD CELLS | 1 | | |
| | BLOOD CELLS | CELL | 1 | | |
| | CELL | CELLS | 1 | | |
| | CELLS | CIRCULATING CELLS | 1 | | |
| | CIRCULATING CELLS | EMBRYONIC FIBROBLAST | | 1 | |

| | | | | | |
|---|---|---|---|---|---|
| | EGGS | EMBRYONIC FIBROBLASTS | | 1 | 1 |
| | ERYTHROCYTES | ERYTHROCYTES | 1 | | |
| | ERYTHROID | ERYTHROID PROGENITOR CELLS | 1 | | 1 |
| | ERYTHROID CELL | FIBROBLAST | 1 | | 1 |
| | ERYTHROID CELLS | FIBROBLASTS | 1 | | 1 |
| | ERYTHROID PROGENITOR CELLS | HEMANGIOBLASTS | | 1 | |
| | ERYTHROID PROGENITORS | HEMATOPOIETIC CELLS | 1 | | 1 |
| | FIBROBLAST | HEMATOPOIETIC STEM CELL | 1 | | |
| | FIBROBLAST CELL | HEMATOPOIETIC STEM CELLS | 1 | | 1 |
| | FIBROBLASTS | LYMPHOCYTES | 1 | | |
| | HEMATOPOIETIC CELLS | MACROPHAGES | 1 | | |
| | HEMATOPOIETIC PROGENITOR CELLS | MESENCHYMAL CELLS | 1 | | 1 |
| | HEMATOPOIETIC PROGENITORS | MESODERMAL CELLS | 1 | | 1 |
| | HEMATOPOIETIC STEM CELL | MYELOID CELLS | 1 | | |
| | HEMATOPOIETIC STEM CELLS | PRIMITIVE ERYTHROID CELLS | | 1 | |
| | LYMPHOCYTES | PROERYTHROBLAST | 1 | | |
| | MACROPHAGES | PROERYTHROBLASTS | 1 | | |
| | MESENCHYMAL CELLS | RBCS | 1 | | |
| | MESODERMAL CELLS | RED BLOOD CELL | 1 | | |
| | MYELOID | RED BLOOD CELLS | 1 | | 1 |
| | MYELOID CELLS | RETICULOCYTE | 1 | | |
| | PROERYTHROBLAST | STEM CELL | 1 | | |
| | PROERYTHROBLASTS | STEM CELLS | 1 | | |
| | RBCS | T-CELLS | 1 | | |
| | RED ... BLOOD CELLS | WHITE BLOOD CELLS | 1 | | |
| | RED BLOOD CELL | | | | |
| | RED BLOOD CELLS | | | | |
| | RETICULOCYTE | | | | |
| | STEM ... CELLS | | | | |
| | STEM CELL | | | | |
| | STEM CELLS | | | | |
| | T-CELLS | | | | |
| | WHITE BLOOD CELLS | | | | |
| file17 | APOPTOTIC CELLS | APCS | | 1 | 1 |
| | CELL | CELL | 1 | | |
| | CELLS | CELLS | 1 | | |
| | CELLS IN VIVO | EMBRYONIC STEM CELLS | | 1 | 1 |
| | STEM CELLS | STEM CELLS | 1 | | |

| | | | | | |
|---|---|---|---|---|---|
| file18 | BASAL CELLS | BASAL CELL | | 1 | |
| | CELL | BASAL CELLS | 1 | | |
| | CELLS | CELL | 1 | | |
| | EPITHELIAL CELLS | CELLS | 1 | | |
| | FIBROCYTES | EPITHELIAL CELLS | 1 | | |
| | HAIR CELL | FIBROCYTES | 1 | | |
| | HAIR CELLS | HAIR CELL | 1 | | |
| | STEM CELL | HAIR CELLS | 1 | | |
| | TRANSITIONAL CELLS | SENSORY HAIR CELLS | | 1 | 1 |
| | VESTIBULAR DARK CELLS | STEM CELL | 1 | | |
| | VESTIBULAR HAIR CELLS | STRIAL INTERMEDIATE CELLS | | 1 | 1 |
| | | STRIAL MARGINAL CELLS | | 1 | |
| | | VESTIBULAR DARK CELLS | 1 | | |
| file19 | ADIPOCYTE | ADIPOCYTE | 1 | | |
| | CELL | CELL | 1 | | |
| | CELLULAR | MOTONEURON | 1 | | 1 |
| | MOTONEURON | MOTOR NEURON | 1 | | |
| | MOTOR NEURON | MUSCLE FIBER | | 1 | |
| | ZYGOTES | MUSCLE FIBERS | | 1 | |
| | | NEURON | | 1 | |
| | | TYPE I MUSCLE FIBER | | 1 | |
| | | ZYGOTES | 1 | | |
| file20 | APOPTOTIC ... CELLS | ALVEOLAR EPITHELIAL CELLS | | 1 | 1 |
| | APOPTOTIC CELL | CELL | 1 | | 1 |
| | APOPTOTIC CELLS | CELLS | 1 | | 1 |
| | CELL | EARLY ERYTHROBLAST | 1 | | |
| | CELLS | EPITHELIAL CELLS | 1 | | |
| | CELLS IN ... MESENCHYME | ERYTHROBLAST | | 1 | 1 |
| | CELLULAR | ERYTHROCYTES | 1 | | 1 |
| | EARLY ERYTHROBLAST | FIBROBLASTS | 1 | | 1 |
| | EPITHELIAL CELLS | MACROPHAGE | 1 | | |
| | ERYTHROCYTES | MACROPHAGES | 1 | | |
| | FIBROBLASTS | MEGAKARYOCYTES | 1 | | |
| | HEMATOPOIETIC STEM CELLS | MONOCYTE | 1 | | 1 |
| | MACROPHAGE | NEUTROPHILS | 1 | | |
| | MACROPHAGES | PERITONEAL MACROPHAGES | 1 | | |

| | | | | | |
|---|---|---|---|---|---|
| | MEGAKARYOCYTES | PHAGOCYTES | 1 | | |
| | MESENCHYMAL ... CELLS | PHOTORECEPTOR CELLS | 1 | | 1 |
| | MONOCYTE | PIGMENTED EPITHELIAL CELLS | 1 | | |
| | NEURONAL | PRIMARY LENS FIBER | | 1 | 1 |
| | NEURONAL CELLS | RETINAL CELL | | 1 | 1 |
| | NEUTROPHILS | THYMOCYTES | | 1 | |
| | PERITONEAL MACROPHAGES | TYPE II ALVEOLAR EPITHELIAL CELLS | | 1 | |
| | PHAGOCYTES | | | | |
| | PHOTORECEPTOR CELLS | | | | |
| | PIGMENTED CELLS | | | | 1 |
| | PIGMENTED EPITHELIAL CELLS | | | | |
| | PLATELET | | | | 1 |
| | STEM ... CELLS | | | | 1 |
| file21 | CARTILAGE CELLS | CELL | 1 | | 1 |
| | CELL | CELLS | 1 | | |
| | CELLS | CHONDROCYTES | 1 | | |
| | CELLULAR | MACROPHAGES | 1 | | 1 |
| | CHONDROCYTES | MONONUCLEAR CELLS | 1 | | |
| | EGGS | NEUTROPHILS | 1 | | 1 |
| | FIBROBLAST | OSTEOBLASTS | 1 | | 1 |
| | HYPERTROPHIC CHONDROCYTES | SYNOVIAL CELLS | 1 | | 1 |
| | MACROPHAGES | SYNOVIOCYTES | 1 | | |
| | MESENCHYMAL CELLS | | | | 1 |
| | MONONUCLEAR CELLS | | | | |
| | NEUTROPHILS | | | | |
| | OSTEOBLASTS | | | | |
| | SYNOVIAL CELLS | | | | |
| | SYNOVIOCYTES | | | | |
| file22 | CELL | CELL | 1 | | |
| | CELLS | CELLS | 1 | | |
| | GLIA | MOTOR NEURON | 1 | | 1 |
| | MOTOR NEURON | MOTOR NEURONS | 1 | | |
| | MOTOR NEURONS | MYOTUBES | | 1 | |
| | NEURON | NEURON | 1 | | |
| | NEURONS | NEURONS | 1 | | |
| | SENSORY NEURONS | PROPRIOCEPTIVE NEURON | | 1 | |
| | | PROPRIOCEPTIVE NEURONS | | 1 | |

| | | | | | |
|---|---|---|---|---|---|
| | | SENSORY NEURONS | 1 | | |
| file23 | CELL | CELL | 1 | | |
| | CELLS | CELLS | 1 | | |
| | CELLULAR | NEURON | 1 | | 1 |
| | CNS NEURONS | NEURONS | 1 | | 1 |
| | GRANULE NEURON | PRECURSOR CELL | | 1 | 1 |
| | GRANULE NEURONS | PRECURSOR CELLS | | 1 | 1 |
| | NEURON | PURKINJE NEURON | 1 | | |
| | NEURONAL | | 1 | | 1 |
| | NEURONAL CELL | | | | 1 |
| | NEURONAL CELLS | | | | 1 |
| | NEURONS | | | | |
| | PURKINJE NEURON | | | | |
| file24 | BASAL ... CELLS | CELL | 1 | | 1 |
| | CELL | CELLS | 1 | | |
| | CELLS | EPIDERMAL CELLS | 1 | | |
| | CELLS OF ... ECTODERM | EPITHELIAL CELLS | 1 | | 1 |
| | CELLS WITHIN ... EPIDERMIS | FIBROBLAST | 1 | | 1 |
| | CELLULAR | HAIR CELL | 1 | | 1 |
| | ECTODERMAL CELLS | HEPATOCYTES | 1 | | 1 |
| | EPIDERMAL CELLS | KERATINOCYTE | 1 | | |
| | EPITHELIAL ... CELL | KERATINOCYTES | 1 | | 1 |
| | EPITHELIAL CELLS | MESENCHYMAL CELLS | 1 | | |
| | FIBROBLAST | MULTIPOTENT CELLS | | 1 | |
| | GRANULAR LAYER CELLS | T CELL | 1 | | 1 |
| | HAIR CELL | | | | |
| | HEPATOCYTES | | | | |
| | KERATINOCYTE | | | | |
| | KERATINOCYTES | | | | |
| | MESENCHYMAL CELLS | | | | |
| | MOTILE ... CELLS | | | | 1 |
| | SQUAMOUS CELL | | | | 1 |
| | T CELL | | | | |
| file25 | BIPOLAR CELL | CELL | 1 | | 1 |
| | BIPOLAR CELLS | CELLS | 1 | | 1 |
| | CELL | HORIZONTAL CELLS | 1 | | |

| | | | | | |
|---|---|---|---|---|---|
| | CELLS | MüLLER CELLS | 1 | | |
| | CONE | NEURONS | 1 | | 1 |
| | HORIZONTAL ... CELL | PHOTORECEPTOR CELLS | 1 | | 1 |
| | HORIZONTAL ... CELLS | ROD BIPOLAR CELLS | 1 | | |
| | HORIZONTAL CELL | SENSORY NEURONS | 1 | | 1 |
| | HORIZONTAL CELLS | | | | |
| | MÃ¼LLER CELLS | | | | |
| | NEURONAL | | | | 1 |
| | NEURONS | | | | |
| | OFF ... BIPOLAR CELLS | | | | 1 |
| | ON ... BIPOLAR CELLS | | | | 1 |
| | PHOTORECEPTOR | | | | 1 |
| | PHOTORECEPTOR CELLS | | | | |
| | PHOTORECEPTORS | | | | 1 |
| | PHOTORECEPTORS ... R6 | | | | 1 |
| | PHOTORECEPTORS R1 | | | | 1 |
| | R1 ... PHOTORECEPTORS | | | | 1 |
| | R6 PHOTORECEPTORS | | | | 1 |
| | ROD BIPOLAR | | | | 1 |
| | ROD BIPOLAR CELLS | | | | |
| | ROD PHOTORECEPTORS | | | | 1 |
| | RODS | | | | 1 |
| | SENSORY NEURONS | | | | |
| file26 | ASTROCYTES | ASTROCYTES | 1 | | |
| | ASTROCYTIC | CARDIAC MYOCYTES | 1 | | 1 |
| | CARDIAC MYOCYTES | CELL | 1 | | |
| | CELL | CELLS | 1 | | |
| | CELLS | EMBRYONIC STEM CELLS | | 1 | |
| | CELLULAR | GRANULE CELL | 1 | | 1 |
| | GLIAL | HEPATOCYTE | 1 | | 1 |
| | GRANULE CELL | HEPATOCYTES | 1 | | |
| | HEPATOCYTE | HIPPOCAMPUS | | 1 | |
| | HEPATOCYTES | MUSCLE FIBERS | | 1 | |
| | LYMPHOCYTIC | MYOCYTES | | 1 | 1 |
| | NEURON | NEURON | 1 | | |
| | NEURONAL | NEURONS | 1 | | 1 |
| | NEURONS | OOCYTES | 1 | | |
| | OOCYTES | PYRAMIDAL NEURONS | 1 | | |

| | | | | | |
|---|---|---|---|---|---|
| | PHAGOCYTIC CELLS | STEM CELLS | 1 | | 1 |
| | PURKINJE ... CELL | | | | 1 |
| | PYRAMIDAL NEURONS | | | | |
| | STEM CELLS | | | | |
| file27 | ASTROCYTE | ADRENAL CHROMAFFIN CELLS | | 1 | 1 |
| | ASTROCYTES | ASTROCYTES | 1 | | |
| | ASTROCYTIC | BLOOD CELLS | | 1 | 1 |
| | ASTROCYTIC CELL | CELL | 1 | | 1 |
| | ASTROGLIAL ... CELL | CELLS | 1 | | 1 |
| | ASTROGLIAL CELL | CHROMAFFIN CELLS | 1 | | 1 |
| | ASTROGLIAL CELLS | CULTURED CELLS | | 1 | 1 |
| | CELL | ENDODERM CELLS | 1 | | |
| | CELLS | FIBROBLASTS | 1 | | |
| | CELLS OF ... STRATUM GRANULOSUM | FORESKIN FIBROBLASTS | | 1 | 1 |
| | CELLULAR | GLIAL CELL | 1 | | 1 |
| | CHROMAFFIN CELLS | GLIAL CELLS | 1 | | |
| | ENDODERM CELLS | HIPPOCAMPUS | | 1 | |
| | ENDODERMAL ... CELLS | MESODERMAL CELLS | 1 | | 1 |
| | FIBROBLASTS | MYOBLAST | 1 | | |
| | GLIAL | NEURON | 1 | | 1 |
| | GLIAL CELL | NEURONS | 1 | | |
| | GLIAL CELLS | PLATELET | 1 | | |
| | MESODERMAL CELLS | PURKINJE CELLS | 1 | | |
| | MYOBLAST | PYRAMIDAL NEURON | 1 | | |
| | NEURON | PYRAMIDAL NEURONS | 1 | | |
| | NEURONAL | RADIAL GLIAL CELLS | 1 | | 1 |
| | NEURONAL ... CELL | RED BLOOD CELLS | 1 | | 1 |
| | NEURONAL ... CELLS | | | | 1 |
| | NEURONAL CELLS | | | | 1 |
| | NEURONS | | | | |
| | PLATELET | | | | |
| | PURKINJE CELLS | | | | |
| | PURKINJE-CELL | | | | 1 |
| | PURKINJE-CELLS | | | | 1 |
| | PYRAMIDAL NEURON | | | | |
| | PYRAMIDAL NEURONS | | | | |
| | RADIAL GLIAL CELLS | | | | |
| | RED BLOOD CELLS | | | | |

| | | | | | |
|---|---|---|---|---|---|
| file28 | AFFERENT CELL | CELL | 1 | | 1 |
| | AFFERENT SENSORY NEURONS | CELLS | 1 | | 1 |
| | AFFERENTS ... NEURONS | MOTOR NEURON | 1 | | 1 |
| | APOPTOTIC CELLS | MOTOR NEURONS | 1 | | 1 |
| | CELL | MUSCLE FIBERS | | 1 | |
| | CELLS | NEUROBLAST | 1 | | |
| | CELLULAR | NEUROBLASTS | 1 | | 1 |
| | MOTOR NEURON | NEURON | 1 | | |
| | MOTOR NEURONS | NEURONS | 1 | | |
| | NEUROBLAST | PEPTIDERGIC NEURONS | 1 | | |
| | NEUROBLASTS | PROPRIOCEPTIVE NEURONS | | 1 | |
| | NEURON | SENSORY NEURON | 1 | | |
| | NEURONAL | SENSORY NEURONS | 1 | | 1 |
| | NEURONAL CELL | | | | 1 |
| | NEURONS | | | | |
| | PEPTIDERGIC NEURONS | | | | |
| | SENSORY ... NEURONS | | | | |
| | SENSORY NEURON | | | | |
| | SENSORY NEURONS | | | | |
| file29 | CELL | CELL | 1 | | |
| | CELLS | CELLS | 1 | | |
| | EGGS | CEREBELLAR GRANULE CELLS | | 1 | 1 |
| | GRANULE CELL | GRANULE CELL | 1 | | |
| | GRANULE CELLS | GRANULE CELLS | 1 | | |
| | NEURON | HIPPOCAMPUS | | 1 | 1 |
| | NEURONAL | MYELINATING SCHWANN CELLS | | 1 | 1 |
| | NEURONAL CELL | NEURONS | 1 | | 1 |
| | NEURONS | PURKINJE CELL | 1 | | |
| | PURKINJE CELL | SCHWANN CELL | 1 | | |
| | SCHWANN CELL | SCHWANN CELLS | 1 | | |
| | SCHWANN CELLS | | | | |
| | SCHWANN-CELL | | | | 1 |
| | SPERMATOZOA | | | | 1 |
| | STEM ... CELLS | | | | 1 |
| file30 | CELL | CELLS | 1 | | |
| | CELLS | CELL | 1 | | |

| | | | | | |
|---|---|---|---|---|---|
| file31 | BLOOD CELL | BLOOD CELL | 1 | | |
| | CELL | CELL | 1 | | |
| | CELLS | CELLS | 1 | | |
| | ENDOTHELIAL CELL | ENDOTHELIAL CELL | 1 | | |
| | EPITHELIAL CELLS | EPITHELIAL CELLS | 1 | | |
| | GOBLET CELL | GOBLET CELL | 1 | | |
| | GOBLET CELLS | GOBLET CELLS | 1 | | |
| | HEMATOPOIETIC CELL | IMMUNE CELL | 1 | | 1 |
| | IMMUNE CELL | IMMUNE CELLS | 1 | | |
| | IMMUNE CELLS | LYMPHOCYTE | 1 | | |
| | LYMPHOCYTE | LYMPHOCYTES | 1 | | |
| | LYMPHOCYTES | MAST CELL | 1 | | |
| | MAST CELL | MAST CELLS | 1 | | |
| | MAST CELLS | MONONUCLEAR CELL | 1 | | |
| | MONONUCLEAR CELL | NEUTROPHIL | 1 | | |
| | NEUTROPHIL | NEUTROPHILS | 1 | | |
| | NEUTROPHILS | PANETH CELL | 1 | | |
| | PANETH CELL | STEM CELL | 1 | | |
| | STEM CELL | T CELL | 1 | | |
| | T CELL | | | | |
| file32 | CELLS | CELL | | 1 | |
| | CELLS ... OF ... EPITHELIUM | CELLS | 1 | | 1 |
| | CELLULAR | NATURAL KILLER CELL | 1 | | 1 |
| | NATURAL KILLER CELL | RECEPTOR CELLS | | 1 | |
| | TASTE CELLS | TASTE RECEPTOR CELLS | 1 | | 1 |
| | TASTE RECEPTOR CELLS | | | | |
| | TRCS | | | | 1 |
| file33 | CELL | CELL | 1 | | |
| | CELLS | CELLS | 1 | | |
| | GLIA | HIPPOCAMPUS | | 1 | 1 |
| | NEURONAL | NEURAL CELL | | 1 | 1 |
| | NEURONES | NEURONS | 1 | | 1 |
| | NEURONS | OLFACTORY BULBS | | 1 | |
| file34 | CELLS | CELL | | 1 | |
| | HEPATOCYTE | CELLS | 1 | | |

| | | | | | |
|---|---|---|---|---|---|
| | MUSCLE PRECURSOR CELL | HEPATOCYTE | 1 | | |
| | MUSCLE PRECURSOR CELLS | MUSCLE FIBERS | | 1 | |
| | MUSCLE PRECURSORS | MUSCLE PRECURSOR CELL | 1 | | 1 |
| | MYOGENIC PRECURSORS | MUSCLE PRECURSOR CELLS | 1 | | 1 |
| | | MYOTUBULES | | 1 | |
| | | PRECURSOR CELL | | 1 | |
| | | PRECURSOR CELLS | | 1 | |
| file35 | CELL | CELL | 1 | | |
| | CELLS | CELLS | 1 | | |
| | CELLULAR | EMBRYONIC STEM CELLS | | 1 | 1 |
| | ENDODERM ... CELLS | ENDODERM CELLS | 1 | | |
| | ENDODERM CELLS | EXTRAEMBRYONIC CELLS | 1 | | |
| | EXTRAEMBRYONIC CELLS | GERM CELLS | | 1 | |
| | NEURONAL | NEURONS | 1 | | 1 |
| | NEURONS | PRIMORDIAL GERM CELLS | 1 | | |
| | PCGS | STEM CELLS | 1 | | 1 |
| | PGCS | | | | 1 |
| | PRIMITIVE GERM CELLS ... PCGS | | | | 1 |
| | PRIMORDIAL GERM CELLS | | | | |
| | STEM CELLS | | | | |
| file36 | ASTROCYTES | ASTROCYTES | 1 | | |
| | BIPOLAR CELLS | CELL | 1 | | 1 |
| | CELL | CELLS | 1 | | |
| | CELLS | CONE CELL | 1 | | |
| | CONE | CONE CELLS | 1 | | 1 |
| | CONE CELL | ESCS | | 1 | |
| | CONE CELLS | GLIAL CELL | 1 | | |
| | CONE PHOTORECEPTORS | IMMATURE ASTROCYTES | | 1 | 1 |
| | CONES | PHOTORECEPTOR CELL | 1 | | 1 |
| | GLIAL CELL | PHOTORECEPTOR CELLS | 1 | | |
| | MÃ¼LLER GLIA | RETINAL CELLS | | 1 | 1 |
| | MÃ¼LLER GLIAL | | | | 1 |
| | PHOTORECEPTOR | | | | 1 |
| | PHOTORECEPTOR CELL | | | | |
| | PHOTORECEPTOR CELLS | | | | |
| | PHOTORECEPTORS | | | | 1 |
| | PHOTORECEPTORS IN ... RETINA | | | | 1 |

| | | | | | |
|---|---|---|---|---|---|
| | RADIAL GLIA | | | | 1 |
| | ROD ... PHOTORECEPTORS | | | | 1 |
| | ROD CELL | | | | 1 |
| | ROD PHOTORECEPTOR | | | | 1 |
| | ROD PHOTORECEPTOR CELLS | | | | 1 |
| | RODS | | | | 1 |
| file37 | CELL | CELL | 1 | | |
| | OOCYTES | CELLS | 1 | | |
| | CELLS | KIDNEY CELL | | 1 | |
| | | KIDNEY CELLS | | 1 | |
| | | OOCYTES | 1 | | |
| file38 | CELL | CELL | 1 | | |
| | CELLS | CELLS | 1 | | |
| | MYOCYTE | MYOCYTE | 1 | | |
| | MYOCYTES | MYOCYTES | 1 | | |
| | SPERM | SPERM | 1 | | |
| | STEM ... CELLS | | | | |
| | STEM CELLS | | | | 1 |
| file39 | CELL | CELL | 1 | | |
| | CELLS | CELLS | 1 | | |
| | CELLULAR | CORTICAL NEURONS | | 1 | 1 |
| | GRANULE CELLS | FOREBRAIN NEURONS | | 1 | |
| | NEURONAL | GRANULE CELLS | 1 | | 1 |
| | NEURONS | HIPPOCAMPAL NEURONS | | 1 | |
| | OLFACTORY SENSORY NEURONS | HIPPOCAMPAL PYRAMIDAL NEURONS | | 1 | 1 |
| | OOCYTES | HIPPOCAMPUS | | 1 | |
| | OUTPUT NEURONS | MITRAL CELLS | | 1 | |
| | PRINCIPAL ... NEURONS | NEURONS | 1 | | 1 |
| | PRINCIPAL NEURONS | OLFACTORY BULB | | 1 | |
| | PYRAMIDAL CELLS | OLFACTORY SENSORY NEURONS | 1 | | |
| | PYRAMIDAL NEURONS | OOCYTES | 1 | | |
| | | OUTPUT NEURONS | 1 | | |
| | | PYRAMIDAL CELLS | 1 | | |
| | | PYRAMIDAL NEURONS | 1 | | |
| | | SENSORY NEURONS | | 1 | |

| | | | | | |
|---|---|---|---|---|---|
| file40 | CELL | CELL | 1 | | |
| | CELLS | CELLS | 1 | | |
| | CELLULAR | CULTURED CELLS | | 1 | 1 |
| | FIBROBLAST CELL | EMBRYONIC FIBROBLAST | | 1 | 1 |
| | FIBROBLASTS | EMBRYONIC FIBROBLASTS | | 1 | |
| | GLIAL CELLS | FIBROBLAST | 1 | | |
| | HEPATOCYTE | FIBROBLASTS | 1 | | |
| | | GLIAL CELLS | 1 | | |
| | | HEPATOCYTE | 1 | | |
| file41 | B CELL | BRONCHIAL EPITHELIAL CELLS | | 1 | 1 |
| | CELL | CELL | 1 | | |
| | CELLS | CELLS | 1 | | |
| | CELLULAR | EPIDERMAL CELLS | 1 | | 1 |
| | EPIDERMAL CELLS | EPITHELIAL CELLS | 1 | | |
| | EPITHELIAL CELLS | FIBROBLAST | 1 | | |
| | FIBROBLAST | FIBROBLASTS | 1 | | |
| | FIBROBLASTS | | | | |
| file42 | ASTROCYTES | ACTIVATED ASTROCYTES | | 1 | |
| | CELL | ACTIVATED MICROGLIA | | 1 | |
| | CELLS | ASTROCYTES | 1 | | |
| | EGGS | CELL | 1 | | 1 |
| | GLIAL | CELLS | 1 | | 1 |
| | MICROGLIA | CORTICAL NEURONS | | 1 | |
| | MICROGLIAL | HIPPOCAMPUS | | 1 | |
| | NEURONAL | MICROGLIA | 1 | | 1 |
| | NEURONS | NEURONS | 1 | | |
| file43 | ADIPOCYTE | ADIPOCYTE | 1 | | |
| | ADIPOCYTES | ADIPOCYTES | 1 | | |
| | CELL | BONE CELL | | 1 | |
| | CELLS | BONE CELLS | | 1 | |
| | CELLULAR | BONE MARROW STROMAL CELLS | | 1 | 1 |
| | CHONDROCYTES | CELL | 1 | | |
| | FIBROBLAST | CELLS | 1 | | |
| | FIBROBLASTS | CFU-F | | 1 | |
| | GLIAL CELLS | CHONDROCYTES | 1 | | |
| | HYPERTROPHIC CHONDROCYTES | EMBRYONIC CELL | | 1 | |

| | | | | | |
|---|---|---|---|---|---|
| | MESENCHYMAL CELL | EMBRYONIC FIBROBLAST | | 1 | |
| | MESENCHYMAL CELLS | FIBROBLAST | 1 | | |
| | MULTIPOTENTIAL PROGENITOR CELLS | FIBROBLASTS | 1 | | 1 |
| | NEURONS | GLIAL CELLS | 1 | | |
| | OLIGODENDROCYTES | HYPERTROPHIC CHONDROCYTES | 1 | | |
| | OSTEOBLAST | MESENCHYMAL CELL | 1 | | |
| | OSTEOBLASTS | MESENCHYMAL CELLS | 1 | | |
| | OSTEOCLAST | MESENCHYMAL PRECURSOR CELL | | 1 | |
| | OSTEOCLASTS | MESENCHYMAL STEM CELL | 1 | | |
| | STEM ... CELLS | NEURONS | 1 | | |
| | STEM CELL | OLIGODENDROCYTES | 1 | | |
| | STEM CELLS | OSTEOBLAST | 1 | | |
| | STROMAL CELLS | OSTEOBLASTS | 1 | | |
| | | OSTEOCLAST | 1 | | |
| | | OSTEOCLASTS | 1 | | |
| | | PRECURSOR CELL | | 1 | |
| | | STEM CELL | 1 | | |
| | | STEM CELLS | 1 | | |
| | | STROMAL CELLS | 1 | | |
| file44 | CELL | CELL | 1 | | |
| | CELLS | CELLS | 1 | | |
| | CELLULAR | EMBRYONIC FIBROBLASTS | | 1 | 1 |
| | ERYTHROCYTES | ERYTHROCYTES | 1 | | |
| | EUKARYOTIC CELLS | FIBROBLAST | 1 | | 1 |
| | FIBROBLAST | FIBROBLASTS | 1 | | |
| | FIBROBLASTS | KERATINOCYTES | | 1 | |
| | NEURONAL | OOCYTE | 1 | | 1 |
| | OOCYTE | OOCYTES | 1 | | |
| | OOCYTES | PLATELET | 1 | | |
| | PLATELET | PLATELETS | 1 | | |
| | PLATELETS | SPERM | 1 | | |
| | SPERM | | | | |
| | STEMS CELLS | | | | 1 |
| file45 | | | | | |
| file46 | CELL | CELL | 1 | | |
| | CELLS | CELLS | 1 | | |

| | | | | | |
|---|---|---|---|---|---|
| | EGGS | CEREBELLAR GRANULE CELLS | | 1 | 1 |
| | GLIAL CELL | GRANULE CELLS | 1 | | 1 |
| | GRANULAR CELLS | HIPPOCAMPUS | | 1 | 1 |
| | GRANULE CELLS | NEURAL CELL | | 1 | |
| | NEURON | NEURON | 1 | | |
| | PYRAMIDAL CELLS | PYRAMIDAL CELLS | 1 | | |
| | SPERMATOZOA | | | | 1 |
| | STEM ... CELLS | | | | 1 |
| file47 | CELL | CELL | 1 | | |
| | CELLS | CELLS | 1 | | |
| | CELLULAR | GERM CELL | 1 | | 1 |
| | GERM CELL | GERM CELLS | 1 | | |
| | GERM CELLS | GERMLINE STEM CELLS | 1 | | |
| | GERMLINE STEM CELLS | SOMATIC CELLS | | 1 | |
| | MULTIPOTENT ... STEM CELLS | STEM CELL | 1 | | 1 |
| | STEM CELL | STEM CELLS | | 1 | |
| file48 | | | | | |
| file49 | AMACRINE | AMACRINE CELLS | 1 | | 1 |
| | AMACRINE CELLS | CELL | 1 | | |
| | BIPOLAR CELLS | CELLS | 1 | | 1 |
| | CELL | HORIZONTAL CELLS | 1 | | |
| | CELLS | NEURONS | 1 | | |
| | CONE | PHOTORECEPTOR CELL | 1 | | 1 |
| | CONE PHOTORECEPTOR | PHOTORECEPTOR CELLS | 1 | | 1 |
| | CONE PHOTORECEPTORS | PINEALOCYTES | 1 | | 1 |
| | CONES | | | | 1 |
| | GANGLION CELL | | | | 1 |
| | GANGLION CELLS | | | | 1 |
| | HORIZONTAL CELLS | | | | |
| | MÃ¼LLER GLIA | | | | 1 |
| | MULTIPOTENT ... CELLS | | | | 1 |
| | NEURONS | | | | |
| | PHOTORECEPTOR | | | | 1 |
| | PHOTORECEPTOR CELL | | | | |
| | PHOTORECEPTOR CELLS | | | | |
| | PHOTORECEPTOR CELLS IN ... RETINA | | | | 1 |

| | | | | | |
|---|---|---|---|---|---|
| | PHOTORECEPTORS | | | | 1 |
| | PHOTORECEPTORS IN ... RETINA | | | | 1 |
| | PINEALOCYTES | | | | |
| | RETINAL PHOTORECEPTOR | | | | 1 |
| | RETINAL PHOTORECEPTOR CELLS | | | | 1 |
| | RETINAL PHOTORECEPTORS | | | | 1 |
| | ROD ... PHOTORECEPTORS | | | | 1 |
| | ROD PHOTORECEPTOR | | | | 1 |
| | ROD PHOTORECEPTORS | | | | 1 |
| | RODS | | | | 1 |
| file50 | CELL | CELL | 1 | | |
| | CELLS | CELLS | 1 | | |
| | CELLULAR | EMBRYONIC FIBROBLASTS | | 1 | 1 |
| | FIBROBLAST | EMBRYONIC STEM CELL | | 1 | |
| | FIBROBLASTS | EMBRYONIC STEM CELLS | | 1 | |
| | HEPATOCYTES | FIBROBLAST | 1 | | |
| | STEM CELL | FIBROBLASTS | 1 | | |
| | STEM CELLS | HEPATOCYTES | 1 | | |
| | | STEM CELL | 1 | | |
| | | STEM CELLS | 1 | | |
| file51 | APOPTOTIC CELLS | CELL | 1 | | 1 |
| | CELL | CELLS | 1 | | |
| | CELL IN ... EPIBLAST ... REGION | HIPPOCAMPUS | | 1 | 1 |
| | CELLS | LYMPHOCYTES | 1 | | |
| | CELLS IN ... EXTRAEMBRYONIC TISSUES | NEURONS | 1 | | 1 |
| | CELLS OF ... ECTODERM | STEM CELL | 1 | | 1 |
| | CELLULAR | STEM CELLS | 1 | | 1 |
| | DIPLOID ... CELL | TROPHOBLAST CELLS | 1 | | 1 |
| | EXTRAEMBRYONIC CELLS | TROPHOBLAST GIANT CELLS | | 1 | 1 |
| | LYMPHOCYTES | | | | |
| | NEURONAL | | | | 1 |
| | NEURONS | | | | |
| | RED BLOOD CELLS | | | | 1 |
| | STEM CELL | | | | |
| | STEM CELLS | | | | |
| | TROPHOBLAST ... CELL | | | | 1 |
| | TROPHOBLAST ... CELLS | | | | |

| | | | | | |
|---|---|---|---|---|---|
| | TROPHOBLAST CELLS | | | | |
| file52 | PANCREATIC BETA CELL | BETA CELL | | 1 | |
| | | CELL | | 1 | |
| | | PANCREATIC BETA CELL | 1 | | |
| file53 | APOPTOTIC CELL | CELL | 1 | | 1 |
| | APOPTOTIC CELLS | CELLS | 1 | | 1 |
| | BLOOD CELLS | CHONDROCYTE | 1 | | 1 |
| | CELL | ENDOTHELIAL CELLS | 1 | | |
| | CELLS | FIBROBLAST | 1 | | |
| | CELLULAR | FOLLICLE CELLS | 1 | | 1 |
| | CHONDROCYTE | GERM CELL | 1 | | |
| | ENDOTHELIAL ... CELLS | GERM CELLS | 1 | | |
| | ENDOTHELIAL CELL | PRECURSOR CELLS | | 1 | 1 |
| | FIBROBLAST | SERTOLI CELL | 1 | | |
| | FOLLICLE CELLS | SERTOLI CELLS | 1 | | |
| | GERM CELL | SOMATIC CELL | | 1 | |
| | GERM CELLS | VASCULAR ENDOTHELIAL CELLS | 1 | | |
| | PLATELET | | | | 1 |
| | POLYGONAL CELLS | | | | 1 |
| | SERTOLI | | | | 1 |
| | SERTOLI CELL | | | | |
| | SERTOLI CELLS | | | | |
| | SUPPORTING CELL | | | | 1 |
| | SUPPORTING CELLS | | | | 1 |
| | VASCULAR ENDOTHELIAL CELLS | | | | |
| file54 | CELL | CELL | 1 | | |
| | CELLS | CELLS | 1 | | |
| | FIBROBLASTIC CELL | EMBRYONIC FIBROBLASTS | | 1 | 1 |
| | FIBROBLASTS | FIBROBLASTS | 1 | | |
| | STEM ... CELLS | SOMATIC CELL | | 1 | 1 |
| | | SOMATIC CELLS | | 1 | |
| file55 | AMELOBLASTS | AMELOBLASTS | 1 | | |
| | BASAL CELLS | BASAL CELLS | 1 | | |
| | CELL | CELL | 1 | | |
| | CELLS | CELLS | 1 | | |

| | | | | | |
|---|---|---|---|---|---|
| | CELLS OF ... STRATIFIED EPITHELIA | ECTODERMAL CELLS | 1 | | 1 |
| | CELLS OF EPIDERMIS | EMBRYONIC CELLS | | 1 | 1 |
| | CELLULAR | EPIDERMAL CELL | 1 | | 1 |
| | ECTODERMAL CELLS | EPITHELIAL CELL | 1 | | |
| | EPIDERMAL CELL | EPITHELIAL CELLS | 1 | | |
| | EPITHELIAL CELL | KERATINOCYTES | 1 | | |
| | EPITHELIAL CELLS | LYMPHOCYTES | 1 | | |
| | KERATINOCYTES | MACROPHAGES | 1 | | |
| | LYMPHOCYTES | NEUTROPHILS | 1 | | |
| | MACROPHAGES | OOCYTES | 1 | | |
| | NEUTROPHILS | STEM CELL | 1 | | |
| | OOCYTES | STEM CELLS | 1 | | |
| | STEM CELL | THYMIC EPITHELIAL CELLS | | 1 | |
| | STEM CELLS | THYMOCYTE | | 1 | |
| | | THYMOCYTES | | 1 | |
| file56 | BLOOD CELLS | BLOOD CELLS | 1 | | |
| | CELL | CELL | 1 | | |
| | CELLS | CELLS | 1 | | |
| | CELLULAR | EMBRYONIC FIBROBLASTS | | 1 | 1 |
| | ENUCLEATING ... CELLS | EMBRYONIC STEM CELL | | 1 | 1 |
| | FIBROBLASTS | EMBRYONIC STEM CELLS | | 1 | |
| | STEM CELL | FIBROBLASTS | 1 | | |
| | STEM CELLS | STEM CELL | 1 | | |
| | | STEM CELLS | 1 | | |
| file57 | PLATELET | PLATELET | 1 | | |
| file58 | CELL | CELL | 1 | | |
| | CELLS | CELLS | 1 | | |
| | CONE | CEREBRAL CORTEX NEURONS | | 1 | 1 |
| | CONE PHOTORECEPTOR | CORTICAL NEURONS | | 1 | 1 |
| | CONE PHOTORECEPTORS | GLIAL CELLS | 1 | | 1 |
| | DOPAMINERGIC NEURONS | HIPPOCAMPAL NEURONS | | 1 | 1 |
| | GANGLION CELL | HIPPOCAMPUS | | 1 | 1 |
| | GANGLION CELLS | INTERNEURONS | 1 | | 1 |
| | GANGLION CELLS OF ... RETINA | NEURON | 1 | | 1 |
| | GLIA | NEURONS | 1 | | 1 |
| | GLIAL CELLS | OOCYTES | 1 | | |

| | | | | | |
|---|---|---|---|---|---|
| | INTERNEURONS | | | | |
| | NEURON | | | | |
| | NEURONAL | | | | 1 |
| | NEURONS | | | | |
| | OOCYTES | | | | |
| | PHOTORECEPTOR | | | | 1 |
| | PHOTORECEPTOR ... NEURONS OF ... RETINA | | | | 1 |
| | PHOTORECEPTOR NEURONS | | | | 1 |
| | PHOTORECEPTORS | | | | 1 |
| | PHOTOSENSORY NEURON | | | | 1 |
| | PHOTOSENSORY NEURONS | | | | 1 |
| | PHOTOSENSORY NEURONS IN ... RETINA | | | | 1 |
| | PHOTOSENSORY NEURONS OF ... RETINA | | | | 1 |
| | PRIMARY ... NEURON | | | | 1 |
| | PRIMARY ... NEURONS | | | | 1 |
| | RECEPTORAL ... NEURONS | | | | 1 |
| | RECEPTORAL NEURONS | | | | 1 |
| | ROD PHOTORECEPTOR | | | | 1 |
| | ROD PHOTORECEPTOR NEURONAL | | | | 1 |
| | ROD PHOTOSENSORY NEURONS | | | | 1 |
| file59 | APOPTOTIC CELLS | CELL | 1 | | 1 |
| | CELL | CELLS | 1 | | |
| | CELLS | MESENCHYMAL CELL | 1 | | |
| | EPITHELIAL ... CELL | MUSCLE CELL | | 1 | 1 |
| | MESENCHYMAL CELL | MUSCLE CELLS | | 1 | |
| | NC ... CELL | SMOOTH MUSCLE CELL | 1 | | 1 |
| | NCCS | SMOOTH MUSCLE CELLS | 1 | | 1 |
| | NEURAL CREST CELL | STEM CELLS | 1 | | 1 |
| | NEURAL CREST CELLS | | | | 1 |
| | SMOOTH MUSCLE CELL | | | | |
| | SMOOTH MUSCLE CELLS | | | | |
| | STEM CELLS | | | | |
| file60 | CELL | CELL | 1 | | |
| | CELLS | CELLS | 1 | | |
| | CELLULAR | EMBRYONIC CELL | | 1 | 1 |
| | FIBROBLAST | EMBRYONIC CELLS | | 1 | |
| | FIBROBLASTS | EMBRYONIC FIBROBLASTS | | 1 | |

| | | | | | |
|---|---|---|---|---|---|
| | PROMYELOCYTIC | EMBRYONIC STEM CELL | | 1 | 1 |
| | STEM CELL | EMBRYONIC STEM CELLS | | 1 | |
| | STEM CELLS | FIBROBLAST | 1 | | |
| | | FIBROBLASTS | 1 | | |
| | | STEM CELL | 1 | | |
| | | STEM CELLS | 1 | | |
| file61 | APOPTOTIC CELLS | CELL | 1 | | 1 |
| | CELL | CELLS | 1 | | |
| | CELLS | CHONDROCYTE | 1 | | |
| | CHONDROCYTE | CHONDROCYTES | 1 | | |
| | CHONDROCYTES | FIBROBLAST | 1 | | |
| | FIBROBLAST | HYPERTROPHIC CHONDROCYTES | 1 | | |
| | FIBROBLASTIC | OSTEOBLAST | 1 | | 1 |
| | FIBROBLASTIC CELLS | OSTEOBLASTS | 1 | | 1 |
| | HYPERTROPHIC CHONDROCYTES | OSTEOCLAST | 1 | | |
| | OSTEOBLAST | OSTEOCLASTS | 1 | | |
| | OSTEOBLAST CELLS | OSTEOPROGENITOR CELLS | 1 | | 1 |
| | OSTEOBLASTIC | | | | 1 |
| | OSTEOBLASTS | | | | |
| | OSTEOCLAST | | | | |
| | OSTEOCLASTS | | | | |
| | OSTEOPROGENITOR CELLS | | | | |
| | OSTEOPROGENITORS | | | | 1 |
| file62 | BLOOD CELLS | BLOOD CELLS | 1 | | |
| | CELL | CELL | 1 | | |
| | CELLS | CELLS | 1 | | |
| | LYMPHOCYTE | LYMPHOCYTE | 1 | | |
| | LYMPHOCYTES | LYMPHOCYTES | 1 | | |
| | REGULATORY T CELL | REGULATORY T CELL | 1 | | |
| | T CELL | T CELL | 1 | | |
| file63 | CELL | CELL | 1 | | |
| | CELLS | CELLS | 1 | | |
| | EPITHELIAL CELLS | GLIAL CELL | 1 | | 1 |
| | GLIAL CELL | STROMAL CELL | 1 | | |
| | STROMAL CELL | STROMAL CELLS | 1 | | |
| | STROMAL CELLS | | | | |

| | | | | |
|---|---|---|---|---|
| file64 | APOPTOTIC CELLS | CELL | 1 | 1 |
| | CELL | CELLS | 1 | |
| | CELLS | GAMETES | 1 | |
| | CELLULAR | GERM CELL | 1 | 1 |
| | GAMETES | GERM CELLS | 1 | |
| | GERM CELL | GONOCYTES | 1 | |
| | GERM CELLS | MALE GERM CELL | 1 | |
| | GERM-CELL | MALE GERM CELLS | 1 | 1 |
| | GONOCYTES | MULTINUCLEATE CELLS | 1 | 1 |
| | LEYDIG CELLS | OOCYTE | 1 | 1 |
| | MALE GERM CELL | OOCYTES | 1 | |
| | MALE GERM CELLS | SERTOLI CELL | 1 | |
| | MULTINUCLEATE CELLS | SERTOLI CELLS | 1 | |
| | MULTINUCLEATED CELLS | SOMATIC CELL | 1 | 1 |
| | OOCYTE | SOMATIC CELLS | 1 | |
| | OOCYTES | SPERM | 1 | |
| | PRIMARY SPERMATOCYTES | SPERMATID | 1 | |
| | SERTOLI | SPERMATIDS | 1 | 1 |
| | SERTOLI CELL | SPERMATOCYTE | 1 | |
| | SERTOLI CELLS | SPERMATOCYTES | 1 | |
| | SPERM | ZYGOTE | 1 | |
| | SPERMATID | | | |
| | SPERMATIDS | | | |
| | SPERMATOCYTE | | | |
| | SPERMATOCYTES | | | |
| | SPERMATOGONIA | | | 1 |
| | SPERMATOZOA | | | 1 |
| | STEM ... CELLS | | | 1 |
| | ZYGOTE | | | |
| file65 | LYMPHOCYTES | LYMPHOCYTES | 1 | |
| | CELLS | CELLS | 1 | |
| | PURKINJE CELLS | PURKINJE CELLS | 1 | |
| file66 | ADIPOCYTES | ADIPOCYTES | 1 | |
| | AMACRINE | AMACRINE CELL | 1 | 1 |
| | AMACRINE ... CELLS | AMACRINE CELLS | 1 | |
| | AMACRINE CELL | AMACRINE NEURONS | 1 | |

| | | | | |
|---|---|---|---|---|
| AMACRINE CELLS | BIPOLAR NEURONS | 1 | | |
| AMACRINE NEURONS | CELL | 1 | | |
| APOPTOTIC CELLS | CELLS | 1 | | 1 |
| BIPOLAR | CHOLINERGIC NEURONS | 1 | | 1 |
| BIPOLAR CELL | ENTEROCYTES | 1 | | 1 |
| BIPOLAR CELLS | FIBROBLAST | 1 | | 1 |
| BIPOLAR CELLS IN ... RETINA | FIBROBLASTS | 1 | | 1 |
| BIPOLAR NEURONS | FOREBRAIN NEURONS | | 1 | |
| BIPOLARS | INTERNEURONS | 1 | | 1 |
| CELL | MüLLER CELLS | 1 | | |
| CELLS | NEURON | 1 | | |
| CHOLINERGIC NEURONS | NEURONS | 1 | | |
| CNS NEURONS | NULL CELLS | | 1 | |
| CONE | RETINAL CELL | | 1 | 1 |
| CONE BIPOLAR CELLS | RETINAL CELLS | | 1 | 1 |
| CONES | RETINAL PROGENITOR CELL | | 1 | 1 |
| ENTEROCYTES | ROD BIPOLAR CELLS | 1 | | |
| FIBROBLAST | | | | |
| FIBROBLASTS | | | | |
| GANGION CELLS | | | | 1 |
| GANGLION | | | | 1 |
| GANGLION ... CELL | | | | 1 |
| GANGLION ... CELLS | | | | 1 |
| GANGLION ... CELLS IN ... RETINA | | | | 1 |
| GANGLION CELL | | | | 1 |
| GANGLION CELLS | | | | 1 |
| GANGLION NEURONS | | | | 1 |
| GLIAL | | | | 1 |
| HORIZONTAL ... CELLS | | | | 1 |
| HORIZONTAL ... NEURONS | | | | 1 |
| INTERNEURONS | | | | |
| MÃ¼LLER CELLS | | | | 1 |
| MÃ¼LLER GLIA CELL | | | | 1 |
| NEURON | | | | |
| NEURONAL | | | | 1 |
| NEURONS | | | | |
| ON BIPOLAR CELLS | | | | 1 |
| PHOTORECEPTOR | | | | 1 |
| PHOTORECEPTORS | | | | 1 |

| | | | | | |
|---|---|---|---|---|---|
| | RETINAL ... GANGLION ... CELLS | | | | 1 |
| | RETINAL ... ROD CELLS | | | | 1 |
| | RETINAL BIPOLAR ... CELLS | | | | 1 |
| | ROD ... BIPOLAR CELLS | | | | |
| | ROD ... CELL | | | | 1 |
| | ROD ... CELLS | | | | 1 |
| | ROD ... CELLS IN ... RETINA | | | | 1 |
| | ROD BIPOLAR ... CELLS | | | | |
| | ROD BIPOLAR CELLS | | | | |
| | ROD PHOTORECEPTORS | | | | 1 |
| | RODS | | | | 1 |
| file67 | CELL | ARCS | | 1 | |
| | CELLS | CELL | 1 | | |
| | EGGS | CELLS | 1 | | 1 |
| | GAMETE | EMBRYONIC STEM CELL | | 1 | 1 |
| | GAMETES | GAMETES | 1 | | |
| | GERM CELL | GERM CELL | 1 | | |
| | OOCYTE | OOCYTE | 1 | | |
| | OOCYTES | OOCYTES | 1 | | |
| | SPERM | SPERM | 1 | | |
| | SPERMATIDS | SPERMATIDS | 1 | | |
| | SPERMATOCYTE | SPERMATOCYTE | 1 | | |
| | SPERMATOCYTES | SPERMATOCYTES | 1 | | |
| | SPERMATOGONIA | STEM CELL | 1 | | 1 |
| | SPORE | | | | 1 |
| | STEM CELL | | | | |
| | STEM CELLS | | | | 1 |
| Total | | | 513 | 144 | 343 |
| | | Accuracy | 0,78 | | |
| | | Recall | 0,60 | | |
| | | F-Measure | 0,68 | | |
