## Supplementary_Excel_file_S1 CRAFT 1.0 derived for "OntoContext, a new python package for gene contextualization based on the annotation of biomedical texts"

| Num file | Reference CRAFT 1.0 Annotation | OntoContext Annotation | TP | FP | FN |
|---|---|---|---|---|---|
| file1 | CONES | PHOTORECEPTORS | | 1 | |
| | MACROPHAGES | MACROPHAGES | 1 | | |
| | CELLS | CELLS | 1 | | |
| | RETINAL GANGLION CELL | RETINAL GANGLION CELL | 1 | | |
| | ROD ... PHOTORECEPTORS | GANGLION CELL | | 1 | 1 |
| | RODS | CELL | | 1 | |
| | CONE PHOTORECEPTORS | CONE | | 1 | |
| | | RODS | 1 | | |
| | | CONE PHOTORECEPTORS | 1 | | |
| | | CONES | 1 | | |
| | | GANGLION | | 1 | |
| file2 | CELL | CELL | 1 | | |
| | STEM ... CELLS | CELLS | 1 | | 1 |
| | CELLS | | | | |
| file3 | OLIGODENDROCYTE | | | | |
| | CELLULAR | 1 | | | 1 |
| file4 | CELLS | CELLS | 1 | | |
| | CELL | CELL | 1 | | |
| | BLASTOMERES | BLASTOMERES | 1 | | |
| | ENDODERM CELLS | ENDODERM CELLS | 1 | | |
| | STEM CELLS | STEM CELLS | 1 | | |
| | EUKARYOTIC CELLS | EUKARYOTIC CELLS | 1 | | |
| file5 | CELLULAR | CELLULAR | 1 | | |
| | EPITHELIAL CELLS | EPITHELIAL CELLS | 1 | | |
| | CELLS | CELLS | 1 | | |
| | CELL | CELL | 1 | | |
| | B LYMPHOCYTES | OSTEOBLASTS | 1 | | 1 |
| | EPITHELIAL CELL | EPITHELIAL CELL | 1 | | |
| | OSTEOBLASTS | LYMPHOCYTES | | 1 | |
| | MUSCLE CELL | MUSCLE CELL | 1 | | |
| | MACROPHAGE | MACROPHAGE | 1 | | |
| file6 | SUPPORTING CELL | SUPPORTING CELL | 1 | | |

| | |
|---|---|
| Accuracy | 0,94 |
| Recall | 0,89 |
| F-Measure | 0,91 |

| | | | | | |
|---|---|---|---|---|---|
| | NEURON | NEURON | 1 | | |
| | SUPPORTING CELLS | SUPPORTING CELLS | 1 | | |
| | NEURON AFFERENTS | PILLAR CELLS | 1 | | |
| | OUTER HAIR CELL | OUTER HAIR CELL | 1 | | |
| | SENSORY NEURONS | SENSORY NEURONS | 1 | | |
| | PRIMARY NEURONS | PRIMARY NEURONS | 1 | | |
| | INNER ... HAIR CELLS | CELL | 1 | | 1 |
| | VESTIBULAR HAIR CELL | NEURONS | 1 | | |
| | CELL | NEURONAL | 1 | | |
| | NEURONS | GANGLION | 1 | | |
| | NEURONAL | HAIR CELL | 1 | | |
| | INNER HAIR CELL | OUTER HAIR CELLS | 1 | | |
| | HAIR CELL | CELLS | 1 | | |
| | OUTER HAIR CELLS | SENSORY NEURON | 1 | | |
| | CELLS | CELLULAR | 1 | | |
| | SENSORY NEURON | NEURON AFFERENTS | 1 | | |
| | CELLULAR | GANGLION CELLS | | 1 | |
| | PILLAR CELLS | HAIR CELLS | 1 | | |
| | HAIR CELLS | MOTONEURONS | 1 | | |
| | MOTONEURONS | INNER HAIR CELLS | 1 | | |
| | INNER HAIR CELLS | DEITERS' CELLS | 1 | | |
| | DEITERS' CELLS | | | | |
| file7 | EOSINOPHILES | EOSINOPHILES | 1 | | |
| | PLATELETS | CARDIOMYOCYTES | 1 | | |
| | CELLULAR | MONOCYTES | | 1 | |
| | LEUCOCYTES | LEUCOCYTES | 1 | | |
| | RED BLOOD CELLS | RED BLOOD CELLS | 1 | | |
| | CELLS | CELLS | 1 | | |
| | CARDIOMYOCYTES | PLATELETS | 1 | | |
| | BASOPHILES | BASOPHILES | 1 | | |
| | ERYTHROCYTES | ERYTHROCYTES | 1 | | |
| | ERYTHROCYTE | BLOOD CELL | | 1 | |
| | CELL | ERYTHROCYTE | 1 | | |
| | PLATELET | CELL | 1 | | |
| | MONOCYTES | PLATELET | 1 | | |
| | RED BLOOD CELL | BLOOD CELLS | | 1 | |
| | LYMPHOCYTES | CELLULAR | 1 | | |

| | | | | | |
|---|---|---|---|---|---|
| | NEUTROPHILES | RED BLOOD CELL | 1 | | |
| | LEUKOCYTES | LYMPHOCYTES | 1 | | |
| | | NEUTROPHILES | 1 | | |
| | | LEUKOCYTES | 1 | | |
| file8 | CELL | CELL | 1 | | |
| | NEURONAL | NEURONAL | 1 | | |
| file9 | SPERM | SPERM | 1 | | |
| | SPERMATID | SPERMATID | 1 | | |
| | CELLS | CELLS | 1 | | |
| | NEURON | NEURON | 1 | | |
| | T-CELL | T-CELL | 1 | | |
| | CELL | ERYTHROID | | 1 | |
| | NEURONS | CELL | 1 | | |
| | ERYTHROID CELLS | NEURONS | 1 | | |
| | NEURONAL | ERYTHROID CELLS | 1 | | |
| | OLFACTORY NEURON | NEURONAL | 1 | | |
| | OLFACTORY NEURONS | OLFACTORY NEURON | 1 | | |
| | | OLFACTORY NEURONS | 1 | | |
| file10 | NEURON | NEURONAL CELLS | 1 | | |
| | CELLS | CELLULAR | 1 | | |
| | NEURONAL CELLS | CELLS | 1 | | |
| | ASTROCYTES | NEURON | 1 | | |
| | CELL | ASTROCYTES | 1 | | |
| | NEURONS | CELL | 1 | | |
| | CELLULAR | NEURONS | 1 | | |
| | NEURONAL CELL | NEURONAL | 1 | | |
| file11 | PIGMENT CELL | PIGMENT CELL | 1 | | |
| | MELANOCYTE | MELANOCYTE | 1 | | |
| | STEM CELLS | DERMAL PAPILLA CELLS | 1 | | 1 |
| | DERMAL PAPILLA CELLS | CELLS | 1 | | |
| | CELLS | MELANOBLAST | 1 | | |
| | MELANOBLAST | DERMAL PAPILLAE CELLS | 1 | | |
| | DERMAL PAPILLAE CELLS | PIGMENT-CELL | 1 | | |
| | PIGMENT-CELL | CELL | 1 | | |

| | | | | | |
|---|---|---|---|---|---|
| | CELL | KERATINOCYTES | 1 | | |
| | KERATINOCYTES | CELLULAR | 1 | | |
| | CELLULAR | MELANOCYTES | 1 | | |
| | MELANOCYTES | PIGMENT CELLS | 1 | | |
| | MESENCHYMAL CELLS | MESENCHYMAL CELLS | 1 | | |
| | PIGMENT CELLS | MELANOBLASTS | 1 | | |
| | MELANOBLASTS | | | | |
| file12 | FIBROBLASTS | FIBROBLASTS | 1 | | |
| | ENDOTHELIAL CELL | ENDOTHELIAL CELL | 1 | | |
| | CELLULAR | EPITHELIAL CELLS | 1 | | |
| | EPITHELIAL CELLS | PLATELET | 1 | | |
| | CELLS IN ... MESENCHYMAL REGIONS | CELLS | 1 | | 1 |
| | CELLS | LUNG EPITHELIAL CELL | 1 | | |
| | CELLS IN ... EPITHELIAL ... REGIONS | CELL | 1 | | 1 |
| | EPITHELIAL CELLS ... IN ... LUNGS | MESENCHYME CELLS | 1 | | 1 |
| | CELL | CELLULAR | 1 | | |
| | PLATELET | LUNG EPITHELIAL CELLS | 1 | | |
| | EPITHELIAL ... CELLS | | | | 1 |
| | MESENCHYME CELLS | | | | |
| | EPITHELIAL CELLS IN ... LUNGS | | | | 1 |
| | LUNG EPITHELIAL CELL | | | | |
| | LUNG EPITHELIAL CELLS | | | | |
| file13 | CELL | FIBROBLASTS | 1 | | |
| | EPITHELIAL CELLS | EPITHELIAL CELLS | 1 | | |
| | CELLS | CELLS | 1 | | |
| | FIBROBLASTS | CELL | 1 | | |
| | CELLULAR | CELLULAR | 1 | | |
| | ENTEROCYTES | ENTEROCYTES | 1 | | |
| file14 | NEURONAL | NEURONAL | 1 | | |
| | DOPAMINE CELLS | CELLULAR | 1 | | |
| | CELLS | CELLS | 1 | | |
| | CELLULAR | CELL | | 1 | |
| | DOPAMINERGIC CELL | DOPAMINE CELLS | 1 | | |
| | DOPAMINERGIC CELLS | DOPAMINERGIC CELL | 1 | | |
| | | DOPAMINERGIC CELLS | 1 | | |

| | | | | |
|---|---|---|---|---|
| file15 | CELL | CELL | 1 | |
| | OOCYTE | OOCYTE | 1 | |
| | CELLS | CELLS | 1 | |
| | CELLULAR | CELLULAR | 1 | |
| file16 | PROERYTHROBLAST | PROERYTHROBLAST | 1 | |
| | RBCS | HEMATOPOIETIC PROGENITORS | 1 | |
| | STEM CELL | RBCS | 1 | |
| | ERYTHROCYTES | STEM CELL | 1 | |
| | STEM CELLS | ERYTHROCYTES | 1 | |
| | HEMATOPOIETIC CELLS | PROMYELOCYTIC | | 1 |
| | RETICULOCYTE | STEM CELLS | 1 | |
| | WHITE BLOOD CELLS | HEMATOPOIETIC CELLS | | 1 |
| | STEM ... CELLS | RETICULOCYTE | 1 | |
| | HEMATOPOIETIC STEM CELL | HEMATOPOIETIC STEM CELL | 1 | |
| | RED BLOOD CELL | RED BLOOD CELL | 1 | |
| | T-CELLS | T-CELLS | 1 | |
| | CELL | CELL | 1 | |
| | RED ... BLOOD CELLS | FIBROBLAST CELL | 1 | |
| | HEMATOPOIETIC PROGENITOR CELLS | HEMATOPOIETIC PROGENITOR CELLS | 1 | |
| | HEMATOPOIETIC PROGENITORS | ERYTHROID CELL | 1 | |
| | ERYTHROID CELLS | BLOOD CELLS | 1 | |
| | MESENCHYMAL CELLS | MESENCHYMAL CELLS | 1 | |
| | CELLS | EGGS | 1 | |
| | ERYTHROID CELL | WHITE BLOOD CELLS | 1 | |
| | APOPTOTIC CELLS | APOPTOTIC CELLS | 1 | |
| | MYELOID CELLS | MYELOID CELLS | 1 | |
| | EGGS | CELLS | 1 | |
| | ERYTHROID | ERYTHROID | 1 | |
| | FIBROBLASTS | FIBROBLASTS | 1 | |
| | CIRCULATING CELLS | CIRCULATING CELLS | 1 | |
| | FIBROBLAST | FIBROBLAST | 1 | |
| | MYELOID | MYELOID | 1 | |
| | LYMPHOCYTES | LYMPHOCYTES | 1 | |
| | FIBROBLAST CELL | HEMATOPOIETIC STEM CELLS | 1 | |
| | HEMATOPOIETIC STEM CELLS | MESODERMAL CELLS | 1 | |
| | MESODERMAL CELLS | ERYTHROID PROGENITOR CELLS | 1 | |

| | | | | | |
|---|---|---|---|---|---|
| | ERYTHROID PROGENITOR CELLS | MACROPHAGES | 1 | | |
| | MACROPHAGES | RED BLOOD CELLS | 1 | | |
| | RED BLOOD CELLS | ERYTHROID PROGENITORS | 1 | | |
| | ERYTHROID PROGENITORS | BLOOD CELL | 1 | | |
| | BLOOD CELL | ERYTHROID CELLS | 1 | | |
| | BLOOD CELLS | PROERYTHROBLASTS | 1 | | |
| | PROERYTHROBLASTS | | | | |
| file17 | CELL | CELL | 1 | | |
| | CELLS | CELLS | 1 | | |
| | CELLS IN VIVO | CELLS IN VIVO | 1 | | |
| | STEM CELLS | STEM CELLS | 1 | | |
| | APOPTOTIC CELLS | APOPTOTIC CELLS | 1 | | |
| file18 | FIBROCYTES | FIBROCYTES | 1 | | |
| | VESTIBULAR DARK CELLS | HAIR CELLS | 1 | | |
| | BASAL CELLS | BASAL CELLS | 1 | | |
| | HAIR CELLS | HAIR CELL | 1 | | |
| | EPITHELIAL CELLS | EPITHELIAL CELLS | 1 | | |
| | CELLS | CELLS | 1 | | |
| | STEM CELL | STEM CELL | 1 | | |
| | CELL | CELL | 1 | | |
| | HAIR CELL | BASAL CELL | 1 | | |
| | VESTIBULAR HAIR CELLS | VESTIBULAR DARK CELLS | 1 | | |
| | TRANSITIONAL CELLS | TRANSITIONAL CELLS | 1 | | |
| | | GANGLION | | 1 | |
| file19 | MOTOR NEURON | MOTOR NEURON | 1 | | |
| | ZYGOTES | NEURON | | 1 | |
| | ADIPOCYTE | ZYGOTES | 1 | | |
| | CELL | ADIPOCYTE | 1 | | |
| | MOTONEURON | CELL | 1 | | |
| | CELLULAR | MOTONEURON | 1 | | |
| | | CELLULAR | 1 | | |
| file20 | MESENCHYMAL ... CELLS | EPITHELIAL CELLS | 1 | | 1 |
| | EPITHELIAL CELLS | ERYTHROCYTES | 1 | | |
| | ERYTHROCYTES | PERITONEAL MACROPHAGES | 1 | | |

| | | | | | |
|---|---|---|---|---|---|
| | PERITONEAL MACROPHAGES | APOPTOTIC CELL | 1 | | |
| | STEM ... CELLS | CELL | 1 | | 1 |
| | APOPTOTIC CELL | NEURONAL | 1 | | |
| | CELL | MEGAKARYOCYTES | 1 | | |
| | PLATELET | GANGLION | | 1 | |
| | NEURONAL | PHOTORECEPTOR | | 1 | |
| | CELLS | APOPTOTIC CELLS | 1 | | |
| | APOPTOTIC ... CELLS | CELLS | 1 | | |
| | APOPTOTIC CELLS | NEURONAL CELLS | 1 | | |
| | MEGAKARYOCYTES | MONOCYTE | 1 | | |
| | NEURONAL CELLS | FIBROBLASTS | 1 | | |
| | MONOCYTE | PHAGOCYTES | 1 | | |
| | FIBROBLASTS | CELLULAR | 1 | | |
| | PHAGOCYTES | NEUTROPHILS | 1 | | |
| | CELLULAR | PIGMENTED EPITHELIAL CELLS | 1 | | |
| | NEUTROPHILS | GANGLION CELLS | | 1 | |
| | PIGMENTED EPITHELIAL CELLS | MACROPHAGE | 1 | | |
| | HEMATOPOIETIC STEM CELLS | EARLY ERYTHROBLAST | 1 | | |
| | MACROPHAGE | PHOTORECEPTOR CELLS | 1 | | |
| | EARLY ERYTHROBLAST | MACROPHAGES | 1 | | |
| | PHOTORECEPTOR CELLS | PIGMENTED CELLS | 1 | | |
| | MACROPHAGES | | | | |
| | PIGMENTED CELLS | | | | |
| | CELLS IN ... MESENCHYME | | | | 1 |
| file21 | MONONUCLEAR CELLS | MONONUCLEAR CELLS | 1 | | |
| | CELLS | MACROPHAGES | 1 | | |
| | HYPERTROPHIC CHONDROCYTES | CELLS | 1 | | 1 |
| | MACROPHAGES | SYNOVIOCYTES | 1 | | |
| | NEUTROPHILS | CELL | 1 | | |
| | EGGS | OSTEOBLASTS | 1 | | |
| | SYNOVIOCYTES | CHONDROCYTES | 1 | | |
| | CELL | CELLULAR | 1 | | |
| | FIBROBLAST | SYNOVIAL CELLS | 1 | | 1 |
| | SYNOVIAL CELLS | NEUTROPHILS | 1 | | |
| | CHONDROCYTES | EGGS | 1 | | |
| | CELLULAR | | | | |
| | OSTEOBLASTS | | | | |

| | | | | | |
|---|---|---|---|---|---|
| | MESENCHYMAL CELLS | | | | 1 |
| | CARTILAGE CELLS | | | | 1 |
| file22 | MOTOR NEURON | MOTOR NEURON | 1 | | |
| | GLIA | GANGLION NEURONS | | 1 | |
| | CELLS | CELLS | 1 | | |
| | NEURON | NEURON | 1 | | |
| | CELL | CELL | 1 | | |
| | NEURONS | NEURONS | 1 | | |
| | MOTOR NEURONS | MOTOR NEURONS | 1 | | |
| | SENSORY NEURONS | SENSORY NEURONS | 1 | | |
| | | GLIA | 1 | | |
| | | GANGLION | | 1 | |
| file23 | NEURONAL CELLS | GRANULE NEURON | 1 | | 1 |
| | GRANULE NEURON | NEURONAL | 1 | | |
| | PURKINJE NEURON | PURKINJE NEURON | 1 | | |
| | CELLS | CELLS | 1 | | |
| | NEURON | NEURON | 1 | | |
| | CNS NEURONS | CNS NEURONS | 1 | | |
| | CELLULAR | CELL | 1 | | |
| | CELL | NEURONS | 1 | | |
| | NEURONS | CELLULAR | 1 | | |
| | NEURONAL | GRANULE NEURONS | 1 | | |
| | GRANULE NEURONS | NEURONAL CELL | 1 | | |
| | NEURONAL CELL | | | | |
| file24 | HAIR CELL | HAIR CELL | 1 | | |
| | KERATINOCYTE | KERATINOCYTE | 1 | | |
| | SQUAMOUS CELL | CELLULAR | 1 | | |
| | CELLULAR | EPIDERMAL CELLS | 1 | | |
| | EPITHELIAL CELLS | EPITHELIAL CELLS | 1 | | |
| | CELLS | CELLS | 1 | | |
| | GRANULAR LAYER CELLS | T CELL | 1 | | |
| | T CELL | CELL | 1 | | |
| | CELL | FIBROBLAST | 1 | | |
| | EPITHELIAL ... CELL | SQUAMOUS CELL | 1 | | |
| | FIBROBLAST | KERATINOCYTES | 1 | | |

| | | | | |
|---|---|---|---|---|
| | BASAL ... CELLS | HEPATOCYTES | 1 | 1 |
| | KERATINOCYTES | MESENCHYMAL CELLS | 1 | |
| | MOTILE ... CELLS | | | 1 |
| | HEPATOCYTES | | | |
| | CELLS OF ... ECTODERM | | | 1 |
| | MESENCHYMAL CELLS | | | |
| | ECTODERMAL CELLS | | | 1 |
| | CELLS WITHIN ... EPIDERMIS | | | 1 |
| | EPIDERMAL CELLS | | | |
| file25 | PHOTORECEPTORS | CELL | 1 | |
| | PHOTORECEPTORS ... R6 | BIPOLAR CELLS | 1 | 1 |
| | HORIZONTAL CELLS | PHOTORECEPTOR CELLS | 1 | |
| | PHOTORECEPTORS R1 | PHOTORECEPTOR | 1 | 1 |
| | CONE | BIPOLAR CELL | 1 | |
| | OFF ... BIPOLAR CELLS | CELLS | 1 | 1 |
| | SENSORY NEURONS | CONE | 1 | |
| | RODS | HORIZONTAL CELLS | 1 | |
| | MÃ¼LLER CELLS | ROD PHOTORECEPTORS | 1 | 1 |
| | HORIZONTAL ... CELL | BIPOLAR | 1 | |
| | CELL | NEURONS | 1 | |
| | BIPOLAR CELL | ROD BIPOLAR CELLS | 1 | |
| | ROD BIPOLAR CELLS | ROD BIPOLAR | 1 | |
| | NEURONAL | NEURONAL | 1 | |
| | NEURONS | RODS | 1 | |
| | PHOTORECEPTOR | SENSORY NEURONS | 1 | |
| | CELLS | PHOTORECEPTORS | 1 | |
| | HORIZONTAL CELL | | | |
| | BIPOLAR CELLS | | | |
| | ROD BIPOLAR | | | |
| | PHOTORECEPTOR CELLS | | | |
| | HORIZONTAL ... CELLS | | | |
| | ROD PHOTORECEPTORS | | | |
| | R1 ... PHOTORECEPTORS | | | 1 |
| | ON ... BIPOLAR CELLS | | | 1 |
| | R6 PHOTORECEPTORS | | | 1 |
| file26 | CELL | HEPATOCYTE | 1 | |

| | | | | | |
|---|---|---|---|---|---|
| | NEURONAL | STEM CELLS | 1 | | |
| | OOCYTES | NEURONAL | 1 | | |
| | CELLULAR | CELLULAR | 1 | | |
| | CELLS | OOCYTES | 1 | | |
| | NEURON | CELLS | 1 | | |
| | LYMPHOCYTIC | NEURON | 1 | | |
| | ASTROCYTES | LYMPHOCYTIC | 1 | | |
| | PYRAMIDAL NEURONS | ASTROCYTES | 1 | | |
| | HEPATOCYTE | PYRAMIDAL NEURONS | 1 | | |
| | PURKINJE ... CELL | MYOCYTES | 1 | | 1 |
| | GRANULE CELL | CELL | 1 | | |
| | NEURONS | GRANULE CELL | 1 | | |
| | PHAGOCYTIC CELLS | NEURONS | 1 | | |
| | STEM CELLS | HEPATOCYTES | 1 | | |
| | ASTROCYTIC | MICROGLIAL | | 1 | |
| | HEPATOCYTES | ASTROCYTIC | 1 | | |
| | GLIAL | GLIAL | 1 | | |
| | CARDIAC MYOCYTES | CARDIAC MYOCYTES | 1 | | |
| file27 | CELLS OF ... STRATUM GRANULOSUM | ASTROGLIAL CELL | 1 | | 1 |
| | ASTROGLIAL CELL | GLIAL CELL | 1 | | |
| | GLIAL CELL | NEURON | 1 | | |
| | NEURON | GLIAL CELLS | 1 | | |
| | GLIAL CELLS | PYRAMIDAL NEURON | 1 | | 1 |
| | ASTROGLIAL ... CELL | PURKINJE-CELLS | 1 | | |
| | PYRAMIDAL NEURON | CELL | 1 | | |
| | ENDODERMAL ... CELLS | PURKINJE-CELL | 1 | | 1 |
| | PURKINJE-CELLS | NEURONS | 1 | | |
| | NEURONAL ... CELLS | MYOBLAST | 1 | | |
| | CELL | NEURONAL | 1 | | |
| | PURKINJE-CELL | ASTROCYTIC | 1 | | |
| | PLATELET | PURKINJE CELLS | 1 | | |
| | MYOBLAST | PLATELET | 1 | | |
| | NEURONAL | CELLS | 1 | | |
| | ASTROCYTIC | NEURONAL CELLS | 1 | | |
| | PURKINJE CELLS | ASTROCYTES | 1 | | |
| | CELLS | PYRAMIDAL NEURONS | 1 | | |
| | NEURONAL CELLS | CHROMAFFIN CELLS | 1 | | |

| | | | | | |
|---|---|---|---|---|---|
| | ASTROCYTES | RADIAL GLIAL CELLS | 1 | | |
| | PYRAMIDAL NEURONS | CELLULAR | 1 | | |
| | FIBROBLASTS | GLIAL | 1 | | |
| | RADIAL GLIAL CELLS | MESODERMAL CELLS | 1 | | |
| | NEURONS | FIBROBLASTS | 1 | | |
| | CELLULAR | RED BLOOD CELLS | 1 | | |
| | GLIAL | ASTROCYTIC CELL | 1 | | |
| | ASTROGLIAL CELLS | BLOOD CELLS | 1 | | |
| | MESODERMAL CELLS | ENDODERM CELLS | 1 | | |
| | CHROMAFFIN CELLS | | | | |
| | RED BLOOD CELLS | | | | |
| | ASTROCYTIC CELL | | | | |
| | ENDODERM CELLS | | | | |
| | NEURONAL ... CELL | | | | 1 |
| | ASTROCYTE | | | | |
| file28 | MOTOR NEURON | MOTOR NEURON | 1 | | |
| | NEUROBLASTS | NEUROBLASTS | 1 | | |
| | NEURONAL | CELLULAR | 1 | | |
| | APOPTOTIC CELLS | APOPTOTIC CELLS | 1 | | |
| | PEPTIDERGIC NEURONS | PEPTIDERGIC NEURONS | 1 | | |
| | CELLS | CELLS | 1 | | |
| | NEURON | NEURON | 1 | | |
| | AFFERENT SENSORY NEURONS | SENSORY NEURON | 1 | | |
| | SENSORY NEURON | CELL | 1 | | |
| | CELL | NEURONS | 1 | | |
| | NEURONS | AFFERENT CELL | 1 | | |
| | AFFERENT CELL | NEURONAL | 1 | | |
| | CELLULAR | MOTOR NEURONS | 1 | | |
| | AFFERENTS ... NEURONS | NEUROBLAST | 1 | | 1 |
| | MOTOR NEURONS | SENSORY NEURONS | 1 | | |
| | NEUROBLAST | NEURONAL CELL | 1 | | |
| | SENSORY NEURONS | GANGLION | | 1 | |
| | SENSORY ... NEURONS | | | | |
| | NEURONAL CELL | | | | |
| file29 | STEM ... CELLS | GRANULE CELLS | 1 | | 1 |
| | GRANULE CELLS | PURKINJE CELL | 1 | | |

| | | | | |
|---|---|---|---|---|
| | PURKINJE CELL | CELLS | 1 | |
| | CELLS | SCHWANN CELLS | 1 | |
| | NEURON | CELL | 1 | |
| | SCHWANN CELLS | GRANULE CELL | 1 | |
| | SCHWANN-CELL | NEURONS | 1 | 1 |
| | CELL | EGGS | 1 | |
| | GRANULE CELL | NEURONAL | 1 | |
| | NEURONS | SPERMATOZOA | 1 | |
| | EGGS | GANGLION | 1 | |
| | NEURONAL | SCHWANN CELL | 1 | |
| | SPERMATOZOA | NEURONAL CELL | 1 | |
| | SCHWANN CELL | | | |
| | NEURONAL CELL | | | |
| file30 | CELL | CELL | 1 | |
| | CELLS | CELLS | 1 | |
| file31 | ENDOTHELIAL CELL | ENDOTHELIAL CELL | 1 | |
| | GOBLET CELL | IMMUNE CELLS | 1 | |
| | IMMUNE CELL | GOBLET CELL | 1 | |
| | MAST CELL | IMMUNE CELL | 1 | |
| | PANETH CELL | EPITHELIAL CELLS | 1 | |
| | CELLS | PANETH CELL | 1 | |
| | STEM CELL | CELLS | 1 | |
| | BLOOD CELL | STEM CELL | 1 | |
| | HEMATOPOIETIC CELL | MAST CELL | 1 | |
| | CELL | BLOOD CELL | 1 | |
| | LYMPHOCYTES | CELL | 1 | |
| | EPITHELIAL CELLS | LYMPHOCYTES | 1 | |
| | MAST CELLS | MAST CELLS | 1 | |
| | T CELL | T CELL | 1 | |
| | NEUTROPHIL | GOBLET CELLS | 1 | |
| | GOBLET CELLS | NEUTROPHILS | 1 | |
| | NEUTROPHILS | MONONUCLEAR CELL | 1 | |
| | MONONUCLEAR CELL | NEUTROPHIL | 1 | 1 |
| | IMMUNE CELLS | LYMPHOCYTE | 1 | |
| | LYMPHOCYTE | | | |

| | | | | | |
|---|---|---|---|---|---|
| file32 | CELLS | CELLS | 1 | | |
| | TRCS | TRCS | 1 | | |
| | TASTE CELLS | TASTE CELLS | 1 | | |
| | NATURAL KILLER CELL | NATURAL KILLER CELL | 1 | | |
| | TASTE RECEPTOR CELLS | CELL | | 1 | |
| | CELLULAR | TASTE RECEPTOR CELLS | 1 | | |
| | CELLS ... OF ... EPITHELIUM | CELLULAR | 1 | | 1 |
| file33 | GLIA | GLIA | 1 | | |
| | CELLS | CELLS | 1 | | |
| | CELL | CELL | 1 | | |
| | NEURONS | NEURONS | 1 | | |
| | NEURONAL | NEURONAL | 1 | | |
| | NEURONES | NEURONES | 1 | | |
| file34 | MUSCLE PRECURSOR CELLS | CELL | 1 | | |
| | CELLS | MUSCLE PRECURSOR CELLS | 1 | | |
| | MYOGENIC PRECURSORS | CELLS | | 1 | 1 |
| | HEPATOCYTE | HEPATOCYTE | 1 | | |
| | MUSCLE PRECURSOR CELL | MUSCLE PRECURSOR CELL | 1 | | |
| | MUSCLE PRECURSORS | MUSCLE PRECURSORS | 1 | | |
| file35 | PRIMORDIAL GERM CELLS | STEM CELLS | 1 | | |
| | PGCS | CELLULAR | 1 | | |
| | STEM CELLS | GERM CELLS | 1 | | |
| | CELLULAR | CELLS | 1 | | |
| | EXTRAEMBRYONIC CELLS | ENDODERM CELLS | 1 | | |
| | CELLS | PCGS | 1 | | |
| | PRIMITIVE GERM CELLS ... PCGS | CELL | 1 | | |
| | PCGS | NEURONS | 1 | | |
| | CELL | PGCS | 1 | | |
| | NEURONS | CONE | | 1 | |
| | ENDODERM CELLS | NEURONAL | 1 | | |
| | NEURONAL | EXTRAEMBRYONIC CELLS | 1 | | |
| | ENDODERM ... CELLS | PRIMORDIAL GERM CELLS | 1 | | |
| | | NEURONAL CELL | | 1 | |
| file36 | PHOTORECEPTORS | PHOTORECEPTORS | 1 | | |

| | | | | | |
|---|---|---|---|---|---|
| | GLIAL CELL | GLIAL CELL | 1 | | |
| | CONE | CONE | 1 | | |
| | CONE CELLS | CONE CELLS | 1 | | |
| | RODS | RODS | 1 | | |
| | PHOTORECEPTOR CELL | PHOTORECEPTOR CELL | 1 | | |
| | MÃ¼LLER GLIAL | CONE CELL | 1 | | 1 |
| | MÃ¼LLER GLIA | CONES | 1 | | 1 |
| | CONE CELL | RADIAL GLIA | 1 | | |
| | BIPOLAR CELLS | CELL | 1 | | |
| | CONES | GANGLION | | 1 | |
| | RADIAL GLIA | ROD CELL | 1 | | |
| | CELL | GLIA | | 1 | |
| | PHOTORECEPTORS IN ... RETINA | PHOTORECEPTOR | 1 | | 1 |
| | CONE PHOTORECEPTORS | CELLS | 1 | | |
| | ROD CELL | GANGLION CELL | 1 | | |
| | PHOTORECEPTOR | ASTROCYTES | 1 | | |
| | CELLS | BIPOLAR | | 1 | |
| | ASTROCYTES | GLIAL | | 1 | |
| | PHOTORECEPTOR CELLS | PHOTORECEPTOR CELLS | 1 | | |
| | ROD PHOTORECEPTOR | ROD PHOTORECEPTOR | 1 | | |
| | ROD ... PHOTORECEPTORS | CONE PHOTORECEPTORS | 1 | | |
| | ROD PHOTORECEPTOR CELLS | | | | 1 |
| file37 | CELL | CELL | 1 | | |
| | OOCYTES | OOCYTES | 1 | | |
| | CELLS | CELLS | 1 | | |
| file38 | STEM ... CELLS | CELL | 1 | | 1 |
| | SPERM | SPERM | 1 | | |
| | CELLS | CELLS | 1 | | |
| | MYOCYTES | MYOCYTE | 1 | | |
| | CELL | MYOCYTES | 1 | | |
| | STEM CELLS | | | | 1 |
| | MYOCYTE | | | | |
| file39 | OOCYTES | OOCYTES | 1 | | |
| | GRANULE CELLS | GRANULE CELLS | 1 | | |
| | OUTPUT NEURONS | CELLULAR | 1 | | |

| | | | | | |
|---|---|---|---|---|---|
| | PRINCIPAL NEURONS | PRINCIPAL NEURONS | 1 | | |
| | CELLULAR | CELLS | 1 | | |
| | CELLS | PYRAMIDAL NEURONS | 1 | | |
| | PRINCIPAL ... NEURONS | CELL | 1 | | 1 |
| | PYRAMIDAL NEURONS | NEURONS | 1 | | |
| | CELL | PYRAMIDAL CELLS | 1 | | |
| | NEURONS | NEURONAL | 1 | | |
| | PYRAMIDAL CELLS | OLFACTORY SENSORY NEURONS | 1 | | |
| | NEURONAL | SENSORY NEURONS | | 1 | |
| | OLFACTORY SENSORY NEURONS | OUTPUT NEURONS | 1 | | |
| file40 | FIBROBLASTS | FIBROBLASTS | 1 | | |
| | HEPATOCYTE | CELL | 1 | | |
| | CELLS | CELLS | 1 | | |
| | CELL | HEPATOCYTE | 1 | | |
| | GLIAL CELLS | GLIAL CELLS | 1 | | |
| | FIBROBLAST CELL | FIBROBLAST | | 1 | |
| | CELLULAR | FIBROBLAST CELL | 1 | | |
| | | CELLULAR | 1 | | |
| | | GLIAL | | 1 | |
| file41 | B CELL | FIBROBLASTS | 1 | | 1 |
| | CELL | EPIDERMAL CELLS | 1 | | |
| | EPIDERMAL CELLS | EPITHELIAL CELLS | 1 | | |
| | EPITHELIAL CELLS | CELLS | 1 | | |
| | CELLS | CELL | 1 | | |
| | FIBROBLASTS | FIBROBLAST | 1 | | |
| | FIBROBLAST | CELLULAR | 1 | | |
| | CELLULAR | | | | |
| file42 | CELLS | EGGS | 1 | | |
| | ASTROCYTES | ASTROCYTES | 1 | | |
| | CELL | CELL | 1 | | |
| | MICROGLIA | MICROGLIA | 1 | | |
| | NEURONS | NEURONS | 1 | | |
| | NEURONAL | NEURONAL | 1 | | |
| | MICROGLIAL | MICROGLIAL | 1 | | |
| | GLIAL | GLIAL | 1 | | |

| | | | | |
|---|---|---|---|---|
| | EGGS | CELLS | 1 | |
| file43 | ADIPOCYTES | ADIPOCYTES | 1 | |
| | STEM CELL | STEM CELL | 1 | |
| | MULTIPOTENTIAL PROGENITOR CELLS | MULTIPOTENTIAL PROGENITOR CELLS | 1 | |
| | GLIAL CELLS | GLIAL CELLS | 1 | |
| | OSTEOBLASTS | OSTEOBLASTS | 1 | |
| | CHONDROCYTES | CHONDROCYTES | 1 | |
| | STEM CELLS | STEM CELLS | 1 | |
| | STEM ... CELLS | CELL | 1 | |
| | CELL | NEURONS | 1 | |
| | NEURONS | MESENCHYMAL CELL | 1 | |
| | MESENCHYMAL CELL | MESENCHYMAL CELLS | 1 | |
| | MESENCHYMAL CELLS | HYPERTROPHIC CHONDROCYTES | 1 | |
| | HYPERTROPHIC CHONDROCYTES | OLIGODENDROCYTES | 1 | |
| | OLIGODENDROCYTES | CELLS | 1 | |
| | CELLS | OSTEOBLAST | 1 | |
| | OSTEOBLAST | ADIPOCYTE | 1 | |
| | ADIPOCYTE | FIBROBLASTS | 1 | |
| | FIBROBLASTS | FIBROBLAST | 1 | |
| | FIBROBLAST | CELLULAR | 1 | |
| | CELLULAR | GLIAL | | 1 |
| | OSTEOCLAST | OSTEOCLAST | 1 | |
| | OSTEOCLASTS | OSTEOCLASTS | 1 | |
| | STROMAL CELLS | STROMAL CELLS | 1 | |
| file44 | OOCYTES | FIBROBLASTS | 1 | |
| | CELLULAR | CELLULAR | 1 | |
| | OOCYTE | OOCYTE | 1 | |
| | PLATELET | SPERM | 1 | |
| | CELLS | CELLS | 1 | |
| | PLATELETS | PLATELETS | 1 | |
| | FIBROBLASTS | ERYTHROCYTES | 1 | |
| | ERYTHROCYTES | OOCYTES | 1 | |
| | SPERM | CELL | 1 | |
| | FIBROBLAST | FIBROBLAST | 1 | |
| | CELL | PLATELET | 1 | |
| | NEURONAL | KERATINOCYTES | 1 | |

| | | | | | |
|---|---|---|---|---|---|
| | EUKARYOTIC CELLS | NEURONAL | 1 | | |
| | STEMS CELLS | STEMS CELLS | 1 | | |
| file45 | | | | | |
| file46 | STEM ... CELLS | GRANULE CELLS | 1 | | 1 |
| | GRANULE CELLS | GRANULAR CELLS | 1 | | |
| | GLIAL CELL | CELLS | 1 | | 1 |
| | CELLS | NEURON | 1 | | |
| | NEURON | CELL | 1 | | |
| | CELL | PYRAMIDAL CELLS | 1 | | |
| | PYRAMIDAL CELLS | SPERMATOZOA | 1 | | |
| | GRANULAR CELLS | EGGS | 1 | | |
| | SPERMATOZOA | | | | |
| | EGGS | | | | |
| file47 | GERM CELLS | STEM CELLS | | 1 | |
| | GERM CELL | CELLULAR | 1 | | |
| | CELLS | GERM CELL | 1 | | |
| | STEM CELL | CELLS | 1 | | |
| | CELL | STEM CELL | 1 | | |
| | GERMCELLS | CELL | 1 | | |
| | MULTIPOTENT ... STEM CELLS | GERMCELLS | 1 | | 1 |
| | CELLULAR | GERM CELLS | 1 | | |
| | GERMLINE STEM CELLS | GERMLINE STEM CELLS | 1 | | |
| file48 | | | | | |
| file49 | RETINAL PHOTORECEPTOR CELLS | PHOTORECEPTORS | 1 | | |
| | PHOTORECEPTORS | HORIZONTAL CELLS | 1 | | |
| | PHOTORECEPTOR CELLS IN ... RETINA | CONE | 1 | | |
| | CONE PHOTORECEPTOR | RETINAL PHOTORECEPTORS | 1 | | |
| | HORIZONTAL CELLS | RODS | 1 | | |
| | CONE | PHOTORECEPTOR CELL | 1 | | |
| | RETINAL PHOTORECEPTORS | BIPOLAR CELLS | 1 | | |
| | RODS | CONES | 1 | | |
| | PHOTORECEPTOR CELL | CELL | 1 | | |
| | PHOTORECEPTORS IN ... RETINA | NEURONS | 1 | | 1 |

| | | | | | |
|---|---|---|---|---|---|
| | MÃ¼LLER GLIA | AMACRINE CELLS | 1 | | 1 |
| | BIPOLAR CELLS | CELLS | 1 | | |
| | CONES | AMACRINE | 1 | | |
| | MULTIPOTENT ... CELLS | GANGLION | | 1 | |
| | CELL | GLIA | | 1 | |
| | NEURONS | PHOTORECEPTOR | 1 | | |
| | AMACRINE CELLS | RETINAL PHOTORECEPTOR | 1 | | |
| | CELLS | GANGLION CELL | 1 | | |
| | AMACRINE | PINEALOCYTES | 1 | | |
| | CONE PHOTORECEPTORS | BIPOLAR | | 1 | |
| | PHOTORECEPTOR | PHOTORECEPTOR CELLS | 1 | | |
| | RETINAL PHOTORECEPTOR | ROD PHOTORECEPTORS | 1 | | |
| | GANGLION CELL | CONE PHOTORECEPTORS | 1 | | |
| | PINEALOCYTES | | | | |
| | GANGLION CELLS | | | | |
| | PHOTORECEPTOR CELLS | | | | |
| | ROD PHOTORECEPTOR | | | | |
| | ROD PHOTORECEPTORS | | | | |
| | ROD ... PHOTORECEPTORS | | | | |
| file50 | FIBROBLASTS | FIBROBLASTS | 1 | | |
| | HEPATOCYTES | STEM CELLS | 1 | | |
| | CELLULAR | CELLULAR | 1 | | |
| | CELLS | CELLS | 1 | | |
| | STEM CELL | STEM CELL | 1 | | |
| | CELL | CELL | 1 | | |
| | FIBROBLAST | FIBROBLAST | 1 | | |
| | STEM CELLS | HEPATOCYTES | 1 | | |
| file51 | STEM CELLS | TROPHOBLAST CELLS | 1 | | |
| | TROPHOBLAST CELLS | STEM CELLS | 1 | | |
| | NEURONAL | APOPTOTIC CELLS | 1 | | |
| | APOPTOTIC CELLS | CELLULAR | 1 | | |
| | EXTRAEMBRYONIC CELLS | CELLS | 1 | | 1 |
| | CELLS | STEM CELL | 1 | | |
| | CELLS IN ... EXTRAEMBRYONIC TISSUES | ERYTHROID | | 1 | |
| | STEM CELL | CELL | 1 | | |
| | CELL | NEURONS | 1 | | |

| | | | | | |
|---|---|---|---|---|---|
| | NEURONS | PROMYELOCYTIC | | 1 | |
| | CELLS OF ... ECTODERM | CONE | | 1 | 1 |
| | TROPHOBLAST ... CELLS | NEURONAL | 1 | | 1 |
| | CELLULAR | LYMPHOCYTES | 1 | | |
| | RED BLOOD CELLS | | | | 1 |
| | LYMPHOCYTES | | | | |
| | CELL IN ... EPIBLAST ... REGION | | | | 1 |
| | TROPHOBLAST ... CELL | | | | 1 |
| | DIPLOID ... CELL | | | | 1 |
| file52 | PANCREATIC BETA CELL | CELL | 1 | 1 | |
| | | PANCREATIC BETA CELL | | | |
| file53 | SERTOLI CELL | SERTOLI CELL | 1 | | |
| | SERTOLI | CHONDROCYTE | 1 | | |
| | ENDOTHELIAL CELL | CELLULAR | 1 | | |
| | CELLS | APOPTOTIC CELL | 1 | | |
| | CHONDROCYTE | SUPPORTING CELLS | 1 | | |
| | APOPTOTIC CELLS | SUPPORTING CELL | 1 | | |
| | GERM CELLS | APOPTOTIC CELLS | 1 | | |
| | SERTOLI CELLS | GERM CELL | 1 | | |
| | APOPTOTIC CELL | SERTOLI CELLS | 1 | | |
| | GERM CELL | CELL | 1 | | |
| | CELL | FOLLICLE CELLS | 1 | | |
| | FOLLICLE CELLS | FIBROBLAST | 1 | | |
| | FIBROBLAST | GERM CELLS | 1 | | |
| | PLATELET | VASCULAR ENDOTHELIAL CELLS | 1 | | 1 |
| | BLOOD CELLS | CELLS | 1 | | 1 |
| | POLYGONAL CELLS | SERTOLI | 1 | | 1 |
| | CELLULAR | | | | |
| | SUPPORTING CELLS | | | | |
| | VASCULAR ENDOTHELIAL CELLS | | | | |
| | SUPPORTING CELL | | | | |
| | ENDOTHELIAL ... CELLS | | | | 1 |
| file54 | STEM ... CELLS | FIBROBLASTS | 1 | | 1 |
| | CELL | CELL | 1 | | |
| | CELLS | CELLS | 1 | | |

| | | | | | |
|---|---|---|---|---|---|
| | FIBROBLASTS | FIBROBLASTIC | | 1 | |
| | FIBROBLASTIC CELL | FIBROBLASTIC CELL | 1 | | |
| file55 | OOCYTES | OOCYTES | 1 | | |
| | MACROPHAGES | STEM CELLS | 1 | | |
| | BASAL CELLS | BASAL CELLS | 1 | | |
| | EPITHELIAL CELLS | MACROPHAGES | 1 | | |
| | CELLS | EPITHELIAL CELLS | 1 | | |
| | CELLS OF ... STRATIFIED EPITHELIA | CELLS | 1 | | 1 |
| | STEM CELL | STEM CELL | 1 | | |
| | AMELOBLASTS | CELLS OF EPIDERMIS | 1 | | |
| | CELLULAR | ECTODERMAL CELLS | 1 | | |
| | CELL | CELL | 1 | | |
| | LYMPHOCYTES | LYMPHOCYTES | 1 | 1 | |
| | CELLS OF EPIDERMIS | KERATINOCYTES | 1 | | |
| | STEM CELLS | AMELOBLASTS | 1 | | |
| | EPITHELIAL CELL | CELLULAR | 1 | | |
| | KERATINOCYTES | EPITHELIAL CELL | 1 | | |
| | NEUTROPHILS | NEUTROPHILS | 1 | | |
| | ECTODERMAL CELLS | EPIDERMAL CELL | 1 | | |
| | EPIDERMAL CELL | | | | |
| file56 | FIBROBLASTS | FIBROBLASTS | 1 | | |
| | CELLULAR | CELLULAR | 1 | | |
| | CELLS | CELLS | 1 | | |
| | STEM CELL | STEM CELL | 1 | | |
| | CELL | CELL | 1 | | |
| | BLOOD CELLS | BLOOD CELLS | 1 | | |
| | STEM CELLS | STEM CELLS | 1 | | |
| | ENUCLEATING ... CELLS | | | | 1 |
| file57 | PLATELET | PLATELET | 1 | | |
| file58 | PHOTORECEPTORS | PHOTORECEPTORS | 1 | | |
| | CONE PHOTORECEPTOR | CONE PHOTORECEPTOR | 1 | | |
| | NEURON | NEURON | 1 | | |
| | GLIAL CELLS | GLIAL CELLS | 1 | | |
| | CONE | CONE | 1 | | |

| | | | | | |
|---|---|---|---|---|---|
| | RECEPTORAL ... NEURONS | INTERNEURONS | 1 | | 1 |
| | INTERNEURONS | PHOTOSENSORY NEURON | 1 | | |
| | RECEPTORAL NEURONS | CELL | 1 | | 1 |
| | PHOTOSENSORY NEURON | NEURONS | | | |
| | CELL | NEURONAL | 1 | | |
| | DOPAMINERGIC NEURONS | PHOTORECEPTOR NEURONS | 1 | | 1 |
| | NEURONS | GANGLION | | 1 | |
| | PHOTOSENSORY NEURONS IN ... RETINA | GLIA | 1 | | |
| | NEURONAL | PHOTORECEPTOR | 1 | | |
| | PRIMARY ... NEURON | ROD PHOTORECEPTOR NEURONAL | 1 | | 1 |
| | PHOTORECEPTOR NEURONS | CELLS | 1 | | |
| | CONE PHOTORECEPTORS | GANGLION CELL | 1 | | |
| | GLIA | GLIAL | 1 | | |
| | PHOTORECEPTOR | GANGLION CELLS | 1 | | |
| | ROD PHOTORECEPTOR NEURONAL | OOCYTES | 1 | | |
| | CELLS | ROD PHOTORECEPTOR | 1 | | |
| | GANGLION CELL | PHOTOSENSORY NEURONS | 1 | | |
| | GANGLION CELLS OF ... RETINA | ROD PHOTOSENSORY NEURONS | 1 | | |
| | PHOTORECEPTOR ... NEURONS OF ... RETINA | | | | 1 |
| | GANGLION CELLS | | | | |
| | OOCYTES | | | | |
| | PHOTOSENSORY NEURONS OF ... RETINA | | | | 1 |
| | ROD PHOTORECEPTOR | | | | |
| | PRIMARY ... NEURONS | | | | |
| | PHOTOSENSORY NEURONS | | | | |
| | ROD PHOTOSENSORY NEURONS | | | | |
| file59 | NEURAL CREST CELL | APOPTOTIC CELLS | 1 | | |
| | EPITHELIAL ... CELL | NEURAL CREST CELL | 1 | | 1 |
| | NEURAL CREST CELLS | CELLS | 1 | | |
| | CELLS | SMOOTH MUSCLE CELLS | 1 | | |
| | APOPTOTIC CELLS | SMOOTH MUSCLE CELL | 1 | | |
| | SMOOTH MUSCLE CELL | CELL | 1 | | |
| | CELL | STEM CELLS | 1 | | |
| | STEM CELLS | NEURAL CREST CELLS | 1 | | |
| | SMOOTH MUSCLE CELLS | MESENCHYMAL CELL | 1 | | |
| | MESENCHYMAL CELL | MUSCLE CELL | 1 | | |
| | NC ... CELL | NCCS | 1 | | 1 |

| | | | | |
|---|---|---|---|---|
| | NCCS | | | |
| file60 | FIBROBLASTS | CELL | 1 | |
| | STEM CELLS | CELLULAR | 1 | |
| | CELLS | CELLS | 1 | |
| | STEM CELL | STEM CELL | 1 | |
| | CELL | FIBROBLASTS | 1 | |
| | FIBROBLAST | FIBROBLAST | 1 | |
| | PROMYELOCYTIC | PROMYELOCYTIC | 1 | |
| | CELLULAR | STEM CELLS | 1 | |
| file61 | OSTEOBLASTIC | OSTEOBLASTIC | 1 | |
| | APOPTOTIC CELLS | OSTEOPROGENITOR CELLS | 1 | |
| | CHONDROCYTE | HYPERTROPHIC CHONDROCYTES | 1 | |
| | OSTEOCLASTS | OSTEOCLASTS | 1 | |
| | CELLS | CELLS | 1 | |
| | OSTEOBLAST CELLS | APOPTOTIC CELLS | 1 | 1 |
| | FIBROBLASTIC | FIBROBLASTIC | 1 | |
| | OSTEOBLAST | OSTEOBLAST | 1 | |
| | CELL | CELL | 1 | |
| | FIBROBLAST | FIBROBLAST | 1 | |
| | OSTEOBLASTS | OSTEOBLASTS | 1 | |
| | OSTEOPROGENITOR CELLS | CHONDROCYTES | 1 | |
| | OSTEOPROGENITORS | OSTEOPROGENITORS | 1 | |
| | CHONDROCYTES | OSTEOCLAST | 1 | |
| | OSTEOCLAST | FIBROBLASTIC CELLS | 1 | |
| | FIBROBLASTIC CELLS | CHONDROCYTE | 1 | |
| | HYPERTROPHIC CHONDROCYTES | | | |
| file62 | CELLS | CELLS | 1 | |
| | REGULATORY T CELL | REGULATORY T CELL | 1 | |
| | T CELL | T CELL | 1 | |
| | CELL | CELL | 1 | |
| | BLOOD CELLS | BLOOD CELLS | 1 | |
| | LYMPHOCYTES | LYMPHOCYTES | 1 | |
| | LYMPHOCYTE | LYMPHOCYTE | 1 | |
| file63 | EPITHELIAL CELLS | GLIAL CELL | 1 | 1 |

| | | | | | |
|---|---|---|---|---|---|
| | GLIAL CELL | CELLS | 1 | | |
| | CELLS | STROMAL CELL | 1 | | |
| | STROMAL CELL | CELL | 1 | | |
| | CELL | STROMAL CELLS | 1 | | |
| | STROMAL CELLS | GLIAL | | 1 | |
| file64 | GAMETES | GAMETES | 1 | | |
| | SERTOLI CELLS | SERTOLI CELLS | 1 | | |
| | SERTOLI CELL | SERTOLI CELL | 1 | | |
| | MULTINUCLEATE CELLS | MULTINUCLEATE CELLS | 1 | | |
| | SPERMATOGONIA | SPERMATOGONIA | 1 | | |
| | MULTINUCLEATED CELLS | MULTINUCLEATED CELLS | 1 | | |
| | GONOCYTES | GONOCYTES | 1 | | |
| | MALE GERM CELL | OOCYTE | 1 | | |
| | STEM ... CELLS | CELL | 1 | | 1 |
| | OOCYTE | PGCS | 1 | | |
| | GERM CELL | GERM CELL | 1 | | |
| | CELL | MALE GERM CELL | 1 | | |
| | GERM-CELL | GERM-CELL | 1 | | |
| | SPERMATOCYTES | SPERMATOZOA | 1 | | |
| | SERTOLI | SERTOLI | 1 | | |
| | SPERMATOCYTE | SPERMATOCYTE | 1 | | |
| | APOPTOTIC CELLS | APOPTOTIC CELLS | 1 | | |
| | CELLS | CELLS | 1 | | |
| | LEYDIG CELLS | CELLULAR | 1 | | |
| | CELLULAR | SPERMATOCYTES | 1 | | |
| | SPERMATOZOA | SPERMATIDS | 1 | | |
| | SPERMATIDS | OOCYTES | 1 | | |
| | OOCYTES | SPERMATID | 1 | | |
| | SPERMATID | MALE GERM CELLS | 1 | | |
| | MALE GERM CELLS | ZYGOTE | 1 | | |
| | ZYGOTE | SPERM | 1 | | |
| | SPERM | GERM CELLS | 1 | | |
| | GERM CELLS | | | | |
| | PRIMARY SPERMATOCYTES | | | | 1 |
| file65 | LYMPHOCYTES | LYMPHOCYTES | 1 | | |
| | CELLS | CELLS | 1 | | |

| | | | | |
|---|---|---|---|---|
| | PURKINJE CELLS | PURKINJE CELLS | 1 | |
| file66 | BIPOLAR CELLS IN ... RETINA | PHOTORECEPTORS | 1 | 1 |
| | FIBROBLAST | AMACRINE NEURONS | 1 | |
| | PHOTORECEPTORS | NEURON | 1 | |
| | AMACRINE NEURONS | CNS NEURONS | 1 | |
| | NEURON | GANGLION NEURONS | 1 | |
| | CNS NEURONS | BIPOLAR NEURONS | 1 | |
| | GANGLION ... CELLS IN ... RETINA | CONE | 1 | |
| | ROD BIPOLAR ... CELLS | RODS | 1 | |
| | GANGLION NEURONS | CHOLINERGIC NEURONS | 1 | |
| | CONE | INTERNEURONS | 1 | |
| | ROD ... BIPOLAR CELLS | BIPOLAR CELLS | 1 | |
| | GANGLION ... CELLS | CONES | 1 | |
| | RODS | ADIPOCYTES | 1 | |
| | CHOLINERGIC NEURONS | CELL | 1 | |
| | INTERNEURONS | AMACRINE CELLS | 1 | |
| | MÃ¼LLER GLIA CELL | ROD BIPOLAR CELLS | 1 | |
| | AMACRINE ... CELLS | NEURONAL | 1 | 1 |
| | MÃ¼LLER CELLS | AMACRINE | 1 | |
| | CONES | GANGLION | 1 | 1 |
| | HORIZONTAL ... NEURONS | ON BIPOLAR CELLS | 1 | |
| | GANGION CELLS | GLIA | 1 | |
| | CELL | PHOTORECEPTOR | 1 | |
| | BIPOLAR CELL | APOPTOTIC CELLS | 1 | |
| | ROD BIPOLAR CELLS | CELLS | 1 | |
| | AMACRINE CELLS | GANGLION CELL | 1 | |
| | NEURONAL | FIBROBLASTS | 1 | |
| | RETINAL ... ROD CELLS | FIBROBLAST | 1 | |
| | ROD ... CELLS IN ... RETINA | NEURONS | 1 | 1 |
| | AMACRINE | GLIAL | 1 | 1 |
| | GANGLION | GANGLION CELLS | 1 | |
| | ON BIPOLAR CELLS | BIPOLARS | 1 | |
| | GANGLION ... CELL | BIPOLAR CELL | 1 | |
| | ADIPOCYTES | BIPOLAR | 1 | 1 |
| | PHOTORECEPTOR | ROD PHOTORECEPTORS | 1 | |
| | APOPTOTIC CELLS | AMACRINE CELL | 1 | |
| | GANGLION CELL | ENTEROCYTES | 1 | |

| | | | | | |
|---|---|---|---|---|---|
| | CELLS | NEURONAL CELL | 1 | | |
| | FIBROBLASTS | ROD BIPOLAR | 1 | | |
| | CONE BIPOLAR CELLS | | | | 1 |
| | BIPOLAR | | | | |
| | BIPOLAR CELLS | | | | |
| | NEURONS | | | | |
| | GLIAL | | | | |
| | GANGLION CELLS | | | | |
| | ROD ... CELLS | | | | |
| | BIPOLARS | | | | |
| | HORIZONTAL ... CELLS | | | | 1 |
| | ROD ... CELL | | | | 1 |
| | RETINAL ... GANGLION ... CELLS | | | | |
| | BIPOLAR NEURONS | | | | |
| | ROD PHOTORECEPTORS | | | | |
| | AMACRINE CELL | | | | |
| | ENTEROCYTES | | | | |
| | RETINAL BIPOLAR ... CELLS | | | | 1 |
| file67 | SPERMATOCYTE | SPERMATOCYTE | 1 | | |
| | OOCYTE | OOCYTES | 1 | | |
| | GAMETES | STEM CELL | 1 | | |
| | STEM CELL | GAMETES | 1 | | |
| | EGGS | SPERM | 1 | | |
| | GERM CELL | EGGS | 1 | | |
| | OOCYTES | GERM CELL | 1 | | |
| | CELL | CELL | 1 | | |
| | SPERM | SPERMATOGONIA | 1 | | |
| | SPERMATOGONIA | OOCYTE | 1 | | |
| | SPERMATIDS | SPERMATOCYTES | 1 | | |
| | STEM CELLS | CELLS | 1 | | |
| | SPORE | SPERMATIDS | 1 | | 1 |
| | SPERMATOCYTES | | | | |
| | CELLS | | | | |
| | GAMETE | | | | 1 |
| | *** | | | | |
| Total | | | 730 | 47 | 93 |
| | | Accuracy | 0,94 | | |

| | |
|---|---|
| Recall | 0,89 |
| F-Measure | 0,91 |
