## Supplementary_Excel_file_S2 Same_dictionary_OntoC_Annot for "OntoContext, a new python package for gene contextualization based on the annotation of biomedical texts"

| Num file | Reference CRAFT 1.0 corpus Annotation | Annotation Ontocontext | NCBO Annotator Annotation | TP_OntoC | FP_OntoC | FN_OntoC | TP_Anno | FP_Anno | FN_Anno |
|---|---|---|---|---|---|---|---|---|---|
| file2 | CELL | CELL | CELL | 1 | | | 1 | | |
| | STEM ... CELLS | CELLS | | 1 | | 1 | | | 1 |
| | CELLS | | | | | | | | 1 |
| file3 | OLIGODENDROCYTE | | OLIGODENDROCYTE | | | 1 | 1 | | |
| | CELLULAR | | | | | 1 | | | 1 |
| file4 | CELLS | CELLS | CELL | 1 | | | 1 | | 1 |
| | CELL | CELL | | 1 | | | | | |
| | BLASTOMERES | BLASTOMERES | | 1 | | | | | 1 |
| | ENDODERM CELLS | ENDODERM CELLS | | 1 | | | | | 1 |
| | STEM CELLS | STEM CELLS | | 1 | | | | | 1 |
| | EUKARYOTIC CELLS | EUKARYOTIC CELLS | | 1 | | | | | 1 |
| file5 | B LYMPHOCYTES | CELL | CELL | 1 | | 1 | 1 | | 1 |
| | CELL | CELLS | EPITHELIAL CELL | 1 | | | 1 | | |
| | CELLS | EPITHELIAL CELL | LENS EPITHELIAL CELL | 1 | | | | 1 | 1 |
| | CELLULAR | EPITHELIAL CELLS | MACROPHAGE | 1 | | 1 | 1 | | |
| | EPITHELIAL CELL | LENS FIBER CELLS | MUSCLE CELL | | 1 | | 1 | | |
| | EPITHELIAL CELLS | LYMPHOCYTES | | | 1 | | | | 1 |
| | MACROPHAGE | MACROPHAGE | | 1 | | | | | |
| | MUSCLE CELL | MUSCLE CELL | | 1 | | | | | |
| | OSTEOBLASTS | OSTEOBLASTS | | 1 | | | | | 1 |
| file6 | CELL | CELL | CELL | 1 | | | 1 | | |
| | CELLS | CELLS | NEURON | 1 | | | 1 | | 1 |
| | CELLULAR | EAR HAIR CELL | OUTER HAIR CELL | 1 | | 1 | 1 | | 1 |
| | DEITERS' CELLS | HAIR CELL | SENSORY NEURON | 1 | | 1 | | | 1 |
| | HAIR CELL | HAIR CELLS | VESTIBULAR HAIR CELL | 1 | | | 1 | | 1 |
| | HAIR CELLS | INNER EAR HAIR CELL | | 1 | | | | | 1 |
| | INNER ... HAIR CELLS | INNER HAIR CELLS | | 1 | | | | | 1 |
| | INNER HAIR CELL | MOTONEURONS | | 1 | | | | | 1 |
| | INNER HAIR CELLS | NEURON | | 1 | | | | | |
| | MOTONEURONS | NEURONS | | 1 | | | | | 1 |
| | NEURON | OUTER HAIR CELL | | 1 | | | | | |
| | NEURON AFFERENTS | OUTER HAIR CELLS | | 1 | | 1 | | | 1 |
| | NEURONAL | PILLAR CELLS | | 1 | | 1 | | | 1 |
| | NEURONS | PRIMARY NEURONS | | 1 | | | | | 1 |
| | OUTER HAIR CELL | SENSORY NEURON | | 1 | | | | | |
| | OUTER HAIR CELLS | SENSORY NEURONS | | 1 | | | | | 1 |
| | PILLAR CELLS | | | | | | | | 1 |
| | PRIMARY NEURONS | | | | | | | | 1 |
| | SENSORY NEURON | | | | | | | | 1 |
| | SENSORY NEURONS | | | | | | | | 1 |
| | SUPPORTING CELL | | | | | 1 | | | 1 |
| | SUPPORTING CELLS | | | | | 1 | | | 1 |
| | VESTIBULAR HAIR CELL | | | | | 1 | | | |
| file38 | CELL | CELL | CELL | 1 | | | 1 | | |
| | CELLS | CELLS | SPERM | 1 | | | 1 | | 1 |
| | MYOCYTE | MYOCYTE | | 1 | | | | | 1 |
| | MYOCYTES | MYOCYTES | | 1 | | | | | 1 |
| | SPERM | SPERM | | 1 | | | 1 | | |
| | STEM ... CELLS | | | | | | | | 1 |
| | STEM CELLS | | | | | 1 | | | 1 |

| | OntoContext | Annotator |
|---|---|---|
| Accuracy | 0,79 | 0,82 |
| Recall | 0,59 | 0,23 |
| F-Measure | 0,68 | 0,36 |

| | | | | | | | | | |
|---|---|---|---|---|---|---|---|---|---|
| file39 | CELL | CELL | CELL | 1 | | | 1 | | |
| | CELLS | CELLS | OLFACTORY BULB | 1 | | | 1 | | 1 |
| | CELLULAR | CORTICAL NEURONS | | | 1 | 1 | | | 1 |
| | GRANULE CELLS | FOREBRAIN NEURONS | | | 1 | | | | 1 |
| | NEURONAL | GRANULE CELLS | | 1 | | 1 | | | 1 |
| | NEURONS | HIPPOCAMPAL NEURONS | | | 1 | | | | 1 |
| | OLFACTORY SENSORY NEURONS | HIPPOCAMPAL PYRAMIDAL NEURONS | | | 1 | 1 | | | 1 |
| | OOCYTES | HIPPOCAMPUS | | | 1 | | | | 1 |
| | OUTPUT NEURONS | MITRAL CELLS | | | 1 | | | | 1 |
| | PRINCIPAL ... NEURONS | NEURONS | | 1 | | 1 | | | 1 |
| | PRINCIPAL NEURONS | OLFACTORY BULB | | | 1 | | | | 1 |
| | PYRAMIDAL CELLS | OLFACTORY SENSORY NEURONS | | 1 | | | | | 1 |
| | PYRAMIDAL NEURONS | OOCYTES | | 1 | | | | | 1 |
| | | OUTPUT NEURONS | | 1 | | | | | |
| | | PYRAMIDAL CELLS | | 1 | | | | | |
| | | PYRAMIDAL NEURONS | | 1 | | | | | |
| | | SENSORY NEURONS | | | 1 | | | | |
| file40 | CELL | CELL | CELL | 1 | | | 1 | | |
| | CELLS | CELLS | EMBRYONIC FIBROBLAST | 1 | | | | 1 | 1 |
| | CELLULAR | CULTURED CELLS | FIBROBLAST | | 1 | 1 | | 1 | 1 |
| | FIBROBLAST CELL | EMBRYONIC FIBROBLAST | HEPATOCYTE | | 1 | 1 | 1 | | 1 |
| | FIBROBLASTS | EMBRYONIC FIBROBLASTS | | | 1 | | | | 1 |
| | GLIAL CELLS | FIBROBLAST | | 1 | | | | | 1 |
| | HEPATOCYTE | FIBROBLASTS | | 1 | | | | | 1 |
| | | GLIAL CELLS | | 1 | | | | | |
| | | HEPATOCYTE | | 1 | | | | | |
| file41 | B CELL | BRONCHIAL EPITHELIAL CELLS | B CELL | | 1 | 1 | 1 | | |
| | CELL | CELL | CELL | 1 | | | 1 | | |
| | CELLS | CELLS | FIBROBLAST | 1 | | | 1 | | |
| | CELLULAR | EPIDERMAL CELLS | | 1 | | 1 | | | 1 |
| | EPIDERMAL CELLS | EPITHELIAL CELLS | | 1 | | | | | 1 |
| | EPITHELIAL CELLS | FIBROBLAST | | 1 | | | | | 1 |
| | FIBROBLAST | FIBROBLASTS | | 1 | | | | | |
| | FIBROBLASTS | | | | | | | | 1 |
| file42 | ASTROCYTES | ACTIVATED ASTROCYTES | | | 1 | | | | 1 |
| | CELL | ACTIVATED MICROGLIA | | | 1 | | | | 1 |
| | CELLS | ASTROCYTES | | 1 | | | | | 1 |
| | EGGS | CELL | | 1 | | 1 | | | 1 |
| | GLIAL | CELLS | | 1 | | 1 | | | 1 |
| | MICROGLIA | CORTICAL NEURONS | | | 1 | | | | 1 |
| | MICROGLIAL | HIPPOCAMPUS | | | 1 | | | | 1 |
| | NEURONAL | MICROGLIA | | 1 | | 1 | | | 1 |
| | NEURONS | NEURONS | | 1 | | | | | 1 |
| file43 | ADIPOCYTE | ADIPOCYTE | BONE CELL | 1 | | | | 1 | 1 |
| | ADIPOCYTES | ADIPOCYTES | CELL | 1 | | | 1 | | 1 |
| | CELL | BONE CELL | EMBRYONIC CELL | | 1 | | | 1 | |
| | CELLS | BONE CELLS | EMBRYONIC FIBROBLAST | | 1 | | | 1 | 1 |
| | CELLULAR | BONE MARROW STROMAL CELLS | FIBROBLAST | | 1 | 1 | 1 | | 1 |
| | CHONDROCYTES | CELL | MESENCHYMAL CELL | 1 | | | 1 | | 1 |
| | FIBROBLAST | CELLS | MESENCHYMAL STEM CELL | 1 | | | | 1 | |
| | FIBROBLASTS | CFU-F | OSTEOBLAST | | 1 | | 1 | | |
| | GLIAL CELLS | CHONDROCYTES | OSTEOCLAST | 1 | | | 1 | | 1 |
| | HYPERTROPHIC CHONDROCYTES | EMBRYONIC CELL | PRECURSOR CELL | | 1 | | | 1 | 1 |

| | | | | | | | | | |
|---|---|---|---|---|---|---|---|---|---|
| | MESENCHYMAL CELL | EMBRYONIC FIBROBLAST | STEM CELL | | 1 | | 1 | | |
| | MESENCHYMAL CELLS | FIBROBLAST | | 1 | | | | | 1 |
| | MULTIPOTENTIAL PROGENITOR CELLS | FIBROBLASTS | | 1 | | 1 | | | 1 |
| | NEURONS | GLIAL CELLS | | 1 | | | | | 1 |
| | OLIGODENDROCYTES | HYPERTROPHIC CHONDROCYTES | | 1 | | | | | 1 |
| | OSTEOBLAST | MESENCHYMAL CELL | | 1 | | | | | |
| | OSTEOBLASTS | MESENCHYMAL CELLS | | 1 | | | | | 1 |
| | OSTEOCLAST | MESENCHYMAL PRECURSOR CELL | | | 1 | | | | |
| | OSTEOCLASTS | MESENCHYMAL STEM CELL | | 1 | | | | | 1 |
| | STEM ... CELLS | NEURONS | | 1 | | | | | |
| | STEM CELL | OLIGODENDROCYTES | | 1 | | | | | |
| | STEM CELLS | OSTEOBLAST | | 1 | | | | | 1 |
| | STROMAL CELLS | OSTEOBLASTS | | 1 | | | | | 1 |
| | | OSTEOCLAST | | 1 | | | | | |
| | | OSTEOCLASTS | | 1 | | | | | |
| | | PRECURSOR CELL | | | 1 | | | | |
| | | STEM CELL | | 1 | | | | | |
| | | STEM CELLS | | 1 | | | | | |
| | | STROMAL CELLS | | 1 | | | | | |
| file44 | CELL | CELL | CELL | 1 | | | 1 | | |
| | CELLS | CELLS | EMBRYONIC FIBROBLAST | 1 | | | | 1 | 1 |
| | CELLULAR | EMBRYONIC FIBROBLASTS | FIBROBLAST | | 1 | 1 | 1 | | |
| | ERYTHROCYTES | ERYTHROCYTES | OOCYTE | 1 | | | 1 | | |
| | EUKARYOTIC CELLS | FIBROBLAST | PLATELET | 1 | | 1 | 1 | | 1 |
| | FIBROBLAST | FIBROBLASTS | SPERM | 1 | | | 1 | | |
| | FIBROBLASTS | KERATINOCYTES | | | 1 | | | | 1 |
| | NEURONAL | OOCYTE | | 1 | | 1 | | | 1 |
| | OOCYTE | OOCYTES | | 1 | | | | | |
| | OOCYTES | PLATELET | | 1 | | | | | 1 |
| | PLATELET | PLATELETS | | 1 | | | | | |
| | PLATELETS | SPERM | | 1 | | | | | 1 |
| | SPERM | | | | | | | | |
| | STEMS CELLS | | | | | 1 | | | 1 |
| file45 | | | | | | | | | |
| file46 | CELL | CELL | CELL | 1 | | | 1 | | |
| | CELLS | CELLS | GLIAL CELL | 1 | | | 1 | | 1 |
| | EGGS | CEREBELLAR GRANULE CELLS | NEURON | | 1 | 1 | 1 | | 1 |
| | GLIAL CELL | GRANULE CELLS | | 1 | | 1 | | | |
| | GRANULAR CELLS | HIPPOCAMPUS | | | 1 | 1 | | | 1 |
| | GRANULE CELLS | NEURAL CELL | | | 1 | | | | 1 |
| | NEURON | NEURON | | 1 | | | | | |
| | PYRAMIDAL CELLS | PYRAMIDAL CELLS | | 1 | | | | | |
| | SPERMATOZOA | | | | | 1 | | | 1 |
| | STEM ... CELLS | | | | | 1 | | | 1 |
| file47 | CELL | CELL | | 1 | | | | | 1 |
| | CELLS | CELLS | | 1 | | | | | 1 |
| | CELLULAR | GERM CELL | | 1 | | 1 | | | 1 |
| | GERM CELL | GERM CELLS | | 1 | | | | | 1 |
| | GERM CELLS | GERMLINE STEM CELLS | | 1 | | | | | 1 |
| | GERMLINE STEM CELLS | SOMATIC CELLS | | | 1 | | | | 1 |
| | MULTIPOTENT ... STEM CELLS | STEM CELL | | 1 | | 1 | | | 1 |
| | STEM CELL | STEM CELLS | | | 1 | | | | 1 |

| | | | | | | | | | |
|---|---|---|---|---|---|---|---|---|---|
| file48 | | | | | | | | | |
| file49 | AMACRINE | AMACRINE CELLS | CELL | 1 | | 1 | 1 | | 1 |
| | AMACRINE CELLS | CELL | PHOTORECEPTOR CELL | 1 | | | 1 | | 1 |
| | BIPOLAR CELLS | CELLS | | 1 | | 1 | | | 1 |
| | CELL | HORIZONTAL CELLS | | 1 | | | | | |
| | CELLS | NEURONS | | 1 | | | | | 1 |
| | CONE | PHOTORECEPTOR CELL | | 1 | | 1 | | | 1 |
| | CONE PHOTORECEPTOR | PHOTORECEPTOR CELLS | | 1 | | 1 | | | 1 |
| | CONE PHOTORECEPTORS | PINEALOCYTES | | 1 | | 1 | | | 1 |
| | CONES | | | | | 1 | | | 1 |
| | GANGLION CELL | | | | | 1 | | | 1 |
| | GANGLION CELLS | | | | | 1 | | | 1 |
| | HORIZONTAL CELLS | | | | | | | | 1 |
| | MÃ¼LLER GLIA | | | | | 1 | | | 1 |
| | MULTIPOTENT ... CELLS | | | | | 1 | | | 1 |
| | NEURONS | | | | | | | | 1 |
| | PHOTORECEPTOR | | | | | 1 | | | 1 |
| | PHOTORECEPTOR CELL | | | | | | | | 1 |
| | PHOTORECEPTOR CELLS | | | | | | | | 1 |
| | PHOTORECEPTOR CELLS IN ... RETINA | | | | | 1 | | | 1 |
| | PHOTORECEPTORS | | | | | 1 | | | 1 |
| | PHOTORECEPTORS IN ... RETINA | | | | | 1 | | | 1 |
| | PINEALOCYTES | | | | | | | | 1 |
| | RETINAL PHOTORECEPTOR | | | | | 1 | | | 1 |
| | RETINAL PHOTORECEPTOR CELLS | | | | | 1 | | | 1 |
| | RETINAL PHOTORECEPTORS | | | | | 1 | | | 1 |
| | ROD ... PHOTORECEPTORS | | | | | 1 | | | 1 |
| | ROD PHOTORECEPTOR | | | | | 1 | | | 1 |
| | ROD PHOTORECEPTORS | | | | | 1 | | | 1 |
| | RODS | | | | | 1 | | | 1 |
| file50 | CELL | CELL | CELL | 1 | | | 1 | | |
| | CELLS | CELLS | EMBRYONIC FIBROBLAST | 1 | | | | 1 | 1 |
| | CELLULAR | EMBRYONIC FIBROBLASTS | EMBRYONIC STEM CELL | | 1 | 1 | | 1 | 1 |
| | FIBROBLAST | EMBRYONIC STEM CELL | FIBROBLAST | | 1 | | 1 | | |
| | FIBROBLASTS | EMBRYONIC STEM CELLS | STEM CELL | | 1 | | | | 1 |
| | HEPATOCYTES | FIBROBLAST | | 1 | | | | | 1 |
| | STEM CELL | FIBROBLASTS | | 1 | | | 1 | | |
| | STEM CELLS | HEPATOCYTES | | 1 | | | | | 1 |
| | | STEM CELL | | 1 | | | | | |
| | | STEM CELLS | | 1 | | | | | |
| file51 | APOPTOTIC CELLS | CELL | CELL | 1 | | 1 | 1 | | 1 |
| | CELL | CELLS | STEM CELL | 1 | | | 1 | | |
| | CELL IN ... EPIBLAST ... REGION | HIPPOCAMPUS | TROPHOBLAST GIANT CELL | | 1 | 1 | | 1 | 1 |
| | CELLS | LYMPHOCYTES | | 1 | | | | | 1 |
| | CELLS IN ... EXTRAEMBRYONIC TISSUES | NEURONS | | 1 | | 1 | | | 1 |
| | CELLS OF ... ECTODERM | STEM CELL | | 1 | | 1 | | | 1 |
| | CELLULAR | STEM CELLS | | 1 | | 1 | | | 1 |
| | DIPLOID ... CELL | TROPHOBLAST CELLS | | 1 | | 1 | | | 1 |
| | EXTRAEMBRYONIC CELLS | TROPHOBLAST GIANT CELLS | | | 1 | 1 | | | 1 |
| | LYMPHOCYTES | | | | | | | | 1 |
| | NEURONAL | | | | | 1 | | | 1 |
| | NEURONS | | | | | | | | 1 |
| | RED BLOOD CELLS | | | | | 1 | | | 1 |
| | STEM CELL | | | | | | | | |

| | | | | | | | | | |
|---|---|---|---|---|---|---|---|---|---|
| | STEM CELLS | | | | | | | | 1 |
| | TROPHOBLAST ... CELL | | | | | 1 | | | 1 |
| | TROPHOBLAST ... CELLS | | | | | | | | |
| | TROPHOBLAST CELLS | | | | | | | | 1 |
| file52 | PANCREATIC BETA CELL | BETA CELL | CELL | | 1 | | | 1 | |
| | | CELL | | | 1 | | | | |
| | | PANCREATIC BETA CELL | | 1 | | | | | |
| file53 | APOPTOTIC CELL | CELL | CELL | 1 | | 1 | 1 | | 1 |
| | APOPTOTIC CELLS | CELLS | CHONDROCYTE | 1 | | 1 | 1 | | 1 |
| | BLOOD CELLS | CHONDROCYTE | ENDOTHELIAL CELL | 1 | | 1 | 1 | | 1 |
| | CELL | ENDOTHELIAL CELLS | FIBROBLAST | 1 | | | 1 | | |
| | CELLS | FIBROBLAST | GERM CELL | 1 | | | 1 | | 1 |
| | CELLULAR | FOLLICLE CELLS | PLATELET | 1 | | 1 | 1 | | 1 |
| | CHONDROCYTE | GERM CELL | SERTOLI CELL | 1 | | | 1 | | |
| | ENDOTHELIAL ... CELLS | GERM CELLS | SOMATIC CELL | 1 | | | 1 | | 1 |
| | ENDOTHELIAL CELL | PRECURSOR CELLS | | | 1 | 1 | | | 1 |
| | FIBROBLAST | SERTOLI CELL | | 1 | | | | | 1 |
| | FOLLICLE CELLS | SERTOLI CELLS | | 1 | | | | | 1 |
| | GERM CELL | SOMATIC CELL | | | 1 | | | | 1 |
| | GERM CELLS | VASCULAR ENDOTHELIAL CELLS | | 1 | | | | | 1 |
| | PLATELET | | | | | 1 | | | 1 |
| | POLYGONAL CELLS | | | | | 1 | | | 1 |
| | SERTOLI | | | | | 1 | | | 1 |
| | SERTOLI CELL | | | | | | | | |
| | SERTOLI CELLS | | | | | | | | 1 |
| | SUPPORTING CELL | | | | | 1 | | | 1 |
| | SUPPORTING CELLS | | | | | 1 | | | 1 |
| | VASCULAR ENDOTHELIAL CELLS | | | | | | | | 1 |
| file54 | CELL | CELL | CELL | 1 | | | 1 | | |
| | CELLS | CELLS | SOMATIC CELL | 1 | | | | 1 | 1 |
| | FIBROBLASTIC CELL | EMBRYONIC FIBROBLASTS | | | 1 | 1 | | | 1 |
| | FIBROBLASTS | FIBROBLASTS | | 1 | | | | | 1 |
| | STEM ... CELLS | SOMATIC CELL | | | 1 | 1 | | | 1 |
| | | SOMATIC CELLS | | | 1 | | | | |
| file55 | AMELOBLASTS | AMELOBLASTS | CELL | 1 | | | 1 | | 1 |
| | BASAL CELLS | BASAL CELLS | EPIDERMAL CELL | 1 | | | 1 | | 1 |
| | CELL | CELL | EPITHELIAL CELL | 1 | | | 1 | | |
| | CELLS | CELLS | STEM CELL | 1 | | | 1 | | |
| | CELLS OF ... STRATIFIED EPITHELIA | ECTODERMAL CELLS | THYMOCYTE | 1 | | 1 | | 1 | 1 |
| | CELLS OF EPIDERMIS | EMBRYONIC CELLS | | | 1 | 1 | | | 1 |
| | CELLULAR | EPIDERMAL CELL | | 1 | | 1 | | | 1 |
| | ECTODERMAL CELLS | EPITHELIAL CELL | | 1 | | | | | 1 |
| | EPIDERMAL CELL | EPITHELIAL CELLS | | 1 | | | | | |
| | EPITHELIAL CELL | KERATINOCYTES | | 1 | | | | | |
| | EPITHELIAL CELLS | LYMPHOCYTES | | 1 | | | | | 1 |
| | KERATINOCYTES | MACROPHAGES | | 1 | | | | | 1 |
| | LYMPHOCYTES | NEUTROPHILS | | 1 | | | | | 1 |
| | MACROPHAGES | OOCYTES | | 1 | | | | | 1 |
| | NEUTROPHILS | STEM CELL | | 1 | | | | | 1 |
| | OOCYTES | STEM CELLS | | 1 | | | | | 1 |
| | STEM CELL | THYMIC EPITHELIAL CELLS | | | 1 | | | | |
| | STEM CELLS | THYMOCYTE | | | 1 | | | | 1 |
| | | THYMOCYTES | | | 1 | | | | |

| | | | | | | | | | |
|---|---|---|---|---|---|---|---|---|---|
| file56 | BLOOD CELLS | BLOOD CELLS | CELL | 1 | | | 1 | | 1 |
| | CELL | CELL | EMBRYONIC STEM CELL | 1 | | | | 1 | |
| | CELLS | CELLS | STEM CELL | 1 | | | 1 | | 1 |
| | CELLULAR | EMBRYONIC FIBROBLASTS | | | 1 | 1 | | | 1 |
| | ENUCLEATING ... CELLS | EMBRYONIC STEM CELL | | | 1 | 1 | | | 1 |
| | FIBROBLASTS | EMBRYONIC STEM CELLS | | | 1 | | | | 1 |
| | STEM CELL | FIBROBLASTS | | 1 | | | | | |
| | STEM CELLS | STEM CELL | | 1 | | | | | 1 |
| | | STEM CELLS | | 1 | | | | | |
| file57 | PLATELET | PLATELET | PLATELET | 1 | | | 1 | | |
| file58 | CELL | CELL | CELL | 1 | | | 1 | | |
| | CELLS | CELLS | NEURON | 1 | | | 1 | | 1 |
| | CONE | CEREBRAL CORTEX NEURONS | | | 1 | 1 | | | 1 |
| | CONE PHOTORECEPTOR | CORTICAL NEURONS | | | 1 | 1 | | | 1 |
| | CONE PHOTORECEPTORS | GLIAL CELLS | | 1 | | 1 | | | 1 |
| | DOPAMINERGIC NEURONS | HIPPOCAMPAL NEURONS | | | 1 | 1 | | | 1 |
| | GANGLION CELL | HIPPOCAMPUS | | | 1 | 1 | | | 1 |
| | GANGLION CELLS | INTERNEURONS | | 1 | | 1 | | | 1 |
| | GANGLION CELLS OF ... RETINA | NEURON | | 1 | | 1 | | | 1 |
| | GLIA | NEURONS | | 1 | | 1 | | | 1 |
| | GLIAL CELLS | OOCYTES | | 1 | | | | | 1 |
| | INTERNEURONS | | | | | | | | 1 |
| | NEURON | | | | | | | | |
| | NEURONAL | | | | | 1 | | | 1 |
| | NEURONS | | | | | | | | 1 |
| | OOCYTES | | | | | | | | 1 |
| | PHOTORECEPTOR | | | | | 1 | | | 1 |
| | PHOTORECEPTOR ... NEURONS OF ... RETINA | | | | | 1 | | | 1 |
| | PHOTORECEPTOR NEURONS | | | | | 1 | | | 1 |
| | PHOTORECEPTORS | | | | | 1 | | | 1 |
| | PHOTOSENSORY NEURON | | | | | 1 | | | 1 |
| | PHOTOSENSORY NEURONS | | | | | 1 | | | 1 |
| | PHOTOSENSORY NEURONS IN ... RETINA | | | | | 1 | | | 1 |
| | PHOTOSENSORY NEURONS OF ... RETINA | | | | | 1 | | | 1 |
| | PRIMARY ... NEURON | | | | | 1 | | | 1 |
| | PRIMARY ... NEURONS | | | | | 1 | | | 1 |
| | RECEPTORAL ... NEURONS | | | | | 1 | | | 1 |
| | RECEPTORAL NEURONS | | | | | 1 | | | |
| | ROD PHOTORECEPTOR | | | | | 1 | | | 1 |
| | ROD PHOTORECEPTOR NEURONAL | | | | | 1 | | | 1 |
| | ROD PHOTOSENSORY NEURONS | | | | | 1 | | | 1 |
| file59 | APOPTOTIC CELLS | CELL | CELL | 1 | | 1 | 1 | | 1 |
| | CELL | CELLS | MESENCHYMAL CELL | 1 | | | 1 | | |
| | CELLS | MESENCHYMAL CELL | SMOOTH MUSCLE CELL | 1 | | | 1 | | 1 |
| | EPITHELIAL ... CELL | MUSCLE CELL | | | 1 | 1 | | | 1 |
| | MESENCHYMAL CELL | MUSCLE CELLS | | | 1 | | | | 1 |
| | NC ... CELL | SMOOTH MUSCLE CELL | | 1 | | 1 | | | 1 |
| | NCCS | SMOOTH MUSCLE CELLS | | 1 | | 1 | | | 1 |
| | NEURAL CREST CELL | STEM CELLS | | 1 | | 1 | | | 1 |
| | NEURAL CREST CELLS | | | | | 1 | | | 1 |
| | SMOOTH MUSCLE CELL | | | | | | | | |
| | SMOOTH MUSCLE CELLS | | | | | | | | 1 |
| | STEM CELLS | | | | | | | | 1 |

| | | | | | | | | | |
|---|---|---|---|---|---|---|---|---|---|
| file60 | CELL | CELL | CELL | 1 | | | 1 | | |
| | CELLS | CELLS | EMBRYONIC CELL | 1 | | | | 1 | 1 |
| | CELLULAR | EMBRYONIC CELL | EMBRYONIC FIBROBLAST | | 1 | 1 | | 1 | 1 |
| | FIBROBLAST | EMBRYONIC CELLS | EMBRYONIC STEM CELL | | 1 | | | 1 | 1 |
| | FIBROBLASTS | EMBRYONIC FIBROBLASTS | STEM CELL | | 1 | | 1 | | 1 |
| | PROMYELOCYTIC | EMBRYONIC STEM CELL | | | 1 | 1 | | | 1 |
| | STEM CELL | EMBRYONIC STEM CELLS | | | 1 | | | | |
| | STEM CELLS | FIBROBLAST | | 1 | | | | | 1 |
| | | FIBROBLASTS | | 1 | | | | | |
| | | STEM CELL | | 1 | | | | | |
| | | STEM CELLS | | 1 | | | | | |
| file61 | APOPTOTIC CELLS | CELL | CELL | 1 | | 1 | 1 | | 1 |
| | CELL | CELLS | CHONDROCYTE | 1 | | | 1 | | |
| | CELLS | CHONDROCYTE | FIBROBLAST | 1 | | | 1 | | 1 |
| | CHONDROCYTE | CHONDROCYTES | OSTEOBLAST | 1 | | | 1 | | |
| | CHONDROCYTES | FIBROBLAST | OSTEOCLAST | 1 | | | 1 | | 1 |
| | FIBROBLAST | HYPERTROPHIC CHONDROCYTES | | 1 | | | | | |
| | FIBROBLASTIC | OSTEOBLAST | | 1 | | 1 | | | 1 |
| | FIBROBLASTIC CELLS | OSTEOBLASTS | | 1 | | 1 | | | 1 |
| | HYPERTROPHIC CHONDROCYTES | OSTEOCLAST | | 1 | | | | | 1 |
| | OSTEOBLAST | OSTEOCLASTS | | 1 | | | | | |
| | OSTEOBLAST CELLS | OSTEOPROGENITOR CELLS | | 1 | | 1 | | | 1 |
| | OSTEOBLASTIC | | | | | 1 | | | 1 |
| | OSTEOBLASTS | | | | | | | | 1 |
| | OSTEOCLAST | | | | | | | | |
| | OSTEOCLASTS | | | | | | | | 1 |
| | OSTEOPROGENITOR CELLS | | | | | | | | 1 |
| | OSTEOPROGENITORS | | | | | 1 | | | 1 |
| file62 | BLOOD CELLS | BLOOD CELLS | CELL | 1 | | | 1 | | 1 |
| | CELL | CELL | LYMPHOCYTE | 1 | | | 1 | | |
| | CELLS | CELLS | REGULATORY T CELL | 1 | | | 1 | | 1 |
| | LYMPHOCYTE | LYMPHOCYTE | T CELL | 1 | | | 1 | | |
| | LYMPHOCYTES | LYMPHOCYTES | | 1 | | | | | 1 |
| | REGULATORY T CELL | REGULATORY T CELL | | 1 | | | | | |
| | T CELL | T CELL | | 1 | | | | | |
| file63 | CELL | CELL | CELL | 1 | | | 1 | | |
| | CELLS | CELLS | GLIAL CELL | 1 | | | 1 | | 1 |
| | EPITHELIAL CELLS | GLIAL CELL | STROMAL CELL | 1 | | 1 | 1 | | 1 |
| | GLIAL CELL | STROMAL CELL | | 1 | | | | | |
| | STROMAL CELL | STROMAL CELLS | | 1 | | | | | |
| | STROMAL CELLS | | | | | | | | 1 |
| file64 | APOPTOTIC CELLS | CELL | CELL | 1 | | 1 | 1 | | 1 |
| | CELL | CELLS | GERM CELL | 1 | | | 1 | | |
| | CELLS | GAMETES | MALE GERM CELL | 1 | | | 1 | | 1 |
| | CELLULAR | GERM CELL | OOCYTE | 1 | | 1 | 1 | | 1 |
| | GAMETES | GERM CELLS | PERITUBULAR MYOID CELL | 1 | | | | 1 | 1 |
| | GERM CELL | GONOCYTES | SERTOLI CELL | 1 | | | 1 | | |
| | GERM CELLS | MALE GERM CELL | SOMATIC CELL | 1 | | | | 1 | 1 |
| | GERM-CELL | MALE GERM CELLS | SPERM | 1 | | 1 | 1 | | 1 |
| | GONOCYTES | MULTINUCLEATE CELLS | SPERMATID | 1 | | 1 | 1 | | 1 |
| | LEYDIG CELLS | OOCYTE | SPERMATOCYTE | 1 | | 1 | 1 | | 1 |
| | MALE GERM CELL | OOCYTES | ZYGOTE | 1 | | | 1 | | |

| | | | | | | | | | |
|---|---|---|---|---|---|---|---|---|---|
| | MALE GERM CELLS | SERTOLI CELL | | 1 | | | | | 1 |
| | MULTINUCLEATE CELLS | SERTOLI CELLS | | 1 | | | | | 1 |
| | MULTINUCLEATED CELLS | SOMATIC CELL | | 1 | | 1 | | | 1 |
| | OOCYTE | SOMATIC CELLS | | 1 | | | | | |
| | OOCYTES | SPERM | | 1 | | | | | 1 |
| | PRIMARY SPERMATOCYTES | SPERMATID | | 1 | | | | | 1 |
| | SERTOLI | SPERMATIDS | | 1 | | 1 | | | 1 |
| | SERTOLI CELL | SPERMATOCYTE | | 1 | | | | | |
| | SERTOLI CELLS | SPERMATOCYTES | | 1 | | | | | 1 |
| | SPERM | ZYGOTE | | 1 | | | | | |
| | SPERMATID | | | | | | | | |
| | SPERMATIDS | | | | | | | | 1 |
| | SPERMATOCYTE | | | | | | | | |
| | SPERMATOCYTES | | | | | | | | 1 |
| | SPERMATOGONIA | | | | | 1 | | | 1 |
| | SPERMATOZOA | | | | | 1 | | | 1 |
| | STEM ... CELLS | | | | | 1 | | | 1 |
| | ZYGOTE | | | | | | | | |
| file65 | LYMPHOCYTES | LYMPHOCYTES | | 1 | | | | | 1 |
| | CELLS | CELLS | | 1 | | | | | 1 |
| | PURKINJE CELLS | PURKINJE CELLS | | 1 | | | | | 1 |
| file66 | ADIPOCYTES | ADIPOCYTES | AMACRINE CELL | 1 | | | 1 | | 1 |
| | AMACRINE | AMACRINE CELL | CELL | 1 | | 1 | 1 | | 1 |
| | AMACRINE ... CELLS | AMACRINE CELLS | FIBROBLAST | 1 | | | 1 | | 1 |
| | AMACRINE CELL | AMACRINE NEURONS | NEURON | 1 | | | 1 | | |
| | AMACRINE CELLS | BIPOLAR NEURONS | RETINAL PROGENITOR CELL | 1 | | | | 1 | |
| | AMACRINE NEURONS | CELL | | 1 | | | | | 1 |
| | APOPTOTIC CELLS | CELLS | | 1 | | 1 | | | 1 |
| | BIPOLAR | CHOLINERGIC NEURONS | | 1 | | 1 | | | 1 |
| | BIPOLAR CELL | ENTEROCYTES | | 1 | | 1 | | | 1 |
| | BIPOLAR CELLS | FIBROBLAST | | 1 | | 1 | | | 1 |
| | BIPOLAR CELLS IN ... RETINA | FIBROBLASTS | | 1 | | 1 | | | 1 |
| | BIPOLAR NEURONS | FOREBRAIN NEURONS | | | 1 | | | | 1 |
| | BIPOLARS | INTERNEURONS | | 1 | | 1 | | | 1 |
| | CELL | MüLLER CELLS | | 1 | | | | | |
| | CELLS | NEURON | | 1 | | | | | 1 |
| | CHOLINERGIC NEURONS | NEURONS | | 1 | | | | | 1 |
| | CNS NEURONS | NULL CELLS | | | 1 | | | | 1 |
| | CONE | RETINAL CELL | | | 1 | 1 | | | 1 |
| | CONE BIPOLAR CELLS | RETINAL CELLS | | | 1 | 1 | | | 1 |
| | CONES | RETINAL PROGENITOR CELL | | | 1 | 1 | | | 1 |
| | ENTEROCYTES | ROD BIPOLAR CELLS | | 1 | | | | | 1 |
| | FIBROBLAST | | | | | | | | |
| | FIBROBLASTS | | | | | | | | 1 |
| | GANGION CELLS | | | | | 1 | | | 1 |
| | GANGLION | | | | | 1 | | | 1 |
| | GANGLION ... CELL | | | | | 1 | | | 1 |
| | GANGLION ... CELLS | | | | | 1 | | | 1 |
| | GANGLION ... CELLS IN ... RETINA | | | | | 1 | | | 1 |
| | GANGLION CELL | | | | | 1 | | | 1 |
| | GANGLION CELLS | | | | | 1 | | | 1 |
| | GANGLION NEURONS | | | | | 1 | | | 1 |
| | GLIAL | | | | | 1 | | | 1 |
| | HORIZONTAL ... CELLS | | | | | 1 | | | 1 |
| | HORIZONTAL ... NEURONS | | | | | 1 | | | 1 |

| | | | | | | | | | |
|---|---|---|---|---|---|---|---|---|---|
| | INTERNEURONS | | | | | | | | 1 |
| | MÃ¼LLER CELLS | | | | | 1 | | | 1 |
| | MÃ¼LLER GLIA CELL | | | | | 1 | | | 1 |
| | NEURON | | | | | | | | |
| | NEURONAL | | | | | 1 | | | 1 |
| | NEURONS | | | | | | | | 1 |
| | ON BIPOLAR CELLS | | | | | 1 | | | 1 |
| | PHOTORECEPTOR | | | | | 1 | | | 1 |
| | PHOTORECEPTORS | | | | | 1 | | | 1 |
| | RETINAL ... GANGLION ... CELLS | | | | | 1 | | | 1 |
| | RETINAL ... ROD CELLS | | | | | 1 | | | 1 |
| | RETINAL BIPOLAR ... CELLS | | | | | 1 | | | 1 |
| | ROD ... BIPOLAR CELLS | | | | | | | | 1 |
| | ROD ... CELL | | | | | 1 | | | 1 |
| | ROD ... CELLS | | | | | 1 | | | 1 |
| | ROD ... CELLS IN ... RETINA | | | | | 1 | | | 1 |
| | ROD BIPOLAR ... CELLS | | | | | | | | |
| | ROD BIPOLAR CELLS | | | | | | | | 1 |
| | ROD PHOTORECEPTORS | | | | | 1 | | | 1 |
| | RODS | | | | | 1 | | | 1 |
| file67 | CELL | ARCS | CELL | | 1 | | 1 | | |
| | CELLS | CELL | EMBRYONIC STEM CELL | 1 | | | 1 | | 1 |
| | EGGS | CELLS | GAMETE | 1 | | 1 | 1 | | 1 |
| | GAMETE | EMBRYONIC STEM CELL | GERM CELL | | 1 | 1 | 1 | | |
| | GAMETES | GAMETES | OOCYTE | 1 | | | 1 | | 1 |
| | GERM CELL | GERM CELL | SPERM | 1 | | | 1 | | |
| | OOCYTE | OOCYTE | SPERMATOCYTE | 1 | | | 1 | | |
| | OOCYTES | OOCYTES | STEM CELL | 1 | | | 1 | | 1 |
| | SPERM | SPERM | | 1 | | | | | |
| | SPERMATIDS | SPERMATIDS | | 1 | | | | | 1 |
| | SPERMATOCYTE | SPERMATOCYTE | | 1 | | | | | |
| | SPERMATOCYTES | SPERMATOCYTES | | 1 | | | | | 1 |
| | SPERMATOGONIA | STEM CELL | | 1 | | 1 | | | 1 |
| | SPORE | | | | | 1 | | | 1 |
| | STEM CELL | | | | | | | | |
| | STEM CELLS | | | | | 1 | | | 1 |
| | | | | 256 | 70 | 175 | 98 | 22 | 333 |
| | | | Accuracy | 0,79 | | | 0,82 | | |
| | | | Recall | 0,59 | | | 0,23 | | |
| | | | F-Measure | 0,68 | | | 0,36 | | |
