## Supplementary_Excel_file_S3 2015 Human Disease derived for "OntoContext, a new python package for gene contextualization based on the annotation of biomedical texts"

| Num file | Refrence Biotext Annotation | OntoContext Annotation | TP | FP | FN |
|---|---|---|---|---|---|
| file1 | | | | | |
| file2 | | | | | |
| file3 | GESTATIONAL DIABETES | GESTATIONAL DIABETES | 1 | | |
| file4 | AUTOSOMAL TRISOMIES | DOWN SYNDROME | 1 | | 1 |
| | BRADYCARDIA | HYDROCEPHALUS | | 1 | 1 |
| | CYTOGENETIC ABERRATIONS | HYDRONEPHROSIS | 1 | | 1 |
| | DOWN SYNDROME | MICROCEPHALY | 1 | | |
| | DUODENAL ATRESIA | PATAU SYNDROME | 1 | | 1 |
| | EDWARD SYNDROME | SPINA BIFIDA | 1 | | 1 |
| | HYDRONEPHROSIS | SYNDROME | | 1 | |
| | MICROCEPHALY | TRISOMY 13 | 1 | | |
| | NONIMMUNE HYDROPS FETALIS | TRISOMY 18 | 1 | | 1 |
| | PATAU SYNDROME | | | | |
| | PYELECTASIS | | | | 1 |
| | SPINA BIFIDA | | | | |
| | TRISOMIES | | | | 1 |
| | TRISOMY 13 | | | | |
| | TRISOMY 18 | | | | |
| | TRISOMY 21 | | | | 1 |
| file5 | PRE-ECLAMPSIA | PRE-ECLAMPSIA | 1 | | |
| | PREECLAMPSIA | PREECLAMPSIA | 1 | | |
| file6 | | | | | |
| file7 | HEMIPLEGIA | HEMIPLEGIA | 1 | | |
| | ASCITES | | | 1 | |
| | MASSIVE FETOMATERNAL HEMORRHAGE | | | 1 | |
| | HYPOTONIA | | | 1 | |
| | HEPATOSPLENOMEGALY | | | 1 | |
| file8 | OMPHALOCELE | OMPHALOCELE | 1 | | |
| | EXSTROPHY OF THE BLADDER | IMPERFORATE ANUS | | 1 | |
| | MENINGOMYELOCELES | | | | 1 |
| | OMPHALOCELE-EXSTROPHY-IMPERFORATE ANUS-SPINAL DEFECTS (OEIS COMPLEX) | | | | 1 |
| | OEIS COMPLEX | | | | 1 |
| | SPINAL DEFECTS | | | | 1 |
| file9 | NONIMMUNE HYDROPS FETALIS | PERICARDIAL EFFUSION | 1 | | 1 |
| | HEMATOMA | | | | 1 |

| | |
|---|---|
| Accuracy | 0,55 |
| Recall | 0,30 |
| F-Measure | 0,39 |

| | | | | | |
|---|---|---|---|---|---|
| | ASCITES | | | | 1 |
| | LUCENCIES | | | | 1 |
| | HYDROTHORAX | | | | 1 |
| | PERICARDIAL EFFUSION | | | | |
| file10 | CHRONIC PROGRESSIVE EXTERNAL OPHTHALMOPLEG | SYNDROME | | 1 | |
| | MITOCHONDRIAL DISEASES | PROGRESSIVE EXTERNAL OPHTHALMOPLEGIA | | 1 | |
| | HYPERTENSION | CHRONIC PROGRESSIVE EXTERNAL OPHTHALMOPLEGIA | 1 | | |
| | | MITOCHONDRIAL DISEASES | 1 | | |
| | | HYPERTENSION | 1 | | |
| file11 | ALZHEIMER DISEASE (AD) | ALZHEIMER DISEASE | 1 | | |
| | | NEUROLOGICAL DISORDERS | | 1 | |
| file12 | OVARIAN CANCER | OVARIAN CANCER | 1 | | |
| | BREAST CANCER | BREAST CANCER | 1 | | |
| | CANCER | CANCER | 1 | | |
| file13 | BREAST CANCER | BREAST CANCER | 1 | | |
| | CANCERS | CANCER | | 1 | |
| | COLON CANCER | CANCERS | 1 | | |
| | PROSTATE CANCER | COLON CANCER | 1 | | |
| | PROSTATE CANCERS | HEREDITARY PROSTATE CANCER | | 1 | |
| | TUMORS | PROSTATE CANCER | 1 | | 1 |
| | | PROSTATE CANCERS | 1 | | |
| file14 | MULTIPLE SCLEROSIS (MS) | MULTIPLE SCLEROSIS | 1 | | |
| file15 | SARCOMAS | SARCOMA | | 1 | |
| | CANCERS | CANCERS | 1 | | |
| | CANCER | PRIMARY CANCERS | | 1 | |
| | RETINOBLASTOMA | CANCER | 1 | | |
| | | SARCOMAS | 1 | | |
| file16 | PREMATURE RUPTURE OF THE MEMBRANES (PROM) | CHORIOAMNIONITIS | | 1 | 1 |
| | PRETERM PROM | | | | 1 |
| file17 | | | | | |
| file18 | PARVOVIRUS INFECTION | PERICARDIAL EFFUSION | 1 | | 1 |
| | PERICARDIAL EFFUSIONS | PERICARDIAL EFFUSIONS | 1 | | |
| | PARVOVIRUS | | | | 1 |

| | | | | | |
|---|---|---|---|---|---|
| | HYDROPS FETALIS | | | | 1 |
| | PARVOVIRUS B19 INFECTION | | | | 1 |
| | PERICARDIAL EFFUSION | | | | |
| file19 | DEAFNESS | | | | 1 |
| file20 | | ALLERGIC | | 1 | |
| file21 | SEIZURES | EPILEPSY | 1 | | 1 |
| | EPILEPSY | ATTENTION DEFICIT HYPERACTIVITY DISORDER | | 1 | |
| | ADHD | ADHD | 1 | | |
| file22 | | | | | |
| file23 | | | | | |
| file24 | | | | | |
| file25 | | | | | |
| file26 | ISCHAEMIC STROKE | | | | 1 |
| | MYOCARDIAL INFARCTION | | | | 1 |
| | CARDIOVASCULAR MORBIDITY AND MORTALITY AFTER MYOCARDIAL INFARCTION | | | | 1 |
| file27 | CARDIAC ISCHAEMIC DEATH | HEART DISEASE | | 1 | 1 |
| | CHLAMYDIA PNEUMONIAE | CORONARY HEART DISEASE | 1 | | 1 |
| | CORONARY HEART DISEASE | | | | |
| | CORONARY-ARTERY DISEASE | | | | 1 |
| | ISCHAEMIA | | | | 1 |
| | MYOCARDIAL INFARCTION | | | | 1 |
| | UNSTABLE ANGINA | | | | 1 |
| file28 | STAGE IB AND IIA CERVICAL CARCINOMA | CARCINOMA | | 1 | 1 |
| | CERVICAL CARCINOMA | | | | 1 |
| file29 | | | | | |
| file30 | | | | | |
| file31 | | | | | |
| file32 | | | | | |
| file33 | | | | | |

| | | | | | |
|---|---|---|---|---|---|
| file34 | | LEECHES | | 1 | |
| file35 | ADENOCARCINOMA | ADENOCARCINOMA | 1 | | |
| | CANCERS | CANCER | | 1 | |
| | COLON CANCER | CANCERS | 1 | | |
| | COLON TUMORS | COLON CANCER | 1 | | 1 |
| | FIBROSIS | | | | 1 |
| | LYMPHATIC INVASION | | | | 1 |
| | NECROSIS | | | | 1 |
| | NUCLEAR ATYPIA | | | | 1 |
| | TUMORS | | | | 1 |
| file36 | GLYCOGEN STORAGE DISEASE TYPE II (GSDII) | GLYCOGEN STORAGE DISEASE TYPE II | 1 | | |
| | | GLYCOGEN STORAGE DISEASE | | 1 | |
| file37 | AUTONOMIC DYSFUNCTION | MULTIPLE SCLEROSIS | 1 | | 1 |
| | CEREBRAL PALSY | PARKINSON'S DISEASE | 1 | | |
| | MULTIPLE SCLEROSIS | CEREBRAL PALSY | 1 | | |
| | NEUROLOGIC DISEASES | | | | 1 |
| | PARKINSON'S DISEASE | | | | |
| file38 | | | | | |
| file39 | MYOFASCIAL TRIGGER POINT PAIN | Group B | | 1 | 1 |
| | MYOFASCIAL TRIGGER POINTS OF UPPER TRAPEZIUS MUSCLE | | | | 1 |
| | MYOFASCIAL TRIGGER POINTS | | | | 1 |
| file40 | OSTEOPOROSIS | OSTEOPOROSIS | 1 | | |
| file41 | C TRACHOMATIS INFECTION | | | | |
| | CHLAMYDIA TRACHOMATIS | | | | |
| | INFECTION | | | | |
| | C TRACHOMATIS | | | | |
| file42 | | SYNDROME | | 1 | |
| file43 | OBESITY | PRIMARY PULMONARY HYPERTENSION | | 1 | |
| | | HYPERTENSION | | 1 | |
| | | NEUROTOXICITY | | 1 | |

| | | | | | |
|---|---|---|---|---|---|
| file44 | ESSENTIAL HYPERTENSION | HYPERTENSION | | 1 | |
| | | ESSENTIAL HYPERTENSION | 1 | | |
| file45 | | | | | |
| file46 | INFECTION WITH ECHINOCOCCUS MULTILOCULARIS | | | | 1 |
| file47 | CHRONIC MYELOID LEUKEMIA (CML) | LEUKEMIA | | 1 | |
| file48 | HUMAN AMNESIC SYNDROME | SYNDROME | | 1 | |
| file49 | | AMNESIA | 1 | | |
| file50 | | | | | |
| file51 | | | | | |
| file52 | | | | | |
| file53 | OSTEOPOROSIS | OSTEOPOROSIS | 1 | | |
| file54 | | | | | |
| file55 | | | | | |
| file56 | TUMOURS | CANCERS | 1 | | 1 |
| | CANCERS | CANCER | 1 | | |
| | COLONIC TUMOURS | | | | 1 |
| | CANCER | | | | |
| file57 | | TELANGIECTASIA | | 1 | |
| | | ATAXIA TELANGIECTASIA | | 1 | |
| file58 | | | | | |
| file59 | SEVERE FOOT INFECTIONS IN DIABETIC PATIENTS | CELLULITIS | | 1 | 1 |
| | FOOT INFECTION IN DIABETIC PATIENTS | | | | 1 |
| | FOOT INFECTIONS | | | | 1 |

| | | | | | |
|---|---|---|---|---|---|
| | DIABETES | | | | 1 |
| file60 | ORONARY-ARTERY THROMB | | | | 1 |
| file61 | CARDIOVASCULAR SYMPTOMS | HEART DISEASE | | 1 | 1 |
| | HEART MURMUR | HYPERTENSION | | 1 | 1 |
| | VALVE DISEASE | | | | 1 |
| | PULMONARY HYPERTENSION | | | | 1 |
| file62 | TRANSFER LESIONS | | | | 1 |
| | HALLUX SUBLUXATION | | | | 1 |
| | HEUMATOID ARTHRITI | | | | 1 |
| file63 | STROKES | | | | 1 |
| | CEREBROVASCULAR ACCIDENT | | | | 1 |
| | CEREBROVASCULAR ACCIDENTS | | | | 1 |
| | STROKE | | | | 1 |
| file64 | HIGH BLOOD PRESSURE | HIGH BLOOD PRESSURE | 1 | | |
| file65 | | | | | |
| file66 | EARLY PARKINSON'S DISEASE | PARKINSON'S DISEASE | 1 | | 1 |
| | PARKINSON'S DISEASE | | | | |
| file67 | | | | | |
| file68 | | osteomalacia | | 1 | |
| file69 | | | | | 1 |
| file70 | | | | | |
| file71 | OTITIS MEDIA | OTITIS MEDIA | 1 | | |
| | PHARYNGITIS | PHARYNGITIS | 1 | | |
| | QUINSY | QUINSY | 1 | | |
| | SINUSITIS | SINUSITIS | 1 | | |
| | SORE THROAT | TONSILLITIS | 1 | | 1 |
| | TONSILLITIS | | 1 | | |
| | UPPER RESPIRATORY CONDITIONS | | | | 1 |
| file72 | POSTOPERATIVE PAIN RELIEF | ANALGESIA | | 1 | 1 |

| | | | | | |
|---|---|---|---|---|---|
| file73 | | | | | |
| file74 | EXTRAMEDULLARY HEMATOPOIESIS | neutropenia | | 1 | 1 |
| | SPLENOMEGALY | | | | 1 |
| | CHRONIC NEUTROPENIA | | | | 1 |
| file75 | | | | | |
| file76 | CROHN'S DISEASE | | | | 1 |
| | ABDOMINAL TUBERCULOSIS | | | | 1 |
| file77 | EMALE STRESS URINARY INCONTINENC | | | | 1 |
| file78 | | | | | |
| file79 | HELICOBACTER PYLORI INFECTION | HELICOBACTER PYLORI INFECTION | 1 | | |
| | CORPAL GASTRITIS | | | | 1 |
| | H. PYLORI INFECTION | | | | 1 |
| file80 | PREECLAMPSIA (PROTEINURIC HYPERTENSION) | HYPERTENSION | | 1 | 1 |
| | PREECLAMPSIA | PREECLAMPSIA | 1 | | |
| file81 | | | | | |
| file82 | | | | | |
| file83 | | | | | |
| file84 | SCHIZOPHRENIC ILLNESSES | | | | 1 |
| | SCHIZOPHRENIC SYMPTOMS | | | | 1 |
| file85 | CLOACAL ANOMALIES | | | | 1 |
| file86 | AMYLOIDOSIS | HEART DISEASE | | 1 | 1 |
| | CARDIAC AMYLOID HEART DISEASE | | | | 1 |
| | CARDIAC AMYLOIDOSIS | | | | 1 |
| | HEART FAILURE | | | | 1 |
| | MYOCARDIAL AMYLOIDOSIS | | | | 1 |
| | PROTEIN DEPOSITION DISEASES | | | | 1 |

| | | | | | |
|---|---|---|---|---|---|
| file87 | DIABETIC RETINOPATHY | | | | 1 |
| | RETINOPATHY | | | | 1 |
| | DIABETES | | | | 1 |
| file88 | GUNSHOT WOUNDS TO THE ABDOMEN | | | | 1 |
| | GUNSHOT WOUNDS | | | | 1 |
| | BLUNT TRAUMA | | | | 1 |
| | BOWEL PERFORATIONS | | | | 1 |
| | STAB WOUNDS | | | | 1 |
| | INTRA-ABDOMINAL HEMORRHAGE | | | | 1 |
| | INTRA-ABDOMINAL INJURY | | | | 1 |
| file89 | ABDOMINAL PAIN | DIARRHOEA | 1 | | 1 |
| | PARASITOSIS | | | | 1 |
| | PARASITATION | | | | 1 |
| | DIARRHOEA | | | | |
| | BLASTOCYSTIS HOMINIS PARASITATION | | | | 1 |
| file90 | | CANCER | | 1 | |
| file91 | | | | | |
| file92 | | | | | |
| file93 | LUNG CANCER | LUNG CANCER | 1 | | |
| | | CANCER | | 1 | |
| file94 | SYNCOPE | | | | 1 |
| | BLEEDING | | | | 1 |
| | MAJOR PULMONARY EMBOLISM | | | | 1 |
| | PRIMARY THROMBOLYSIS | | | | 1 |
| | HEMODYNAMIC COMPROMISE | | | | 1 |
| | CEREBRAL BLEEDING | | | | 1 |
| | CARDIOGENIC SHOCK | | | | 1 |
| | ARTERIAL HYPOTENSION | | | | 1 |
| | CONGESTIVE HEART FAILURE | | | | 1 |
| | CHRONIC PULMONARY DISEASE | | | | 1 |
| file95 | PLEURAL MESOTHELIOMA | ADENOCARCINOMA | 1 | | |
| | COLONIC POLYPS | MALIGNANT MESOTHELIOMA | | 1 | 1 |
| | MESOTHELIOMA | | | | 1 |
| | METASTATIC MALIGNANT MESOTHELIOMA | | | | 1 |
| | MALIGNANT PLEURAL MESOTHELIOMA | | | | 1 |
| | ADENOCARCINOMA | | | | 1 |

| | | | | | |
|---|---|---|---|---|---|
| file96 | PERILESIONAL EDEMA | DYSKINESIAS | | 1 | 1 |
| | VENOUS INFARCTION | APHASIA | | 1 | 1 |
| | EDEMA | PARKINSON'S DISEASE | | 1 | 1 |
| | ADVANCED PARKINSON'S DISEASE | | | | 1 |
| | HEMORRHAGIC COAGULATION NECROSIS | | | | 1 |
| | PARKINSONIAN SYMPTOMS | | | | 1 |
| | ISCHEMIC INFARCTION | | | | 1 |
| | BROCA'S APHASIA | | | | 1 |
| | ISCHEMIC INFARCTIONS | | | | 1 |
| file97 | | | | | |
| file98 | SENSORINEURAL HEARING LOSS | SENSORINEURAL HEARING LOSS | 1 | | |
| | HEARING IMPAIRMENT | | | | 1 |
| | DEVELOPMENTAL PROBLEMS | | | | 1 |
| | HEARING LOSS | | | | 1 |
| file99 | | | | | |
| file100 | GENODERMATOSIS | GENODERMATOSIS | 1 | | |
| | DARIER DISEASE | | | | 1 |
| file102 | HIGH FEVER | SYNDROME | | 1 | 1 |
| | MACULAR ERYTHRODERMA | RASH | 1 | | 1 |
| | TOXIC SHOCK SYNDROME (TSS) | TOXIC SHOCK | | 1 | |
| | SHOCK | TOXIC SHOCK SYNDROME | 1 | | 1 |
| | HEMORRHAGIC SHOCK | | | | 1 |
| | RASH | | | | |
| | INFLAMMATION | | | | 1 |
| | DESQUAMATION | | | | 1 |
| | ENCEPHALOPATHY | | | | 1 |
| | ERYTHRODERMA | | | | 1 |
| file103 | TERATOMA SYNDROME | TERATOMA | | 1 | 1 |
| | PRIMARY OR METASTATIC MEDIASTINAL GERM CELL | SYNDROME | | 1 | 1 |
| | METASTATIC GERM CELL TUMOR | TERATOMAS | | 1 | 1 |
| | PRIMARY GERM CELL TUMOR | | | | 1 |
| | GERM CELL TUMORS | | | | 1 |
| file104 | | | | | |
| file105 | REVASCULARIZATIONS | | | | 1 |
| | INFARCTIONS | | | | 1 |

| | | | | | |
|---|---|---|---|---|---|
| | CORONARY EVENTS | | | | 1 |
| | REVASCULARIZATION | | | | 1 |
| | INFARCTION | | | | 1 |
| | MYOCARDIAL INFARCTION | | | | 1 |
| file106 | | | | | |
| file107 | ON-OBSTRUCTIVE AZOOSPERMI | AZOOSPERMIA | | 1 | 1 |
| file108 | TUMORS | MANTLE CELL LYMPHOMAS | 1 | | 1 |
| | B CELL LYMPHOMAS | LYMPHOMAS | | 1 | 1 |
| | MALIGNANT LYMPHOMAS | MANTLE CELL LYMPHOMA | 1 | | 1 |
| | MANTLE CELL LYMPHOMAS (MCL) | LYMPHOMA | | 1 | |
| | HUMAN LYMPHOID NEOPLASIA | | | | 1 |
| | MANTLE CELL LYMPHOMA | | | | |
| file109 | | | | | |
| file110 | | | | | |
| file111 | PREECLAMPSIA | PREECLAMPSIA | 1 | | |
| file112 | | | | | |
| file113 | | | | | |
| file114 | TYPHOID | TYPHOID | 1 | | |
| | MALARIA | MALARIA | 1 | | |
| | LEPROSY | LEPROSY | 1 | | |
| | TUBERCULOSIS | | | | 1 |
| file115 | | | | | |
| file116 | | | | | |
| file117 | | | | | |
| file118 | | | | | |
| file119 | URINARY TRACT INFECTIONS (UTI) | URINARY TRACT INFECTIONS | 1 | | |
| | UTIS | | | | 1 |
| file120 | | | | | |

| | | | | | |
|---|---|---|---|---|---|
| file121 | ACUTE LYMPHOBLASTIC LEUKAEMIAS | ACUTE LYMPHOBLASTIC LEUKAEMIAS | 1 | | |
| file122 | | | | | |
| file123 | | | | | |
| file124 | | | | | |
| file125 | | | | | |
| file126 | | | | | |
| file127 | PENDRED SYNDROME | GOITRE | | 1 | 1 |
| | CONGENITAL DEAFNESS | SYNDROME | | 1 | 1 |
| | THYROID GOITRE | | | | 1 |
| | HEREDITARY DEAFNESS | | | | 1 |
| file128 | MYELOID LEUKEMIA | MURINE | | 1 | 1 |
| | EMBRYONAL CARCINOMA | LEUKEMIA | | 1 | |
| | | CARCINOMA | | 1 | |
| | | EMBRYONAL CARCINOMA | 1 | | |
| file129 | | | | | |
| file130 | VIVERRID-TYPE RABIES | RABIES | | 1 | 1 |
| | VIVERRID RABIES | | | | 1 |
| file131 | | | | | |
| file132 | ENDOMETRIOSIS | TESTICULAR CANCER | 1 | | |
| | DYSFUNCTION OF THE SEXUAL AND THYROID HORMC | CANCER | | 1 | 1 |
| | HORMONAL DYSFUNCTION | HYPOTHYROIDISM | 1 | | 1 |
| | TESTICULAR CANCER | ENDOMETRIOSIS | 1 | | |
| | HYPOTHYROIDISM | | | | |
| file133 | | | | | |
| file134 | DIABETES MELLITUS | | | | 1 |
| | DYSIPIDEMIAS | | | | 1 |
| | DIABETES 2 | | | | 1 |
| | HYPERLIPIDEMIAS | | | | 1 |
| | DIABETES | | | | 1 |
| file135 | CARDIOVASCULAR DISEASES | CANCER | | 1 | 1 |

| | | | | | |
|---|---|---|---|---|---|
| file136 | ACUTE MIGRAINE | | | | 1 |
| file137 | | SYNDROME | | 1 | |
| file138 | | | | | |
| file139 | | | | | |
| file140 | DIABETES | | | | 1 |
| file141 | CLOT | | | | 1 |
| | DEEP VEIN THROMBOSIS (DVT) | | | | 1 |
| Total | | | 76 | 63 | 178 |
| | | Accuracy | 0,55 | | |
| | | Recall | 0,30 | | |
| | | F-Measure | 0,39 | | |
