## Supplementary_Excel_file_S3 BioText derived for "OntoContext, a new python package for gene contextualization based on the annotation of biomedical texts"

| Num file | Reference BioText Annotation | Annotation OntoContext | TP | FP | FN |
|---|---|---|---|---|---|
| file1 | | | | | |
| file2 | | | | | |
| file3 | GESTATIONAL DIABETES | GESTATIONAL DIABETES | 1 | | |
| | | DIABETES | | | |
| file4 | EDWARD SYNDROME | EDWARD SYNDROME | 1 | | |
| | SPINA BIFIDA | SPINA BIFIDA | 1 | | |
| | DUODENAL ATRESIA | TRISOMY 18 | 1 | | |
| | HYDRONEPHROSIS | PATAU SYNDROME | 1 | | |
| | TRISOMY 18 | AUTOSOMAL TRISOMIES | 1 | | |
| | PATAU SYNDROME | TRISOMY 21 | 1 | | |
| | PYELECTASIS | NONIMMUNE HYDROPS FETALIS | 1 | | |
| | AUTOSOMAL TRISOMIES | TRISOMY 13 | 1 | | |
| | TRISOMY 21 | DUODENAL ATRESIA | 1 | | |
| | NONIMMUNE HYDROPS FETALIS | DOWN SYNDROME | 1 | | |
| | TRISOMY 13 | MICROCEPHALY | 1 | | |
| | MICROCEPHALY | HYDRONEPHROSIS | 1 | | |
| | CYTOGENETIC ABERRATIONS | PYELECTASIS | 1 | | 1 |
| | DOWN SYNDROME | TRISOMIES | 1 | | |
| | TRISOMIES | BRADYCARDIA | 1 | | |
| | BRADYCARDIA | | | | |
| file5 | PRE-ECLAMPSIA | PRE-ECLAMPSIA | 1 | | |
| | PREECLAMPSIA | PREECLAMPSIA | 1 | | |
| file6 | | | | | |
| file7 | HEMIPLEGIA | HEMIPLEGIA | 1 | | |
| | ASCITES | HEPATOSPLENOMEGALY | 1 | | |
| | MASSIVE FETOMATERNAL HEMORRHAGE | MASSIVE FETOMATERNAL HEMORRHAGE | 1 | | |
| | HYPOTONIA | HYPOTONIA | 1 | | |
| | HEPATOSPLENOMEGALY | ASCITES | 1 | | |
| file8 | OMPHALOCELE | OMPHALOCELE | 1 | | |
| | EXSTROPHY OF THE BLADDER | EXSTROPHY OF THE BLADDER | 1 | | |
| | MENINGOMYELOCELES | MENINGOMYELOCELES | 1 | | |
| | OMPHALOCELE-EXSTROPHY-IMPERFORATE ANUS-SPI | OMPHALOCELE-EXSTROPHY-IMPERFORATE ANUS-SPINAL DEF | 1 | | |
| | OEIS COMPLEX | OEIS COMPLEX | 1 | | |
| | SPINAL DEFECTS | SPINAL DEFECTS | 1 | | |
| file9 | NONIMMUNE HYDROPS FETALIS | NONIMMUNE HYDROPS FETALIS | 1 | | |

| | |
|---|---|
| Accuracy | 0,88 |
| Recall | 0,93 |
| F-Measure | 0,91 |

| | | | | | |
|---|---|---|---|---|---|
| | HEMATOMA | HEMATOMA | 1 | | |
| | ASCITES | ASCITES | 1 | | |
| | LUCENCIES | LUCENCIES | 1 | | |
| | HYDROTHORAX | HYDROTHORAX | 1 | | |
| | PERICARDIAL EFFUSION | PERICARDIAL EFFUSION | 1 | | |
| file10 | CHRONIC PROGRESSIVE EXTERNAL OPHTHALMOPLEG | HYPERTENSION | 1 | | |
| | MITOCHONDRIAL DISEASES | CHRONIC PROGRESSIVE EXTERNAL OPHTHALMOPLEGIA (KEAR | 1 | | |
| | HYPERTENSION | MITOCHONDRIAL DISEASES | 1 | | |
| file11 | ALZHEIMER DISEASE (AD) | ALZHEIMER DISEASE (AD) | 1 | | |
| file12 | OVARIAN CANCER | OVARIAN CANCER | 1 | | |
| | BREAST CANCER | BREAST CANCER | 1 | | |
| | CANCER | CANCER | 1 | | |
| file13 | PROSTATE CANCER | PROSTATE CANCER | 1 | | |
| | PROSTATE CANCERS | CANCER | 1 | 1 | |
| | TUMORS | PROSTATE CANCERS | 1 | | |
| | COLON CANCER | TUMORS | 1 | | |
| | CANCERS | COLON CANCER | 1 | | |
| | BREAST CANCER | CANCERS | 1 | | |
| | | BREAST CANCER | | | |
| file14 | MULTIPLE SCLEROSIS (MS) | MULTIPLE SCLEROSIS | 1 | | |
| file15 | SARCOMAS | SARCOMAS | 1 | | |
| | CANCERS | CANCERS | 1 | | |
| | CANCER | RETINOBLASTOMA | 1 | | |
| | RETINOBLASTOMA | CANCER | 1 | | |
| file16 | PREMATURE RUPTURE OF THE MEMBRANES (PROM) | PREMATURE RUPTURE OF THE MEMBRANES (PROM) | 1 | | |
| | PRETERM PROM | PRETERM PROM | 1 | | |
| file17 | | | | | |
| file18 | PARVOVIRUS INFECTION | PARVOVIRUS INFECTION | 1 | | |
| | PERICARDIAL EFFUSIONS | PERICARDIAL EFFUSIONS | 1 | | |
| | PARVOVIRUS | PARVOVIRUS | 1 | | |
| | HYDROPS FETALIS | INFECTION | 1 | | |
| | PARVOVIRUS B19 INFECTION | PARVOVIRUS B19 INFECTION | 1 | | |
| | PERICARDIAL EFFUSION | PERICARDIAL EFFUSION | 1 | | |
| file19 | DEAFNESS | DEAFNESS | 1 | | 1 |

| | | | | | |
|---|---|---|---|---|---|
| file20 | | | | | |
| file21 | SEIZURES | SEIZURES | 1 | | |
| | EPILEPSY | EPILEPSY | 1 | | |
| | ADHD | ADHD | 1 | | |
| file22 | | | | | |
| file23 | | | | | |
| file24 | | | | | |
| file25 | | | | | |
| file26 | ISCHAEMIC STROKE | BLEEDING | | 1 | |
| | MYOCARDIAL INFARCTION | CARDIOVASCULAR MORBIDITY AND MORTALITY AFTER MYOC | 1 | | |
| | CARDIOVASCULAR MORBIDITY AND MORTALITY AFTE | ISCHAEMIC STROKE | 1 | | |
| | | STROKE | | 1 | |
| | | INFARCTION | | 1 | |
| | | MYOCARDIAL INFARCTION | 1 | | |
| file27 | CHLAMYDIA PNEUMONIAE | CHLAMYDIA PNEUMONIAE | 1 | | |
| | UNSTABLE ANGINA | UNSTABLE ANGINA | 1 | | |
| | MYOCARDIAL INFARCTION | INFARCTION | | 1 | |
| | CORONARY-ARTERY DISEASE | MYOCARDIAL INFARCTION | 1 | | |
| | CORONARY HEART DISEASE | CORONARY-ARTERY DISEASE | 1 | | |
| | CARDIAC ISCHAEMIC DEATH | CORONARY HEART DISEASE | 1 | | |
| | ISCHAEMIA | CARDIAC ISCHAEMIC DEATH | 1 | | |
| | | ISCHAEMIA | 1 | | |
| file28 | STAGE IB AND IIA CERVICAL CARCINOMA | CERVICAL CARCINOMA | 1 | | |
| | CERVICAL CARCINOMA | | | | 1 |
| file29 | | | | | |
| file30 | | | | | |
| file31 | | | | | |
| file32 | | | | | |
| file33 | | | | | |

| | | | | |
|---|---|---|---|---|
| file34 | | | | |
| file35 | COLON TUMORS | CANCER | | 1 |
| | LYMPHATIC INVASION | COLON TUMORS | 1 | |
| | FIBROSIS | LYMPHATIC INVASION | 1 | |
| | TUMORS | TUMORS | 1 | |
| | NUCLEAR ATYPIA | NECROSIS | 1 | |
| | COLON CANCER | NUCLEAR ATYPIA | 1 | |
| | CANCERS | COLON CANCER | 1 | |
| | ADENOCARCINOMA | ADENOCARCINOMA | 1 | |
| | NECROSIS | CANCERS | 1 | |
| | | FIBROSIS | 1 | |
| file36 | GLYCOGEN STORAGE DISEASE TYPE II (GSDII) | GLYCOGEN STORAGE DISEASE TYPE II (GSDII) | 1 | |
| file37 | AUTONOMIC DYSFUNCTION | AUTONOMIC DYSFUNCTION | 1 | |
| | MULTIPLE SCLEROSIS | MULTIPLE SCLEROSIS | 1 | |
| | PARKINSON'S DISEASE | PARKINSON'S DISEASE | 1 | |
| | NEUROLOGIC DISEASES | NEUROLOGIC DISEASES | 1 | |
| | CEREBRAL PALSY | CEREBRAL PALSY | 1 | |
| file38 | | | | |
| file39 | MYOFASCIAL TRIGGER POINT PAIN | MYOFASCIAL TRIGGER POINT PAIN | 1 | |
| | MYOFASCIAL TRIGGER POINTS OF UPPER TRAPEZIUS I | MYOFASCIAL TRIGGER POINTS OF UPPER TRAPEZIUS MUSCLE | 1 | |
| | MYOFASCIAL TRIGGER POINTS | MYOFASCIAL TRIGGER POINTS | 1 | |
| file40 | OSTEOPOROSIS | OSTEOPOROSIS | 1 | |
| file41 | C TRACHOMATIS INFECTION | C TRACHOMATIS INFECTION | 1 | |
| | CHLAMYDIA TRACHOMATIS | CHLAMYDIA TRACHOMATIS | 1 | |
| | INFECTION | INFECTION | 1 | |
| | C TRACHOMATIS | C TRACHOMATIS | 1 | |
| file42 | | | | |
| file43 | OBESITY | OBESITY | 1 | |
| | | HYPERTENSION | | 1 |
| | | PULMONARY HYPERTENSION | | 1 |
| file44 | ESSENTIAL HYPERTENSION | ESSENTIAL HYPERTENSION | 1 | |
| | | HYPERTENSION | | 1 |
| file45 | | | | |

| | | | | | |
|---|---|---|---|---|---|
| file46 | INFECTION WITH ECHINOCOCCUS MULTILOCULARIS | INFECTION WITH ECHINOCOCCUS MULTILOCULARIS | 1 | | |
| | | INFECTION | | 1 | |
| file47 | CHRONIC MYELOID LEUKEMIA (CML) | MYELOID LEUKEMIA | | 1 | |
| | | CHRONIC MYELOID LEUKEMIA (CML) | 1 | | |
| file48 | HUMAN AMNESIC SYNDROME | HUMAN AMNESIC SYNDROME | 1 | | |
| file49 | | | | | |
| file50 | | | | | |
| file51 | | | | | |
| file52 | | | | | |
| file53 | OSTEOPOROSIS | OSTEOPOROSIS | 1 | | |
| file54 | | | | | |
| file55 | | | | | |
| file56 | TUMOURS | TUMOURS | 1 | | |
| | CANCERS | CANCERS | 1 | | |
| | COLONIC TUMOURS | COLONIC TUMOURS | 1 | | |
| | CANCER | CANCER | 1 | | |
| file57 | | | | | |
| file58 | | | | | |
| file59 | SEVERE FOOT INFECTIONS IN DIABETIC PATIENTS | FOOT INFECTIONS | 1 | | 1 |
| | FOOT INFECTION IN DIABETIC PATIENTS | INFECTION | | 1 | 1 |
| | FOOT INFECTIONS | DIABETES | 1 | | |
| | DIABETES | | | | |
| file60 | ORONARY-ARTERY THROMB | MYOCARDIAL INFARCTION | | 1 | 1 |
| | | STROKE | | 1 | |
| | | INFARCTION | | | |
| file61 | CARDIOVASCULAR SYMPTOMS | HEART MURMUR | 1 | | |
| | HEART MURMUR | CARDIOVASCULAR SYMPTOMS | 1 | | |
| | VALVE DISEASE | HYPERTENSION | | 1 | |

| | | | | | |
|---|---|---|---|---|---|
| | PULMONARY HYPERTENSION | VALVE DISEASE | 1 | | |
| | | PULMONARY HYPERTENSION | 1 | | |
| file62 | TRANSFER LESIONS | TRANSFER LESIONS | 1 | | |
| | HALLUX SUBLUXATION | HALLUX SUBLUXATION | 1 | | |
| | HEUMATOID ARTHRITI | | | | 1 |
| file63 | STROKES | STROKES | 1 | | |
| | CEREBROVASCULAR ACCIDENT | CEREBROVASCULAR ACCIDENT | 1 | | |
| | CEREBROVASCULAR ACCIDENTS | CEREBROVASCULAR ACCIDENTS | 1 | | |
| | STROKE | STROKE | 1 | | |
| file64 | HIGH BLOOD PRESSURE | HIGH BLOOD PRESSURE | 1 | | |
| file65 | | | | | |
| file66 | EARLY PARKINSON'S DISEASE | EARLY PARKINSON'S DISEASE | 1 | | |
| | PARKINSON'S DISEASE | PARKINSON'S DISEASE | 1 | | |
| file67 | | | | | |
| file68 | | | | | |
| file69 | | | | | |
| file70 | | | | | |
| file71 | UPPER RESPIRATORY CONDITIONS | OTITIS MEDIA | 1 | | 1 |
| | OTITIS MEDIA | PHARYNGITIS | 1 | | |
| | PHARYNGITIS | SINUSITIS | 1 | | |
| | SINUSITIS | QUINSY | 1 | | |
| | QUINSY | SORE THROAT | 1 | | |
| | SORE THROAT | TONSILLITIS | 1 | | |
| | TONSILLITIS | | | | |
| file72 | POSTOPERATIVE PAIN RELIEF | POSTOPERATIVE PAIN RELIEF | 1 | | |
| file73 | | | | | |
| file74 | EXTRAMEDULLARY HEMATOPOIESIS | EXTRAMEDULLARY HEMATOPOIESIS | 1 | | |
| | SPLENOMEGALY | SPLENOMEGALY | 1 | | |
| | CHRONIC NEUTROPENIA | INFLAMMATION | | 1 | |
| | | CHRONIC NEUTROPENIA | 1 | | |

| | | | | | |
|---|---|---|---|---|---|
| file75 | | | | | |
| file76 | CROHN'S DISEASE | CROHN'S DISEASE | 1 | | |
| | ABDOMINAL TUBERCULOSIS | TUBERCULOSIS | | 1 | |
| | | ABDOMINAL TUBERCULOSIS | 1 | | |
| file77 | EMALE STRESS URINARY INCONTINENC | | | | 1 |
| file78 | | | | | |
| file79 | HELICOBACTER PYLORI INFECTION | HELICOBACTER PYLORI INFECTION | 1 | | |
| | CORPAL GASTRITIS | CORPAL GASTRITIS | 1 | | |
| | H. PYLORI INFECTION | INFECTION | | 1 | 1 |
| file80 | PREECLAMPSIA (PROTEINURIC HYPERTENSION) | HYPERTENSION | 1 | 1 | 1 |
| | PREECLAMPSIA | PREECLAMPSIA | 1 | | |
| file81 | | | | | |
| file82 | | | | | |
| file83 | | | | | |
| file84 | SCHIZOPHRENIC ILLNESSES | SCHIZOPHRENIC ILLNESSES | 1 | | |
| | SCHIZOPHRENIC SYMPTOMS | SCHIZOPHRENIC SYMPTOMS | 1 | | |
| file85 | CLOACAL ANOMALIES | CLOACAL ANOMALIES | 1 | | |
| file86 | MYOCARDIAL AMYLOIDOSIS | MYOCARDIAL AMYLOIDOSIS | 1 | | |
| | HEART FAILURE | HEART FAILURE | 1 | | |
| | CARDIAC AMYLOIDOSIS | CARDIAC AMYLOIDOSIS | 1 | | |
| | CARDIAC AMYLOID HEART DISEASE | CARDIAC AMYLOID HEART DISEASE | 1 | | |
| | AMYLOIDOSIS | AMYLOIDOSIS | 1 | | |
| | PROTEIN DEPOSITION DISEASES | PROTEIN DEPOSITION DISEASES | 1 | | |
| file87 | DIABETIC RETINOPATHY | RETINOPATHY | 1 | | |
| | RETINOPATHY | DIABETIC RETINOPATHY | 1 | | |
| | DIABETES | DIABETES | 1 | | |
| file88 | GUNSHOT WOUNDS TO THE ABDOMEN | GUNSHOT WOUNDS | 1 | | 1 |
| | GUNSHOT WOUNDS | BLUNT TRAUMA | 1 | | |
| | BLUNT TRAUMA | BOWEL PERFORATIONS | 1 | | |
| | BOWEL PERFORATIONS | STAB WOUNDS | 1 | | |
| | STAB WOUNDS | INTRA-ABDOMINAL HEMORRHAGE | 1 | | |

| | | | | |
|---|---|---|---|---|
| | INTRA-ABDOMINAL HEMORRHAGE | INTRA-ABDOMINAL INJURY | 1 | |
| | INTRA-ABDOMINAL INJURY | | | |
| file89 | ABDOMINAL PAIN | ABDOMINAL PAIN | 1 | |
| | PARASITOSIS | PARASITOSIS | 1 | |
| | PARASITATION | PARASITATION | 1 | |
| | DIARRHOEA | DIARRHOEA | 1 | |
| | BLASTOCYSTIS HOMINIS PARASITATION | BLASTOCYSTIS HOMINIS PARASITATION | 1 | |
| file90 | | CANCER | | 1 |
| file91 | | | | |
| file92 | | | | |
| file93 | LUNG CANCER | LUNG CANCER | 1 | |
| | | CANCER | | 1 |
| file94 | SYNCOPE | SYNCOPE | 1 | |
| | BLEEDING | HEART FAILURE | | 1 |
| | MAJOR PULMONARY EMBOLISM | BLEEDING | 1 | |
| | PRIMARY THROMBOLYSIS | MAJOR PULMONARY EMBOLISM | 1 | |
| | HEMODYNAMIC COMPROMISE | SHOCK | | 1 |
| | CEREBRAL BLEEDING | HEMODYNAMIC COMPROMISE | 1 | |
| | CARDIOGENIC SHOCK | CEREBRAL BLEEDING | 1 | |
| | ARTERIAL HYPOTENSION | PRIMARY THROMBOLYSIS | 1 | |
| | CONGESTIVE HEART FAILURE | CARDIOGENIC SHOCK | 1 | |
| | CHRONIC PULMONARY DISEASE | ARTERIAL HYPOTENSION | 1 | |
| | | CONGESTIVE HEART FAILURE | 1 | |
| | | CHRONIC PULMONARY DISEASE | 1 | |
| file95 | PLEURAL MESOTHELIOMA | PLEURAL MESOTHELIOMA | 1 | |
| | COLONIC POLYPS | COLONIC POLYPS | 1 | |
| | MESOTHELIOMA | MESOTHELIOMA | 1 | |
| | METASTATIC MALIGNANT MESOTHELIOMA | METASTATIC MALIGNANT MESOTHELIOMA | 1 | |
| | MALIGNANT PLEURAL MESOTHELIOMA | MALIGNANT PLEURAL MESOTHELIOMA | 1 | |
| | ADENOCARCINOMA | ADENOCARCINOMA | 1 | |
| file96 | PERILESIONAL EDEMA | ISCHEMIC INFARCTIONS | 1 | |
| | VENOUS INFARCTION | VENOUS INFARCTION | 1 | |
| | EDEMA | EDEMA | 1 | |
| | ADVANCED PARKINSON'S DISEASE | ADVANCED PARKINSON'S DISEASE | 1 | |
| | HEMORRHAGIC COAGULATION NECROSIS | NECROSIS | | 1 |
| | PARKINSONIAN SYMPTOMS | HEMORRHAGIC COAGULATION NECROSIS | 1 | |

| | | | | |
|---|---|---|---|---|
| | ISCHEMIC INFARCTION | INFARCTIONS | | 1 |
| | BROCA'S APHASIA | INFARCTION | | 1 |
| | ISCHEMIC INFARCTIONS | ISCHEMIC INFARCTION | 1 | |
| | | BROCA'S APHASIA | 1 | |
| | | PERILESIONAL EDEMA | | |
| | | PARKINSON'S DISEASE | 1 | |
| file97 | | | | |
| file98 | SENSORINEURAL HEARING LOSS | SENSORINEURAL HEARING LOSS | 1 | |
| | HEARING IMPAIRMENT | HEARING IMPAIRMENT | 1 | |
| | DEVELOPMENTAL PROBLEMS | DEVELOPMENTAL PROBLEMS | 1 | |
| | HEARING LOSS | HEARING LOSS | 1 | |
| file99 | | | | |
| file100 | GENODERMATOSIS | GENODERMATOSIS | 1 | |
| | DARIER DISEASE | DARIER DISEASE | 1 | |
| file101 | | | | |
| file102 | HIGH FEVER | HIGH FEVER | 1 | |
| | MACULAR ERYTHRODERMA | MACULAR ERYTHRODERMA | 1 | |
| | TOXIC SHOCK SYNDROME (TSS) | TOXIC SHOCK SYNDROME (TSS) | 1 | |
| | SHOCK | NECROSIS | | 1 |
| | HEMORRHAGIC SHOCK | SHOCK | 1 | |
| | RASH | HEMORRHAGIC SHOCK | 1 | |
| | INFLAMMATION | RASH | 1 | |
| | DESQUAMATION | INFLAMMATION | 1 | |
| | ENCEPHALOPATHY | DESQUAMATION | 1 | |
| | ERYTHRODERMA | ENCEPHALOPATHY | 1 | |
| | | ERYTHRODERMA | 1 | |
| file103 | TERATOMA SYNDROME | GERM CELL TUMORS | 1 | |
| | PRIMARY OR METASTATIC MEDIASTINAL GERM CELL | TUMORS | | 1 |
| | METASTATIC GERM CELL TUMOR | METASTATIC GERM CELL TUMOR | 1 | |
| | PRIMARY GERM CELL TUMOR | PRIMARY OR METASTATIC MEDIASTINAL GERM CELL TUMORS | 1 | |
| | GERM CELL TUMORS | TERATOMA SYNDROME | 1 | |
| | | PRIMARY GERM CELL TUMOR | 1 | |
| file104 | | | | |
| file105 | REVASCULARIZATIONS | REVASCULARIZATIONS | 1 | |
| | INFARCTIONS | INFARCTIONS | 1 | |

| | | | | | |
|---|---|---|---|---|---|
| | CORONARY EVENTS | CORONARY EVENTS | 1 | | |
| | REVASCULARIZATION | REVASCULARIZATION | 1 | | |
| | INFARCTION | INFARCTION | 1 | | |
| | MYOCARDIAL INFARCTION | MYOCARDIAL INFARCTION | 1 | | |
| file106 | | | | | |
| file107 | ON-OBSTRUCTIVE AZOOSPERMI | BLEEDING | | 1 | 1 |
| file108 | TUMORS | MALIGNANT LYMPHOMAS | 1 | | |
| | B CELL LYMPHOMAS | MANTLE CELL LYMPHOMAS (MCL) | 1 | | 1 |
| | MALIGNANT LYMPHOMAS | MANTLE CELL LYMPHOMA | 1 | | |
| | MANTLE CELL LYMPHOMAS (MCL) | TUMORS | 1 | | |
| | HUMAN LYMPHOID NEOPLASIA | | | | 1 |
| | MANTLE CELL LYMPHOMA | | | | |
| file109 | | | | | |
| file110 | | | | | |
| file111 | PREECLAMPSIA | PREECLAMPSIA | 1 | | |
| file112 | | | | | |
| file113 | | | | | |
| file114 | TYPHOID | TYPHOID | 1 | | |
| | MALARIA | MALARIA | 1 | | |
| | LEPROSY | LEPROSY | 1 | | |
| | TUBERCULOSIS | TUBERCULOSIS | 1 | | |
| file115 | | | | | |
| file116 | | | | | |
| file117 | | | | | |
| file118 | | | | | |
| file119 | URINARY TRACT INFECTIONS (UTI) | URINARY TRACT INFECTIONS (UTI) | 1 | | |
| | UTIS | UTIS | 1 | | |
| file120 | | | | | |

| | | | | | |
|---|---|---|---|---|---|
| file121 | ACUTE LYMPHOBLASTIC LEUKAEMIAS | ACUTE LYMPHOBLASTIC LEUKAEMIAS | 1 | | |
| file122 | | | | | |
| file123 | | | | | |
| file124 | | | | | |
| file125 | | | | | |
| file126 | | | | | |
| file127 | PENDRED SYNDROME | HEREDITARY DEAFNESS | 1 | | |
| | CONGENITAL DEAFNESS | DEAFNESS | | 1 | |
| | THYROID GOITRE | CONGENITAL DEAFNESS | 1 | | |
| | HEREDITARY DEAFNESS | THYROID GOITRE | 1 | | |
| | | PENDRED SYNDROME | 1 | | |
| file128 | MYELOID LEUKEMIA | MYELOID LEUKEMIA | 1 | | |
| | EMBRYONAL CARCINOMA | EMBRYONAL CARCINOMA | 1 | | |
| file129 | | | | | |
| file130 | VIVERRID-TYPE RABIES | VIVERRID RABIES | 1 | | 1 |
| | VIVERRID RABIES | | | | |
| file131 | | | | | |
| file132 | ENDOMETRIOSIS | TESTICULAR CANCER | 1 | | |
| | DYSFUNCTION OF THE SEXUAL AND THYROID HORMC | CANCER | | 1 | |
| | HORMONAL DYSFUNCTION | DYSFUNCTION OF THE SEXUAL AND THYROID HORMONE SYST | 1 | | |
| | TESTICULAR CANCER | HORMONAL DYSFUNCTION | 1 | | |
| | HYPOTHYROIDISM | HYPOTHYROIDISM | 1 | | |
| | | ENDOMETRIOSIS | 1 | | |
| file133 | | | | | |
| file134 | DIABETES MELLITUS | HEART FAILURE | | 1 | 1 |
| | DYSIPIDEMIAS | DYSIPIDEMIAS | 1 | | |
| | DIABETES 2 | HYPERLIPIDEMIAS | 1 | | 1 |
| | HYPERLIPIDEMIAS | DIABETES | 1 | | |
| | DIABETES | | | | |
| file135 | CARDIOVASCULAR DISEASES | CARDIOVASCULAR DISEASES | 1 | | |

| | | | | | |
|---|---|---|---|---|---|
| | | CANCER | | 1 | |
| file136 | ACUTE MIGRAINE | ACUTE MIGRAINE | 1 | | |
| file137 | | | | | |
| file138 | | | | | |
| file139 | | | | | |
| file140 | DIABETES | DIABETES | 1 | | |
| file141 | CLOT | DEEP VEIN THROMBOSIS (DVT) | 1 | | |
| | DEEP VEIN THROMBOSIS (DVT) | CLOT | 1 | | |
| Total | | | 247 | 33 | 18 |
| | | Accuracy | 0,88 | | |
| | | Recall | 0,93 | | |
| | | F-Measure | 0,91 | | |
