## Supplementary_Excel_file_S4 Annotator_vs_OntoC for "OntoContext, a new python package for gene contextualization based on the annotation of biomedical texts"

| Num file | Reference Biotext Annotation | NCBO Annotator Annotation | OntoContext Annotation | VP_Annot | FP_Annot | FN_Annot | VP_Onto | FP_Onto | FN_Onto |
|---|---|---|---|---|---|---|---|---|---|
| file1 | | | | | | | | | |
| file2 | | | | | | | | | |
| file3 | GESTATIONAL DIABETES | GESTATIONAL DIABETES | GESTATIONAL DIABETES | 1 | | | 1 | | |
| file4 | AUTOSOMAL TRISOMIES | DOWN SYNDROME | DOWN SYNDROME | 1 | | 1 | 1 | | 1 |
| | BRADYCARDIA | PATAU SYNDROME | HYDROCEPHALUS | 1 | | 1 | | 1 | 1 |
| | CYTOGENETIC ABERRATIONS | SPINA BIFIDA | HYDRONEPHROSIS | 1 | | 1 | 1 | | 1 |
| | DOWN SYNDROME | | MICROCEPHALY | | 1 | | 1 | | |
| | DUODENAL ATRESIA | | PATAU SYNDROME | | | 1 | 1 | | 1 |
| | EDWARD SYNDROME | | SPINA BIFIDA | | | 1 | 1 | | 1 |
| | HYDRONEPHROSIS | | SYNDROME | | | 1 | | 1 | |
| | MICROCEPHALY | | TRISOMY 13 | | | 1 | 1 | | |
| | NONIMMUNE HYDROPS FETALIS | | TRISOMY 18 | | | 1 | 1 | | 1 |
| | PATAU SYNDROME | | | | | | | | |
| | PYELECTASIS | | | | | 1 | | | 1 |
| | SPINA BIFIDA | | | | | | | | |
| | TRISOMIES | | | | | 1 | | | 1 |
| | TRISOMY 13 | | | | | 1 | | | |
| | TRISOMY 18 | | | | | 1 | | | |
| | TRISOMY 21 | | | | | 1 | | | 1 |
| file5 | PRE-ECLAMPSIA | ECLAMPSIA | PRE-ECLAMPSIA | 1 | | | 1 | | |
| | PREECLAMPSIA | PRE-ECLAMPSIA | PREECLAMPSIA | 1 | | | 1 | | |
| file6 | | | | | | | | | |
| file7 | HEMIPLEGIA | HEMIPLEGIA | HEMIPLEGIA | 1 | | | 1 | | |
| | ASCITES | | | | | 1 | | | 1 |
| | MASSIVE FETOMATERNAL HEMORRHAGE | | | | | 1 | | | 1 |
| | HYPOTONIA | | | | | 1 | | | 1 |
| | HEPATOSPLENOMEGALY | | | | | 1 | | | 1 |
| file8 | OMPHALOCELE | OMPHALOCELE | OMPHALOCELE | 1 | | | 1 | | |
| | EXSTROPHY OF THE BLADDER | IMPERFORATE ANUS | IMPERFORATE ANUS | | 1 | 1 | | 1 | 1 |
| | MENINGOMYELOCELES | | | | | 1 | | | 1 |
| | OMPHALOCELE-EXSTROPHY-IMPERFORATE ANUS-SPINAL DEFECTS (OEIS COMPLEX) | | | | | 1 | | | 1 |
| | OEIS COMPLEX | | | | | 1 | | | 1 |
| | SPINAL DEFECTS | | | | | 1 | | | 1 |
| file9 | NONIMMUNE HYDROPS FETALIS | PERICARDIAL EFFUSION | PERICARDIAL EFFUSION | 1 | | 1 | 1 | | 1 |
| | HEMATOMA | | | | | 1 | | | 1 |
| | ASCITES | | | | | 1 | | | 1 |
| | LUCENCIES | | | | | 1 | | | 1 |
| | HYDROTHORAX | | | | | 1 | | | 1 |
| | PERICARDIAL EFFUSION | | | | | | | | |
| file10 | CHRONIC PROGRESSIVE EXTERNAL OPH | CHRONIC PROGRESSIVE EXTERNAL OPHTHAL | SYNDROME | 1 | | | | 1 | |
| | MITOCHONDRIAL DISEASES | OPHTHALMOPLEGIA | PROGRESSIVE EXTERNAL OPHTHALMOPLEGIA | | 1 | 1 | | 1 | |
| | HYPERTENSION | SYNDROME | CHRONIC PROGRESSIVE EXTERNAL OPHTHAL | | 1 | | 1 | | |
| | | HYPERTENSION | MITOCHONDRIAL DISEASES | 1 | | | 1 | | |

| | Annotator | OntoContext |
|---|---|---|
| Accuracy | 0,57 | 0,60 |
| Recall | 0,21 | 0,30 |
| F-Measure | 0,31 | 0,40 |

| | | | | | | | | | |
|---|---|---|---|---|---|---|---|---|---|
| | | | HYPERTENSION | | | | 1 | | |
| file11 | ALZHEIMER DISEASE (AD) | ALZHEIMER'S DISEASE | ALZHEIMER DISEASE | 1 | | | 1 | | |
| | | | NEUROLOGICAL DISORDERS | | | | | 1 | |
| file12 | OVARIAN CANCER | OVARIAN CANCER | OVARIAN CANCER | 1 | | | 1 | | |
| | BREAST CANCER | BREAST CANCER | BREAST CANCER | 1 | | | 1 | | |
| | CANCER | CANCER | CANCER | 1 | | | 1 | | |
| file13 | BREAST CANCER | BREAST CANCER | BREAST CANCER | 1 | | | 1 | | |
| | CANCERS | CANCER | CANCER | | 1 | 1 | | 1 | |
| | COLON CANCER | COLON CANCER | CANCERS | 1 | | | 1 | | |
| | PROSTATE CANCER | PROSTATE CANCER | COLON CANCER | 1 | | | 1 | | |
| | PROSTATE CANCERS | | HEREDITARY PROSTATE CANCER | | | 1 | | 1 | |
| | TUMORS | | PROSTATE CANCER | | | 1 | 1 | | 1 |
| | | | PROSTATE CANCERS | | | | 1 | | |
| file14 | MULTIPLE SCLEROSIS (MS) | MULTIPLE SCLEROSIS | MULTIPLE SCLEROSIS | 1 | | | 1 | | |
| file15 | SARCOMAS | SARCOMA | SARCOMA | 1 | | | | 1 | |
| | CANCERS | CANCER | CANCERS | 1 | | 1 | 1 | | |
| | CANCER | | PRIMARY CANCERS | 1 | | | | 1 | |
| | RETINOBLASTOMA | | CANCER | | | 1 | 1 | | |
| | | | SARCOMAS | | | | 1 | | |
| file16 | PREMATURE RUPTURE OF THE MEMBR | CHORIOMNIONITIS | CHORIOAMNIONITIS | | 1 | 1 | | 1 | 1 |
| | PRETERM PROM | | | | | 1 | | | 1 |
| file17 | | | | | | | | | |
| file18 | PARVOVIRUS INFECTION | PERICARDIAL EFFUSIONS | PERICARDIAL EFFUSION | 1 | | 1 | 1 | | 1 |
| | PERICARDIAL EFFUSIONS | | PERICARDIAL EFFUSIONS | | | | 1 | | |
| | PARVOVIRUS | | | | | 1 | | | 1 |
| | HYDROPS FETALIS | | | | | 1 | | | 1 |
| | PARVOVIRUS B19 INFECTION | | | | | 1 | | | 1 |
| | PERICARDIAL EFFUSION | | | | | 1 | | | |
| file19 | DEAFNESS | | | | | 1 | | | 1 |
| file20 | | | | | | | | | |
| file21 | SEIZURES | attention deficit hyperactivity disorder | EPILEPSY | | 1 | 1 | 1 | | 1 |
| | EPILEPSY | ADHD | ATTENTION DEFICIT HYPERACTIVIT | 1 | | | | 1 | |
| | ADHD | EPILEPSY | ADHD | 1 | | | 1 | | |
| file22 | | | | | | | | | |
| file23 | | | | | | | | | |
| file24 | | | | | | | | | |
| file25 | | | | | | | | | |

| | | | | | | | | | |
|---|---|---|---|---|---|---|---|---|---|
| file26 | ISCHAEMIC STROKE | | | | | 1 | | | 1 |
| | MYOCARDIAL INFARCTION | | | | | 1 | | | 1 |
| | CARDIOVASCULAR MORBIDITY AND MORTALITY AFTER MYOCARDIAL INFARCTION | | | | | 1 | | | 1 |
| file27 | CARDIAC ISCHAEMIC DEATH | CORONARY HEART DISEASE | HEART DISEASE | | 1 | 1 | | 1 | 1 |
| | CHLAMYDIA PNEUMONIAE | HEART DISEASE | CORONARY HEART DISEASE | 1 | | 1 | 1 | | 1 |
| | CORONARY HEART DISEASE | | | | | | | | |
| | CORONARY-ARTERY DISEASE | | | | | 1 | | | 1 |
| | ISCHAEMIA | | | | | 1 | | | 1 |
| | MYOCARDIAL INFARCTION | | | | | 1 | | | 1 |
| | UNSTABLE ANGINA | | | | | 1 | | | 1 |
| file28 | STAGE IB AND IIA CERVICAL CARCINOM | CARCINOMA | CARCINOMA | | 1 | 1 | | 1 | 1 |
| | CERVICAL CARCINOMA | | | | | 1 | | | 1 |
| file29 | | | | | | | | | |
| file30 | | | | | | | | | |
| file31 | | | | | | | | | |
| file32 | | | | | | | | | |
| file33 | | | | | | | | | |
| file34 | | | LEECHES | | | | | 1 | |
| file35 | ADENOCARCINOMA | ADENOCARCINOMA | ADENOCARCINOMA | 1 | | | 1 | | |
| | CANCERS | CANCER | CANCER | | 1 | 1 | | 1 | |
| | COLON CANCER | COLON CANCER | CANCERS | 1 | | | 1 | | |
| | COLON TUMORS | | COLON CANCER | | | 1 | 1 | | 1 |
| | FIBROSIS | | | | | 1 | | | 1 |
| | LYMPHATIC INVASION | | | | | 1 | | | 1 |
| | NECROSIS | | | | | 1 | | | 1 |
| | NUCLEAR ATYPIA | | | | | 1 | | | 1 |
| | TUMORS | | | | | 1 | | | 1 |
| file36 | GLYCOGEN STORAGE DISEASE TYPE II ( | GLYCOGEN STORAGE DISEASE II | GLYCOGEN STORAGE DISEASE TYPI | 1 | | | 1 | | |
| | | GLYCOGEN STORAGE DISEASE | GLYCOGEN STORAGE DISEASE | | 1 | | | 1 | |
| file37 | AUTONOMIC DYSFUNCTION | PARKINSON'S DISEASE | MULTIPLE SCLEROSIS | 1 | | 1 | 1 | | 1 |
| | CEREBRAL PALSY | CEREBRAL PALSY | PARKINSON'S DISEASE | 1 | | | 1 | | |
| | MULTIPLE SCLEROSIS | MULTIPLE SCLEROSIS | CEREBRAL PALSY | 1 | | | 1 | | |
| | NEUROLOGIC DISEASES | | | | | 1 | | | 1 |
| | PARKINSON'S DISEASE | | | | | | | | |
| file38 | | | | | | | | | |
| file39 | MYOFASCIAL TRIGGER POINT PAIN | | Group B | | | 1 | | 1 | 1 |
| | MYOFASCIAL TRIGGER POINTS OF UPPER TRAPEZIUS MUSCLE | | | | | 1 | | | 1 |
| | MYOFASCIAL TRIGGER POINTS | | | | | 1 | | | 1 |

| | | | | | | | | | |
|---|---|---|---|---|---|---|---|---|---|
| file40 | OSTEOPOROSIS | OSTEOPOROSIS | OSTEOPOROSIS | 1 | | | 1 | | |
| file41 | C TRACHOMATIS INFECTION | | | | | 1 | | | 1 |
| | CHLAMYDIA TRACHOMATIS | | | | | 1 | | | 1 |
| | INFECTION | | | | | 1 | | | 1 |
| | C TRACHOMATIS | | | | | 1 | | | 1 |
| file42 | | SYNDROME | SYNDROME | | 1 | | | 1 | |
| file43 | OBESITY | HYPERTENSION | PRIMARY PULMONARY HYPERTENSION | | 1 | 1 | | 1 | 1 |
| | | TOXIC ENCEPHALOPATHY | HYPERTENSION | | 1 | | | 1 | |
| | | PRIMARY PULMONARY HYPERTENSION | NEUROTOXICITY | | 1 | | | 1 | |
| file44 | ESSENTIAL HYPERTENSION | ESSENTIAL HYPERTENSION | HYPERTENSION | 1 | | | | 1 | |
| | | HYPERTENSION | ESSENTIAL HYPERTENSION | | 1 | | 1 | | |
| file45 | | | | | | | | | |
| file46 | INFECTION WITH ECHINOCOCCUS MULTILOCULARIS | | | | | 1 | | | 1 |
| file47 | CHRONIC MYELOID LEUKEMIA (CML) | LEUKEMIA | LEUKEMIA | | 1 | 1 | | 1 | 1 |
| file48 | HUMAN AMNESIC SYNDROME | SYNDROME | SYNDROME | | 1 | | | 1 | |
| file49 | | ALZHEIMER'S DISEASE | AMNESIA | | 1 | | | | 1 |
| | | AMNESTIC DISORDER | | | 1 | | | | |
| file50 | | | | | | | | | |
| file51 | | | | | | | | | |
| file52 | | | | | | | | | |
| file53 | OSTEOPOROSIS | OSTEOPOROSIS | OSTEOPOROSIS | 1 | | | 1 | | |
| file54 | | | | | | | | | |
| file55 | | | | | | | | | |
| file56 | TUMOURS | cancer | CANCERS | 1 | | 1 | 1 | | 1 |
| | CANCERS | | CANCER | | | 1 | 1 | | |
| | COLONIC TUMOURS | | | | | 1 | | | 1 |
| | CANCER | | | | | | | | |
| file57 | | ataxia telangiectasia | TELANGIECTASIA | | 1 | | | 1 | |
| | | telangiectasia | ATAXIA TELANGIECTASIA | | 1 | | | 1 | |
| file58 | | | | | | | | | |

| | | | | | | | | | |
|---|---|---|---|---|---|---|---|---|---|
| file59 | SEVERE FOOT INFECTIONS IN DIABETIC | cellulitis | CELLULITIS | | 1 | 1 | | 1 | 1 |
| | FOOT INFECTION IN DIABETIC PATIENTS | | | | | 1 | | | 1 |
| | FOOT INFECTIONS | | | | | 1 | | | 1 |
| | DIABETES | | | | | 1 | | | 1 |
| file60 | ORONARY-ARTERY THROMB | | | | | 1 | | | 1 |
| file61 | CARDIOVASCULAR SYMPTOMS | HYPERTENSION | HEART DISEASE | | 1 | 1 | | 1 | 1 |
| | HEART MURMUR | | HYPERTENSION | | | 1 | | 1 | 1 |
| | VALVE DISEASE | | | | | 1 | | | 1 |
| | PULMONARY HYPERTENSION | | | | | 1 | | | 1 |
| file62 | TRANSFER LESIONS | | | | | 1 | | | 1 |
| | HALLUX SUBLUXATION | | | | | 1 | | | 1 |
| | HEUMATOID ARTHRITI | | | | | 1 | | | 1 |
| file63 | STROKES | | | | | 1 | | | 1 |
| | CEREBROVASCULAR ACCIDENT | | | | | 1 | | | 1 |
| | CEREBROVASCULAR ACCIDENTS | | | | | 1 | | | 1 |
| | STROKE | | | | | 1 | | | 1 |
| file64 | HIGH BLOOD PRESSURE | HIGH BLOOD PRESSURE | HIGH BLOOD PRESSURE | 1 | | | 1 | | |
| file65 | | | | | | | | | |
| file66 | EARLY PARKINSON'S DISEASE | PARKINSON'S DISEASE | PARKINSON'S DISEASE | 1 | | | 1 | | 1 |
| | PARKINSON'S DISEASE | | | | | 1 | | | |
| file67 | | | | | | | | | |
| file68 | | osteomalacia | osteomalacia | | 1 | | | 1 | |
| file70 | | | | | | | | | |
| file71 | OTITIS MEDIA | | OTITIS MEDIA | | | 1 | 1 | | |
| | PHARYNGITIS | | PHARYNGITIS | | | 1 | 1 | | |
| | QUINSY | | QUINSY | | | 1 | 1 | | |
| | SINUSITIS | | SINUSITIS | | | 1 | 1 | | |
| | SORE THROAT | | TONSILLITIS | | | 1 | 1 | | 1 |
| | TONSILLITIS | | | | | 1 | 1 | | |
| | UPPER RESPIRATORY CONDITIONS | | | | | 1 | | | 1 |
| file72 | POSTOPERATIVE PAIN RELIEF | | ANALGESIA | | | 1 | | 1 | 1 |
| file73 | | | | | | | | | |
| file74 | EXTRAMEDULLARY HEMATOPOIESIS | neutropenia | neutropenia | | 1 | 1 | | 1 | 1 |
| | SPLENOMEGALY | | | | | 1 | | | 1 |
| | CHRONIC NEUTROPENIA | | | | | 1 | | | 1 |
| file75 | | | | | 1 | | | | |

| | | | | | | | | | |
|---|---|---|---|---|---|---|---|---|---|
| file76 | CROHN'S DISEASE | ABDOMINAL TUBERCULOSIS | | 1 | | 1 | | | 1 |
| | ABDOMINAL TUBERCULOSIS | | | | | | | | 1 |
| file77 | FEMALE STRESS URINARY INCONTINENC | | | | | 1 | | | 1 |
| file78 | | | | | | | | | |
| file79 | HELICOBACTER PYLORI INFECTION | | HELICOBACTER PYLORI INFECTION | | | 1 | 1 | | |
| | CORPAL GASTRITIS | | | | | 1 | | | 1 |
| | H. PYLORI INFECTION | | | | | 1 | | | 1 |
| file80 | PREECLAMPSIA (PROTEINURIC HYPERTI | HYPERTENSION | HYPERTENSION | | 1 | 1 | | 1 | 1 |
| | PREECLAMPSIA | PREECLAMPSIA | PREECLAMPSIA | 1 | | | 1 | | |
| file81 | | | | | | | | | |
| file82 | | | | | | | | | |
| file83 | | | | | | | | | |
| file84 | SCHIZOPHRENIC ILLNESSES | | | | | 1 | | | 1 |
| | SCHIZOPHRENIC SYMPTOMS | | | | | 1 | | | 1 |
| file85 | CLOACAL ANOMALIES | | | | | 1 | | | 1 |
| file86 | AMYLOIDOSIS | HEART disease | HEART DISEASE | | 1 | 1 | | 1 | 1 |
| | CARDIAC AMYLOID HEART DISEASE | | | | | 1 | | | 1 |
| | CARDIAC AMYLOIDOSIS | | | | | 1 | | | 1 |
| | HEART FAILURE | | | | | 1 | | | 1 |
| | MYOCARDIAL AMYLOIDOSIS | | | | | 1 | | | 1 |
| | PROTEIN DEPOSITION DISEASES | | | | | 1 | | | 1 |
| file87 | DIABETIC RETINOPATHY | diabetic retinopathy | | 1 | | | | | 1 |
| | RETINOPATHY | | | | | 1 | | | 1 |
| | DIABETES | | | | | 1 | | | 1 |
| file88 | GUNSHOT WOUNDS TO THE ABDOMEN | | | | | 1 | | | 1 |
| | GUNSHOT WOUNDS | | | | | 1 | | | 1 |
| | BLUNT TRAUMA | | | | | 1 | | | 1 |
| | BOWEL PERFORATIONS | | | | | 1 | | | 1 |
| | STAB WOUNDS | | | | | 1 | | | 1 |
| | INTRA-ABDOMINAL HEMORRHAGE | | | | | 1 | | | 1 |
| | INTRA-ABDOMINAL INJURY | | | | | 1 | | | 1 |
| file89 | ABDOMINAL PAIN | DIARRHOEA | DIARRHOEA | 1 | | 1 | 1 | | 1 |
| | PARASITOSIS | | | | | 1 | | | 1 |
| | PARASITATION | | | | | 1 | | | 1 |
| | DIARRHOEA | | | | | | | | |
| | BLASTOCYSTIS HOMINIS PARASITATION | | | | | 1 | | | 1 |
| file90 | | CANCER | CANCER | | 1 | | | 1 | |

| | | | | | | | | | |
|---|---|---|---|---|---|---|---|---|---|
| file91 | | | | | | | | | |
| file92 | | | | | | | | | |
| file93 | LUNG CANCER | LUNG CANCER | LUNG CANCER | 1 | | | 1 | | |
| | | CANCER | CANCER | | 1 | | | 1 | |
| file94 | SYNCOPE | | | | | 1 | | | 1 |
| | BLEEDING | | | | | 1 | | | 1 |
| | MAJOR PULMONARY EMBOLISM | | | | | 1 | | | 1 |
| | PRIMARY THROMBOLYSIS | | | | | 1 | | | 1 |
| | HEMODYNAMIC COMPROMISE | | | | | 1 | | | 1 |
| | CEREBRAL BLEEDING | | | | | 1 | | | 1 |
| | CARDIOGENIC SHOCK | | | | | 1 | | | 1 |
| | ARTERIAL HYPOTENSION | | | | | 1 | | | 1 |
| | CONGESTIVE HEART FAILURE | | | | | 1 | | | 1 |
| | CHRONIC PULMONARY DISEASE | | | | | 1 | | | 1 |
| file95 | PLEURAL MESOTHELIOMA | | ADENOCARCINOMA | | | 1 | 1 | | |
| | COLONIC POLYPS | | MALIGNANT MESOTHELIOMA | | | 1 | | 1 | 1 |
| | MESOTHELIOMA | | | | | 1 | | | 1 |
| | METASTATIC MALIGNANT MESOTHELIOMA | | | | | 1 | | | 1 |
| | MALIGNANT PLEURAL MESOTHELIOMA | | | | | 1 | | | 1 |
| | ADENOCARCINOMA | | | | | 1 | | | 1 |
| file96 | PERILESIONAL EDEMA | APHASIA | DYSKINESIAS | | 1 | 1 | | 1 | 1 |
| | VENOUS INFARCTION | PARKINSON'S DISEASE | APHASIA | | 1 | 1 | | 1 | 1 |
| | EDEMA | | PARKINSON'S DISEASE | | | 1 | | 1 | 1 |
| | ADVANCED PARKINSON'S DISEASE | | | | | 1 | | | 1 |
| | HEMORRHAGIC COAGULATION NECROSIS | | | | | 1 | | | 1 |
| | PARKINSONIAN SYMPTOMS | | | | | 1 | | | 1 |
| | ISCHEMIC INFARCTION | | | | | 1 | | | 1 |
| | BROCA'S APHASIA | | | | | 1 | | | 1 |
| | ISCHEMIC INFARCTIONS | | | | | 1 | | | 1 |
| file97 | | | | | | | | | |
| file98 | SENSORINEURAL HEARING LOSS | sensorineural hearing loss | SENSORINEURAL HEARING LOSS | 1 | | | 1 | | |
| | HEARING IMPAIRMENT | | | | | 1 | | | 1 |
| | DEVELOPMENTAL PROBLEMS | | | | | 1 | | | 1 |
| | HEARING LOSS | | | | | 1 | | | 1 |
| file99 | | | | | | | | | |
| file100 | GENODERMATOSIS | | GENODERMATOSIS | | | 1 | 1 | | |
| | DARIER DISEASE | | | | | 1 | | | 1 |
| | | | | 44 | 33 | 161 | 61 | 41 | 144 |
| | | Accuracy | | 0,57 | | | 0,60 | | |
| | | Recall | | 0,21 | | | 0,30 | | |
| | | F-Measure | | 0,31 | | | 0,40 | | |
