## Supplementary_Figure_S1 for "OntoContext, a new python package for gene contextualization based on the annotation of biomedical texts"

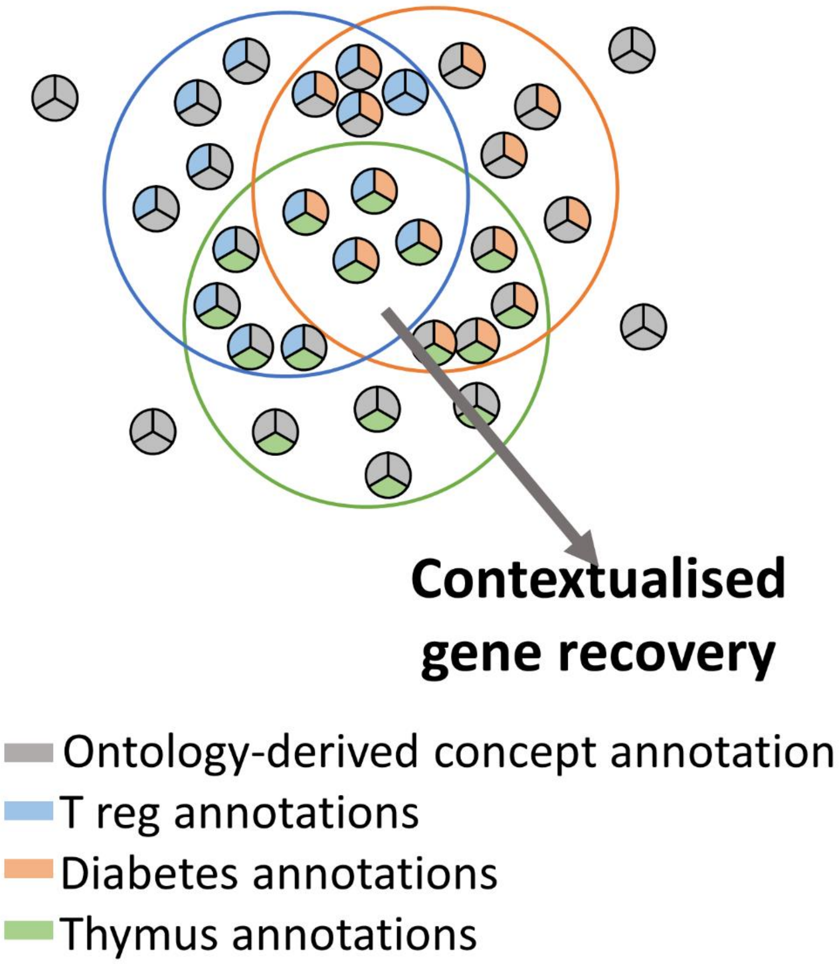


**Fig. S1. From contextualization of texts to gene list recovery.** Texts (symbolized by small circles) annotated with the three ontology-derived dictionaries (in grey) are represented here. In this example, texts annotated by regulatory T cell for Cell population Diabetes for Disease, thymus for Anatomical localization, are labelled in blue, orange or green, respectively, and intersected. The texts in the intersection are taken and the contextualized gene-derived list is recovered.
