## Supplementary_Figure_S2 for "OntoContext, a new python package for gene contextualization based on the annotation of biomedical texts"

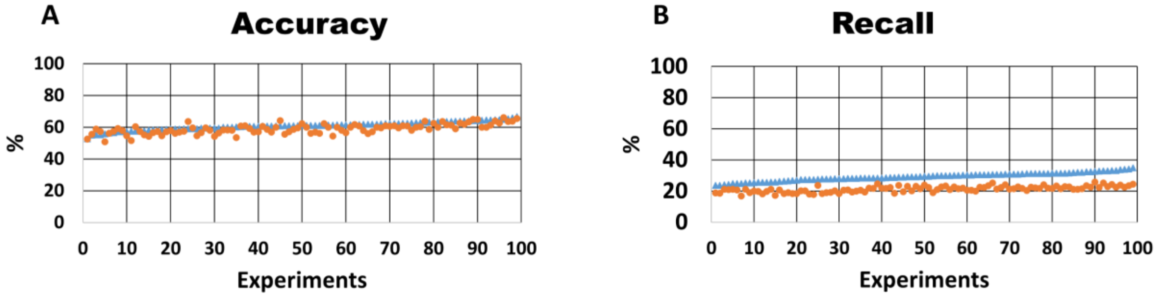


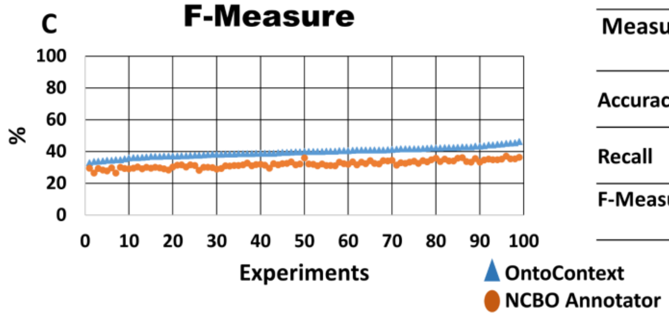


| Measure | Average | SD | Confidence interval | Pearson correlation | p-value (t-test) |
|---|---|---|---|---|---|
| Accuracy | 60.9 | 2.6 | 56-64 | 0.70 | 1.09E-12 |
| | 59.0 | 3.1 | 54-63 | | |
| Recall | 29.2 | 2.6 | 24-33 | 0.66 | 8.17E-63 |
| | 21.3 | 1.9 | 18-24 | | |
| F-Measure | 39.7 | 2.9 | 34-44 | 0.89 | 4.66E-76 |
| | 32.0 | 2.3 | 28-35 | | |

**Fig. S2. Comparing OntoContext and NCBO Annotator performances.** A k-fold derived validation test for the OntoContext vs. NCBO Annotator tool performance comparison using 2015 Human Disease Ontology-derived dictionary (the 75% sampling corpus in Figure 2). We considered the obtained NCBO Annotator and OntoContext annotations for 100 of the 141 abstracts of the BioText, and for each validation round, we drew 75% (75) of these 141 texts and assessed performances. Each pair of blue (OntoContext) and orange (NCBO Annotator) points represents one of the 99 measures of Accuracy (A), Recall (B) and F-Measure (C) in percentage. Results are sorted by increasing OntoContext performance values. The performances of both tools are represented on the same experience. The table summarizes the mean values, with the standard deviations (SD). The confidence interval was calculated using the 95% quantile. The correlation test is Pearson correlation between the OntoContext values and the NCBO annotator values. The p-value was calculated based on paired t-test.
