## Supplementary_Note_S1 for "OntoContext, a new python package for gene contextualization based on the annotation of biomedical texts"

**Note S1. Complexity calculation.** Principles of a classical exact matching algorithm (*Search exact matching (a1)*) vs. "*annot*" module (*annot (a2))* algorithm. The compared complexity performances of these two methods are shown.

| *Search exact matching (a1):* | *annot (a2):* |
|---|---|
| ***Lc** = List of **L** concepts*<br>***Sentence** = Sentence with length **Le***<br><br>*For i in L:*<br>*If i in Sentence:*<br>*list_sortiei.append(i)*<br><br>***Complexity***<br>$C_{a1} = L*(Le+1)$<br>$= L*Le+L$ | ***Lc** = List of **L** concepts*<br>***Dict_tag** = Dictionary of* $L_{dic}$ *concepts grouped by POS_Tag*<br>***Dict_tag[i]** = List of* $L_{dic[i]}$ *concepts Tagged by the same morphosyntactical **tag TS** = POS_Taged(sentence) with length* $L_{TS}$<br><br>*For i in Dict_tag.keys():*<br>*If i in TS:*<br>*If Dic_tag(i) in Sentece(POs_Tag):*<br>*list_sortiei.append(Dic_tag(i))*<br>***Complexity***<br>$C_{a2} = L_{dic}*(1+ L_{TS} +L_{Dict\_tag[i]})$<br>$= L_{dic}+ L_{TS}*L_{dic} + L_{Dict\_tag[i]}*L_{dic}$ |

Considering the Table S1

→$L \gg Ldic$ and $Le = LTs$

→$L * le \gg Ldic * LTs$

→$L * le + L \gg Ldic * LTs + Ldic$

→$Ca1 \gg Ca2 - (LDict_{tag[i]} * Ldic)$

→$Ca1 + (LDict_{tag[i]} * Ldic) \gg Ca2$

We know that: $LDict_{tag[i]} * Ldic \ll L * Le + L$ (see Table S1)

→$Ca1 \gg Ca2$
