## Supplementary_Table_S1 for "OntoContext, a new python package for gene contextualization based on the annotation of biomedical texts"

**Table S1. Sources and characteristics of the dictionaries used in this work.**

| Dictionary name | Type | Source downloaded on the 13/04/2015 | Terms[a] | Labels[b] |
|---|---|---|---|---|
| CRAFT | Expert | CRAFT corpus annotations, CRAFT corpus (http://bionlp-corpora.sourceforge.net/CRAFT/) | 407 | 114 |
| 2012 Cell Ontology | Ontology (cell type) | 2012 Cell Ontology, from CRAFT Corpus (http://bionlp-corpora.sourceforge.net/CRAFT/) | 2,628 | 345 |
| 2015 Cell Ontology | Ontology (cell type) | 2015 Cell Ontology, OBO Foundry / Basic ontology https://raw.githubusercontent.com/obophenotype/cell-ontology/master/cl-basic.obo) | 8,523 | 1,393 |
| BioText | Expert (pathologies) | BioText corpus (http://biotext.berkeley.edu/data/dis_treat_data.html) | 245 | 101 |
| 2015 Human Disease Ontology | Ontology (pathologies) | 2015 Human Disease Ontology, OBO Foundry / Basic ontology (http://www.berkeleybop.org/ontologies/doid.obo) | 22,957 | 3,802 |
| 2015 UBERON Ontology | Ontology (anatomy) | 2015 UBERON Ontology, OBO Foundry / Composite-vertebrate (http://berkeleybop.org/ontologies/uberon/composite-vertebrate.obo) | 4,691 | 589 |

[a] Number of concepts with their synonyms and plural form in the dictionary

[b] Number of labels obtained after the morphosyntactic labelling
