## Supplementary_Excel_file_legends for "OntoContext, a new python package for gene contextualization based on the annotation of biomedical texts"

**Supplementary excel file legend**

**Excel file S1. OntoContext performance on Cell population's annotation according to three dictionaries.** The 67 texts from the CRAFT 1.0 corpus are annotated with OntoContext. The annotation of each text is performed according to three dictionaries (see Table 1 and Figure 3) and results are given in separate excel tables. Using the CRAFT 1.0 as a reference, the occurrence of terms identified with OntoContext is classified as True Positive (TP) or False Positive (FP) and False Negative (FN). The OntoContext performance (Accuracy, Recall, F-measure) is calculated at the bottom of each table.

**Excel file S2. OntoContext and NCBO annotator performance for Cell population's annotations using the 2015 Cell Ontology-derived dictionary.** The CRAFT 1.0 Annotation is considered as reference for the NCBO Annotator Annotation and the OntoContext Annotation. "1" identify occurrence of terms as True Positive (TP) or False Positive (FP) and False Negative (FN). In this experience, we annotate only 35 texts from CRAFT 1.0 corpus. The performances presented here are calculated for all the 35 texts.

**Excel file S3. OntoContext performance on Disease's annotation according to two dictionaries.** The 141 texts from the BioText corpus are annotated with OntoContext. The annotation of each text is performed according to two dictionaries (see Table 2 and Figure S2) and results are given in separate excel tables. Using the BioText as a reference, the occurrence of terms identified with OntoContext is classified as True Positive (TP) or False Positive (FP) and False Negative (FN). The OntoContext performance (Accuracy, Recall, F-measure) is calculated at the bottom of each table.

**Excell file S4. OntoContext and NCBO Annotator performance for disease's annotations using the 2015 Human Disease Ontology-derived dictionary.** The BioText Annotation is considered as reference for the NCBO Annotator Annotation and the OntoContext Annotation. "1" identify occurrence of terms as True Positive (TP) or False Positive (FP) and False Negative (FN). In this experience, we annotate only 100 texts from BioText corpus. The performances presented here are calculated for all the 100 texts.
